# Ancient balanced polymorphism underlies long-standing adaptation for seasonal camouflage in the least weasel

**DOI:** 10.1101/2025.11.14.688436

**Authors:** Inês Miranda, Raquel Ruivo, Liliana Farelo, Marcela Alvarenga, Zbigniew Borowski, Morten Elmeros, Daniela C. Kalthoff, Juha Merilä, Jürg P. Müller, Laurent Schley, Franz Suchentrunk, Janne Sundell, Mónica Rodrigues, Margarida Santos-Reis, Carlos Rodríguez Fernandes, Karol Zub, Jeffrey M. Good, L. Scott Mills, L. Filipe C. Castro, Love Dalén, José Melo-Ferreira

## Abstract

Unraveling how adaptive traits originate and evolve is key to understanding the mechanisms shaping species’ diversity and their adaptive potential. Seasonal color molts, from summer-brown to winter-white, evolved in at least 21 mammals and birds to maintain camouflage in environments with seasonal snow, but the occurrence of winter-brown morphs reflects seemingly convergent local adaptation to distinct snow conditions. In the least weasel (*Mustela nivalis*), alternative winter morphs map to the pigmentation gene *MC1R*, but the evolutionary history and functional basis of this variation remain unknown. Using *in vitro* cellular assays, we show that winter-brown coats are caused by a derived protein-coding amino acid substitution that reduces MC1R affinity to its ligands, ASIP and α-MSH. Using targeted enrichment and sequencing, we find that this mutation arose *de novo* within the species, around one million years ago, and was maintained across the geographically structured populations generated during its evolution in Europe. Using simulations, we show that genetic drift cannot explain the long-term maintenance of this variant, which is likely driven by spatially varying selection acting on the phenotypic polymorphism, anchoring local adaptive responses. Our results underscore how long-standing adaptive variation can fuel recurrent adaptation to heterogeneous environments through time.

## INTRODUCTION

Phenotypic variation within species often results from adaptation to locally varying environmental conditions across distribution ranges. The study of the mechanisms underlying the evolution of adaptive traits in natural populations has gained increasing attention^1^, particularly to understand how this variation is originated^2–4^ and the relative roles of gene flow, selection, and drift in maintaining adaptive variation and potential through time^5,6^. Locally adapted morphs reflect the past adaptive evolution of species responding to environmental challenges. Understanding the genetic basis and evolutionary mechanisms that shaped current standing genetic variation within species, especially adaptive genetic variation, can be particularly important to assess the adaptive potential of these species in the face of environmental change^7,8^, as this information can be used to inform predictions of species resilience under new conditions^9–11^.

Seasonal coat color change, from summer-brown to winter-white coats, for camouflage against seasonal snow cover, emerged as a recurrent adaptive response to annual selective pressures in at least 18 mammal and three bird species across the Northern Hemisphere^12,13^. In these species, winter color variation across their range reflects adaptation to local snow conditions, with winter-brown morphs prevailing in regions with shorter or more ephemeral snow duration^13^. Importantly, this adaptive polymorphism may be a key ingredient for future adaptive responses^14–16^, as these species face compromised camouflage of winter-white morphs due to climate change-induced decreases in snow cover duration^17^.

Least weasels, *Mustela nivalis*, are small carnivores with a circumpolar distribution across the Northern Hemisphere. Both brown and white winter morphs exist in the species, with brown being predominant in some southern and coastal areas of the species’ distribution range, both in Eurasia and North America^13,18,19^. In Europe, winter coloration morph has been used as a criterion for subspecies classification^19^, but this division does not correspond to the species population structure and evolutionary history^20–22^. For consistency, we will here use the subspecies nomenclature to refer to the alternative coloration morphs from Europe: *M. n. nivalis* (hereafter *nivalis*) are winter-white, show a straight dorsoventral line in the summer-brown coat, and are generally smaller; *M. n. vulgaris* (hereafter *vulgaris*) are winter-brown, with a ragged dorsoventral line and distinctive brown cheek spots, and are generally larger^18,19^ (Fig. 1A). Crossing experiments between *nivalis* and *vulgaris* specimens^23^ and genomic analyses^22^ have shown that winter pelage color and dorsoventral line type segregate together, suggesting the same or tightly linked genetic bases. Whole-genome scans for winter color association based on museum specimens have shown that alternative color morphs in *M. nivalis* map to the genomic region of the melanocortin-1 receptor (*MC1R*) gene^22^, a well-known component of the melanogenesis pathway^24^. MC1R is a receptor on the membrane of melanocytes that is signaled by 1) the α-melanocyte stimulating hormone (α-MSH)^25^, which promotes receptor activity and leads to the production of eumelanin (brown to black pigment), or 2) the agouti-signaling protein (ASIP), which inhibits MC1R activity^26^, leading to the production of pheomelanin (red to yellow pigment) or inhibiting melanin production^27^. This previous work^22^ pinpointed a candidate amino acid substitution in the MC1R protein caused by two consecutive single-nucleotide polymorphisms (SNPs) in the same codon but could not establish it as the causal mutation. The *MC1R vulgaris* allele was found to be derived, with a signature of a recent selective sweep in the Swedish winter-brown morph, implying a post-glacial adaptive response to decreasing snow. Yet, such a signature was not evident in the Polish population, hinting at a potentially complex haplotype structure for the winter-brown variant^22^.

**Fig. 1.**
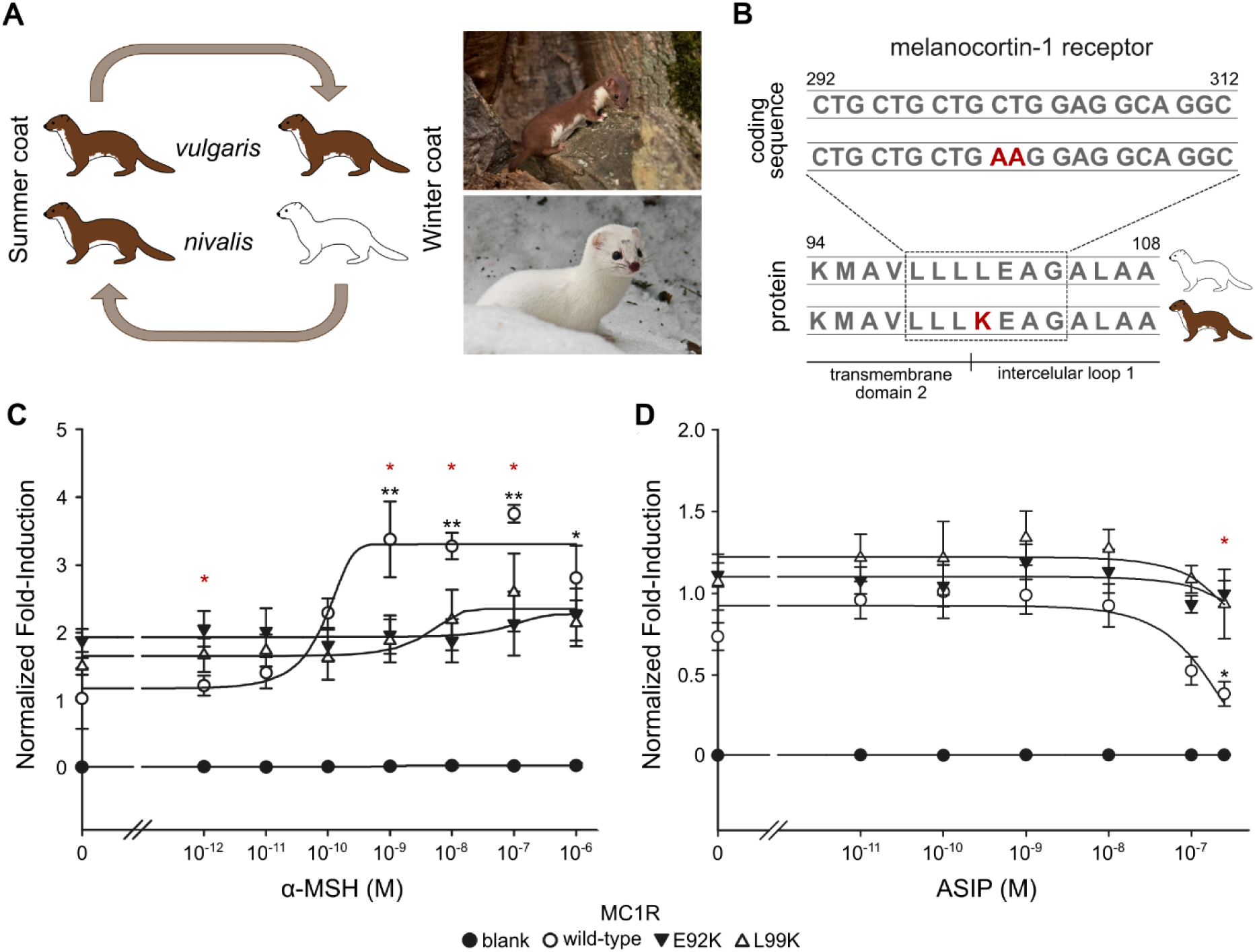
Seasonal color polymorphism and functional characterization of MC1R *vulgaris* allele. **(A)** *Left:* Seasonal coat color variation in *Mustela nivalis*, including alternative winter color morphs – *nivalis* (white) and *vulgaris* (brown). *Right*: *M. nivalis* specimens harboring a *vulgaris* coat (top) and a winter *nivalis* coat (bottom). Photos: Karol Zub. **(B)** Location of two consecutive variants in the *MC1R* coding sequence and respective leucine (L) to lysine (K) substitution in the resulting protein^22^. **(C-D)** *In vitro* MC1R receptor activity, as measured by luciferase activity in response to cAMP production, normalized by maximum forskolin activation, for three alternative MC1R constructs (wild-type mouse protein, constitutively-active E92K mouse mutant^28^, and L99K mouse mutant, homologous to *M. nivalis* substitution in panel B), in response to exposure to increasing concentrations of **(C)** α-MSH and **(D)** ASIP. Black asterisks denote minimum significant differences detected in receptor activity for a given construct, compared to exposure to lower ligand concentrations. Red asterisks denote minimum significant differences detected in receptor activity between the wild-type and mutant constructs exposed to the same ligand conditions. For full details on pairwise comparisons, see Supplementary Tables S2 to S4.

Here, we first functionally validate the candidate MC1R mutation and show that it reduces ligand affinity to the receptor, compatible with the resulting winter-brown pelage. Second, we investigate the evolutionary origin of the causal mutation, comparing genomic variation across 11 weasel species, including two other species showing brown-to-white seasonal color variation and polymorphism of the winter color. Third, we infer the history and haplotype structure of the *MC1R* region within the framework of the modeled population demographic history of the species. Finally, we simulate the evolution of the adaptive locus to understand the relative roles of genetic drift and selection in allowing the maintenance of the adaptive variants along the evolution of least weasels. Ultimately, our study provides a detailed view of the evolutionary mechanisms underlying the repeated evolution of a key locally adaptive trait and the population dynamics underlying the maintenance of adaptations through time.

## RESULTS

### MC1R mutation underlies reduced receptor affinity to ASIP and α-MSH

An amino acid variant in MC1R has been identified as a candidate for determining discrete alternative winter coloration in the least weasel, i.e., brown or white^22^. This mutation results in the substitution of a leucine (L) for a lysine (K) in residue 101 of the protein sequence (*nivalis* L101K *vulgaris*; Fig. 1B). To assess the functional impact of the amino acid mutation on ligand affinity, we conducted *in vitro* cellular assays, in which we transfected HEK 293-T cells with vectors containing different MC1R sequences and exposed them to increasing concentrations of α-MSH (N = 3 replicates) or ASIP (N = 6). We used well-established patterns of MC1R activity in mice as the base-model for our experiments, comparing receptor activity patterns among an L99K mutant, expressing a mutation homologous to the candidate *M. nivalis* L101K substitution (Fig. 1B), and the mice wild-type and E92K mutant MC1R (known to be constitutively active, increasing basal activity and severely reducing responsiveness to the agonist α-MSH)^28,29^.

We found that both the wild-type and the E92K mutant recovered the expected activity patterns based on previous descriptions^28,29^. Wild-type MC1R activity increased when exposed to increasing concentrations of α-MSH (non-linear regression (NLR), P < 0.001; Fig. 1C; Supplementary Table S1)^28^ but decreased in response to ASIP (NLR, P < 0.05; Fig. 1D; see also ref.^30^). The E92K MC1R mutant was mostly unresponsive to the varying concentrations of the ligands, consistent with its described constitutive activation^28,29^, even though small changes in activity are detected for concentrations equal to or above 10^-7^ M of α-MSH (NLR, P < 0.05; Supplementary Table S1; Fig. 1C; see also ref.^29^). In addition, basal receptor activity of the constitutive mutant was 1.5 to 1.8 times higher than that of the wild-type and about 50% of the maximum activity level of the wild-type receptor under α-MSH stimulation (Fig. 1C), following previous descriptions^28,29^. The L99K mutant showed increased activity relative to the wild-type across all ASIP concentrations, with higher ASIP concentrations trending toward decreasing receptor activity (but P > 0.05 for NLR; Supplementary Table S1; Fig. 1D). Exposure to α-MSH showed increased activity with increasing concentrations of the ligand (NLR, P < 0.05; Supplementary Table S1), but milder than the wild-type and with higher basal activity (Fig. 1C). Additionally, statistically significant differences in activity were found for the wild-type MC1R across ligand concentrations, for both α-MSH and ASIP (ANOVA and Student-Newman-Keuls test, P < 0.05; Supplementary Table S2; Fig. 1C-D) but not in the E92K and L99K constructs (ANOVA, P > 0.05; Supplementary Table S2). Conversely, significant activity differences were found between the wild-type receptor and both E92K and L99K mutants exposed to the same ligand concentration (ANOVA and Student-Newman-Keuls test, P < 0.05; Supplementary Tables S3 and S4) but not between E92K and L99K (ANOVA, P > 0.05). These results show that the L99K mutation alters MC1R function, increasing its basal activity and reducing its affinity to α-MSH and likely ASIP.

We also phenotyped least weasel museum skins representative of *nivalis* (N = 55) and *vulgaris* (N = 70) morphs, sampled both during summer and winter, to assess whether functional MC1R differences could impact the coat color expressed in phenotypically brown specimens. Using a standardized digital photography approach to collect color data along the dorsum of the specimens, we found that the brown color of winter *vulgaris* specimens is significantly lighter than that of the summer *vulgaris* and *nivalis* pelage (Kruskal-Wallis test, P < 0.05) but no significant differences were found between the summer pelages (Kruskal-Wallis test, P > 0.05; Supplementary Text S1; Supplementary Figs. S1 and S2). This may result from changes in MC1R function in *vulgaris* specimens, affecting their winter phenotype.

### The derived winter-brown allele is specific to least weasels

We next addressed the origin of winter color polymorphism in the least weasel in the context of other mustelid species. We analyzed targeted enrichment and re-sequencing data (∼5 Mb, including the association region surrounding *MC1R* and anonymous genome-wide regions; see Methods for capture design details) from *M. nivalis* and 10 other mustelid species from the *Mustela* and *Neogale* genera, including two other species with seasonal coat color change and polymorphism of the winter color, the stoat (*M. erminea*) and the long-tailed weasel (*N. frenata*)^18,31^. Our dataset comprised 12 *M. nivalis* of both color morphs (capture and sequencing output reported below; Supplementary Data S1 and S2) and 27 specimens of other mustelid species, of which 24 were kept for analyses (average of 37X target coverage, 297-fold enrichment, and 30% off-bait sequencing; Supplementary Data S3 and S4). The dataset was complemented with publicly available whole-genome data for three *M. nigripes* and two outgroup specimens, one wolverine (*Gulo gulo*) and one European otter (*Lutra lutra*) (Supplementary Data S3)^32,33^.

Using a coalescent-based approach^34^, we reconstructed a highly supported species tree (based on 993 genome-wide capture fragments; ∼1.91 Mb) that recovered the phylogenetic relationships among species previously described from multilocus approaches^35,36^ (Fig. 2A; Supplementary Fig. S3). In comparison, a maximum-likelihood phylogeny centered on *MC1R* (40 kb region, based on homozygosity decay within *M. nivalis*; see below) resulted in a similar topological arrangement of the tree, including the monophyly of *M. nivalis* and the broad species relationships, with no allele sharing across species (Fig. 2B; Supplementary Fig. S4). Genotyping of the *MC1R* causal mutation across all individuals further showed that the *vulgaris MC1R* variant is derived and occurs exclusively in *M. nivalis*, being absent from other sampled mustelid species, including those with similar seasonal coat color change phenotype variation (Fig. 2C; Supplementary Table S5). Using GEVA^37^, we further estimated the origin of the winter-brown *vulgaris* allele at 0.96 million years ago (Mya), based on the time to the most recent common ancestor (TMRCA) of *M. nivalis* sequences harboring alternative alleles (using an extended least weasel dataset, reported below). This estimate post-dates the divergence of the species to its closest relative, *M. altaica*, estimated from 2 to 2.87 Mya^35,36,38^. Overall, our results support a species-specific origin of the *M. nivalis* winter-brown *vulgaris* variant. Interestingly, genotyping of North American least weasels (N = 5, all from the United States; Supplementary Data S1 and S2) showed that the European winter-brown mutation appears absent, including from specimens expressing winter-brown morphs (N = 3; Supplementary Table S6).

**Fig. 2.**
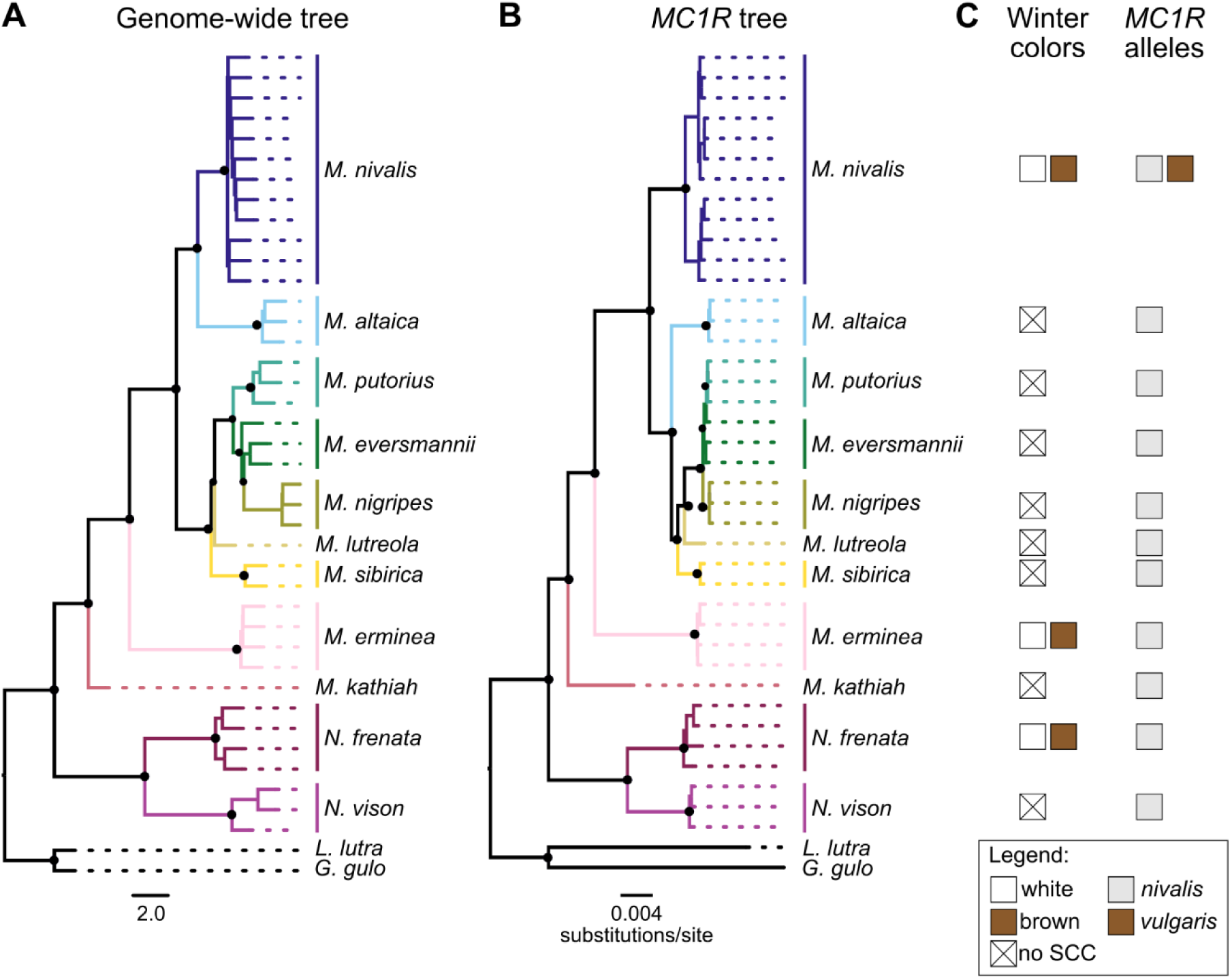
The origin of *MC1R vulgaris* alleles. **(A)** Coalescent genome-wide species tree estimated based on anonymous capture targets (993 genomic fragments, ∼1.91 Mb sequence length), for 11 *Mustela* and *Neogale* species. *Lutra lutra* and *Gulo gulo* were used as outgroups. **(B)** Maximum likelihood *MC1R* gene tree estimated based on a 40 kb region centered on *MC1R* SNPs (see Fig. 4C), for the same sample set. All major nodes in both trees have (A) posterior probability = 1.0 or (B) bootstrap values = 1.0, respectively (black dots). See Supplementary Figs. S3 and S4 for detailed tree support values. **(C)** Winter phenotypes and *M. nivalis MC1R* alleles identified in each species. SCC = seasonal coat color.

### Two anciently diverged lineages within European least weasel populations

We investigated the evolution of the winter-brown *vulgaris* variant in European *M. nivalis*, expanding the capture and sequencing to a total of 247 European *M. nivalis* (*nivalis*, N = 103; *vulgaris*, N = 88; and uncertain phenotype, N = 56), originally sampled in nine countries, including three winter morph transition zones in Sweden, Poland, and the Alps (Fig. 3A; Supplementary Data S1). All samples originated from specimens stored at natural history museums or research collections, and capture efficiency and specificity were high (average 37X target coverage, 328-fold enrichment, and 16% off-bait sequencing; Supplementary Data S2), resulting in 242 European *M. nivalis* kept for analyses.

**Fig. 3.**
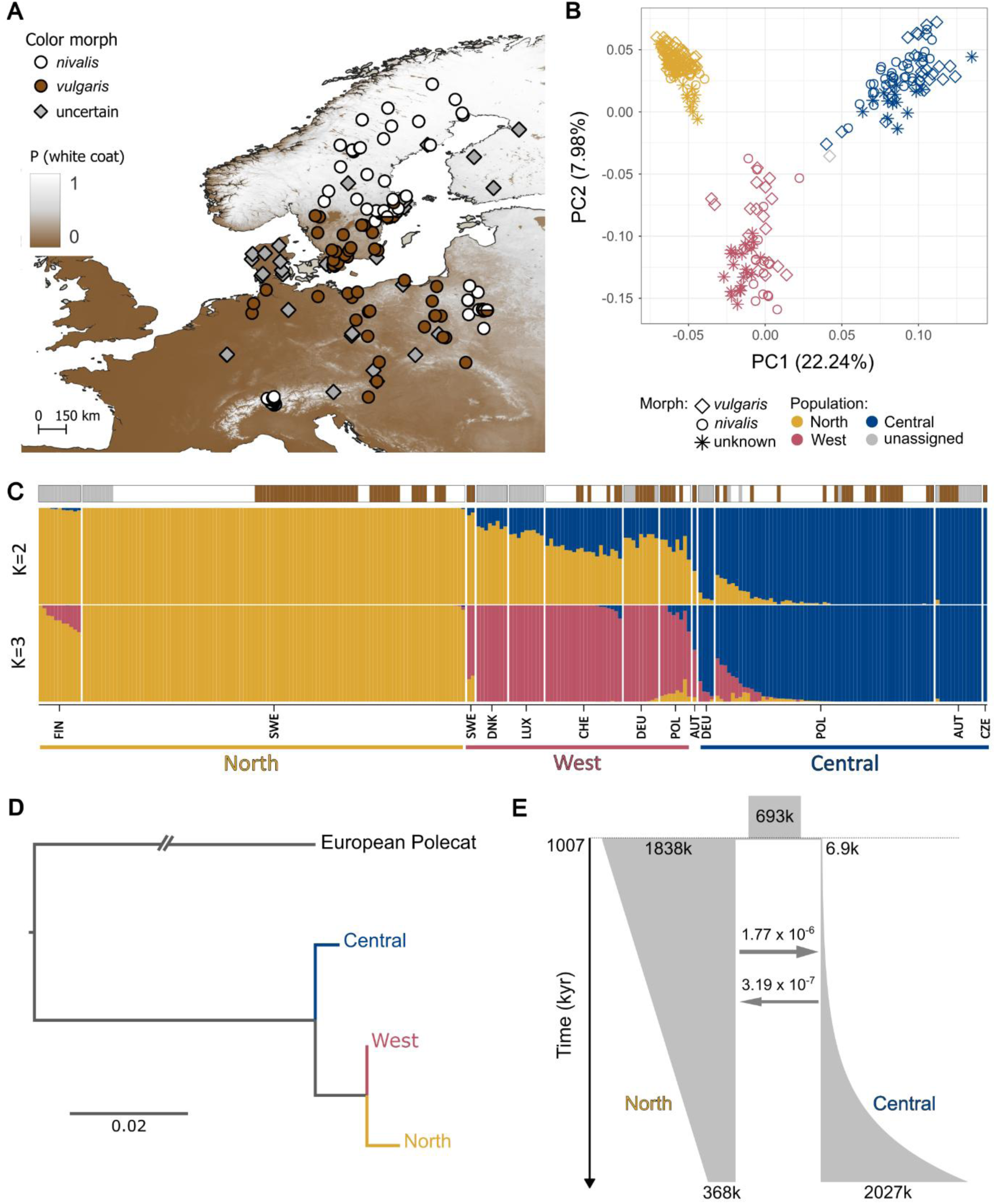
Population structure and demographic history of European least weasels. **(A)** Sampling localities of European least weasels used in this study; a single symbol indicates overlapping localities. Circle colors indicate sampled coloration morphs, based on *a priori* phenotypic information, and diamonds denote *a priori* uncertain phenotype. Sampling is overlaid on the clinal distribution of winter colors, shown as the probability of winter-white coats across the European distribution range of the species (adapted from ref.^13^, with data available in ref.^41^). The species range was obtained from the IUCN Red List (https://www.iucnredlist.org/). **(B)** Principal component analysis of genetic variation, based on 8,140 genome-wide SNPs, inferred from anonymous capture regions, for the complete dataset of analyzed European least weasels (N = 242). Individuals are colored according to their population assignment, based on the best number of ancestral clusters (K = 3; panel C). **(C)** Admixture proportions inferred considering two and three genetic clusters, based on the same SNP set as (B). Samples’ countries of origin and inferred genetic lineages are identified below the plot. Visually phenotyped color morphs (*nivalis*, *vulgaris*, uncertain) are shown above the plot, following the color code from (A). **(D)** Population tree estimated based on allele frequencies inferred from 8,140 SNPs. The European polecat (*M. putorius*) was used as an outgroup. Original outputs, including associated residuals, are shown in Supplementary Fig. S6. **(E)** Joint demographic history of North and Central European least weasel lineages. Point estimates of the best-fitting model for effective population sizes (thousands of individuals, k), migration rates (proportion of migrants/generation), and divergence time (thousands of years, kyr) are shown in the plot. Confidence intervals for the estimates are available in Supplementary Table S7.

We started by exploring the population history of European least weasels, which provided the background context to understand the evolution of the adaptive polymorphism. Using a dataset of 8,140 unlinked genome-wide SNPs (N = 242 individuals; Fig. 3A; Supplementary Data S2), a principal component analysis (PCA) showed that samples segregate along three main axes of differentiation (Fig. 3B), which was further supported by an ADMIXTURE analysis, where K = 3 was identified as the number of genetic clusters that best explains genetic variation within the dataset (Fig. 3C; Supplementary Fig. S5A). These clusters broadly correspond to the geographic distribution of samples (Supplementary Fig. S5B) and are hereafter designated as North, West, and Central lineages. The reconstruction of a population tree under the TreeMix algorithm suggested that the North and West lineages are more closely related (Fig. 3D; Supplementary Fig. S6A), a result also supported by a neighbor-joining tree based on genetic distances (Supplementary Fig. S6B) and the ADMIXTURE results for K = 2 (Fig. 3C).

We next modeled the parameters of the demographic history of least weasels in Europe used ∂a∂i^39^ in conjunction with GADMA^40^, focusing on the North and Central lineages, which were consistently identified as the most divergent across analyses (Fig. 3; Supplementary Fig. S6; insufficient dataset power hindered the estimation of a joint demographic model for the three identified lineages). The best demographic scenario showed a good fit to the empirical data (Supplementary Fig. S7) and indicated an estimated divergence time of ∼1 million years, followed by population divergence with asymmetrical migration (Fig. 3E). In addition, our best-fit model inferred a strong population bottleneck in the Central lineage, followed by exponential growth, whereas the North lineage was inferred to undergo population decline since the lineages split (Fig 3E; Supplementary Table S7).

### Ancient winter-brown variant originated before the least weasel diversification in Europe

We then focused on analyzing variation along the *MC1R* genomic region. We tested whether we could find other genetic variants along this region that could be associated with the different dorsoventral lines that also characterize the *nivalis* and *vulgaris* morphs, which previous work suggested to be determined by the same locus that causes the winter coloration differences or a tightly linked one^22,23^. We compared case-control association tests in two distinct sample groups, based on the available phenotypic information, focusing either on winter color (white, N = 66, or brown, N = 40; 156,141 SNPs) or on the dorsoventral line type (straight, N = 35, or ragged, N = 86; 204,777 SNPs; Supplementary Data S2). Our results showed that *MC1R* remained the strongest association peak in both datasets, with the same functionally impactful mutations (two consecutive SNPs causing one amino acid change; Fig. 1) being the top associated variants in both analyses (Supplementary Fig. S8A-B,D-E). These were the only two SNPs along the *MC1R* region that showed a perfect association with alternative phenotypes in both traits (Supplementary Fig. S8C,F; Supplementary Table S6), which are inherited as dominant winter-brown color and ragged dorsoventral line^23^. Additional genotype-to-phenotype association tests, using a combined dataset based on both phenotyped coloration traits (*nivalis,* N = 101, or *vulgaris*, N = 87; 218,609 SNPs; Supplementary Data S2), resulted in an increased association signal in *MC1R* (Fig. 4A; Supplementary Fig. S9) while retaining the perfect association between phenotype and genotype at the causal SNPs (Fig. 4B; Supplementary Table S6), supporting that *MC1R* variation underlies both traits.

**Fig. 4.**
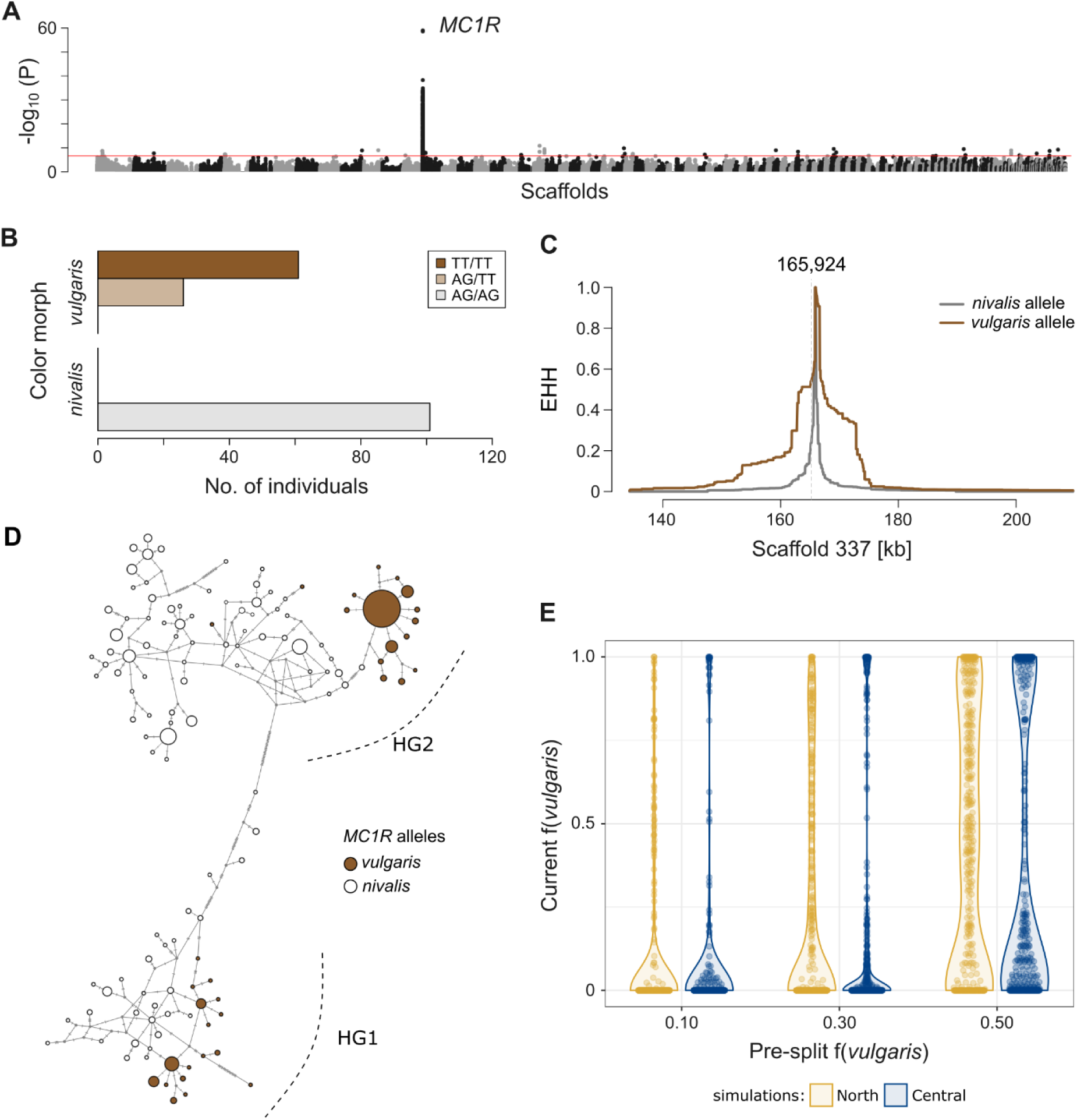
The evolution and maintenance of *MC1R* variation. **(A)** Case-control genotype-to-phenotype mapping of color morph (*nivalis*, N = 101, or *vulgaris*, N = 88), based on 218,609 SNPs across all captured genomic regions. The red line indicates the Bonferroni-corrected threshold of P = 0.05. The top associations are the two consecutive *MC1R* SNPs (Fig. 1B). **(B)** Genotypes per color morph at the causal *MC1R* SNPs, for all specimens used in (A). **(C)** Extended haplotype homozygosity estimates based on statistically phased haplotypes, centered on the first *MC1R* SNP (position 165,924; scaffold 337). **(D)** Neighbor-joining haplotype network based on 484 phased haplotypes for a 40 kb *MC1R*-centred region. Each circle is one haplotype, with size proportional to the number of individuals that share it. Dashes in each branch denote mutated sites. Circle colors denote alternative alleles at *MC1R* SNPs in each haplotype. **(E)** Forward-in-time simulations of *vulgaris* phenotype frequency, f(*vulgaris*), under neutral genetic drift, following the demographic history inferred between North and Central lineages (Fig. 3E). Models seeded a dominant neutral mutation (selection coefficient, s = 0) at varying frequencies one generation before the population split, to control for distinct possibilities of f(*vulgaris*) at the time of lineage divergence. A total of 400 replicates were simulated per model.

To explore genetic diversity surrounding *MC1R* SNPs, we estimated extended haplotype homozygosity (EHH)^42^ based on statistically phased haplotypes and found evidence for quick homozygosity decay with increasing distance to the causal variants (EHH < 0.1 at a maximum of 12,478 bp from focal SNPs; Fig. 4C), suggesting little linkage disequilibrium surrounding these SNPs. Reconstructing the haplotype structure using phased genotypes across all analyzed European specimens (N = 484 haplotypes) showed the occurrence of two very distinct *MC1R* haplotypic groups (Fig. 4D), which are geographically structured: haplogroup 1 (HG1) occurs almost exclusively in specimens from the Central European least weasel lineage, whereas haplogroup 2 (HG2) is the most predominant in both North and West lineages, but also occurs in Central specimens (Supplementary Figs. S10 and S11). The derived winter-brown *MC1R* allele was associated with both divergent haplogroups (Fig. 4D; Supplementary Fig. S10), which explains the rapid decay of linkage disequilibrium in the flanking region of the causal SNPs (Fig. 4C). D_XY_ estimates between specimens harboring the same *MC1R* alleles, but with distinct flanking variation, showed that the divergence of alternative haplogroups falls within the 5th to 95th percentile of genome-wide D_XY_ estimates (Supplementary Fig. S12), showing that the *MC1R* local haplotype structure coincides with the wide genomic signal. Also, dating the divergence of the alternative *MC1R* haplogroups under a Bayesian inference approach^43^ resulted in an estimated TMRCA of 0.95 Mya (95% highest posterior density (HPD) of 0.63 to 1.26 Mya; Supplementary Fig. S13; Supplementary Table S8), which is consistent with the age of the most basal split of least weasel genomic lineages inferred in our demographic models (Fig. 3E; Supplementary Table S7) and with the estimated age of the winter-brown variant (see above). These results suggest a scenario where the *vulgaris MC1R* allele arose at the onset of the least weasel’s diversification in Europe.

### MC1R diversity in least weasels is maintained by long-term selection

Our data showed that, in each of the North and Central lineages, both winter-brown and winter-white morphs are segregating (Supplementary Figs. S10 and S11), and previous modeling of winter phenotype distribution predicted intermediate probabilities of occurrence of each morph in our sampling areas^13^ (see Supplementary Table S9). We found that this polymorphism occurs within populations with no appreciable genetic structure (Fig. 3B,C), whereas winter color phenotypes are geographically structured (Fig. 3A), correlating with variation in winter snow coverage^13^. Using forward-in-time simulations^44^ based on our inferred demographic model (Fig. 3E), we tested whether the history of these populations could explain the maintenance of the polymorphism, segregating as a neutral locus, in the two genetic lineages simultaneously. Our simulations rejected the maintenance of the *vulgaris* winter-brown morph at intermediate frequencies (0.20 to 0.80) in both diverging genetic lineages, considering varying frequencies of the *vulgaris* phenotype at the onset of lineage divergence: 10% (P = 0.0075), 30% (P = 0.0300), and 50% (P = 0.0425) (Fig. 4E). Our results thus showed that genetic drift alone would lead to the loss of the color polymorphism and cannot explain its maintenance along the ∼1 million years of evolution of European least weasels, suggesting that selection is at play.

We also found that specimens harboring specific variation at *MC1R* can have significant local reductions of genetic variation when compared to the genome-wide level (P < 0.05, Kruskal–Wallis test; Supplementary Fig. S14), which would be compatible with the effects of recent selection acting on locally favorable genotypes. As an example, we found significant reductions of both nucleotide diversity (π) and Tajima’s D for specimens harboring HG2 with *vulgaris* alleles (P < 0.001; Supplementary Fig. S14), and we found evidence for a local *MC1R* selective sweep in the same set of specimens (i.e., North and West lineage winter-brown specimens; Supplementary Fig. S15), with an estimated selection coefficient of s = 0.003 (Supplementary Fig. S16).

## DISCUSSION

Our work dissects the functional basis, origin, and evolutionary history of winter coat color variation in the least weasel, a polymorphism anchoring local adaptive responses for camouflage that is repeated across mammal species with the ability to change their coat color annually, matching seasonal snow. While this polymorphism had been shown to map to the genomic region of *MC1R*^22^, a key pigmentation gene^24^, our functional assays linked a single dominant derived amino acid variant in MC1R to the determination of brown winter coats. We found that the *vulgaris* MC1R allele alters the receptor function by increasing its basal activity and reducing affinity to its ligands, α-MSH and ASIP, with activity patterns more like those of the constitutive E92K receptor than the wild-type MC1R (Fig. 1). Previous work has linked seasonal molt to white to up-regulation of the *ASIP* gene^14^, which shows an expression peak during the autumn molt^45,46^. Our results are thus compatible with the hypothesis that the reduced ligand affinity of the *vulgaris* MC1R variant leads to an insufficient response to the expected pulse of *ASIP* expression, preventing color change from brown to white. These findings establish a clear link between a single amino acid mutation and its functional impact in a wild species. Mutations in *MC1R* have been widely linked to static color variation in numerous species^47–49^, but our results expand this knowledge by showing how the same gene can modulate polymorphism in a seasonally flexible trait, acting as a switch that turns off the capacity for winter whitening. These results also contrast with findings on hare species (*Lepus* spp.), in which seasonal camouflage polymorphism has been linked to *cis*-regulatory variation in the *ASIP* gene^14–16^, resulting in decreased *ASIP* expression in non-white winter morphs^15^. Both mechanisms can hamper winter whitening caused by an *ASIP* expression peak during the molt^45,46^, either increasing the threshold for antagonist-induced MC1R response or turning the *ASIP* expression pulse insufficient to induce color molting. We also found that *vulgaris* individuals carrying the winter-brown mutation have a lighter brown pelage in winter than either morph in summer (Supplementary Figs. S1 and S2). This result is consistent with previous qualitative descriptions of lighter winter fur based on observations of least weasel pelts, to which the contribution of a higher density of grey underfur has been hypothesized^50^. A minor response to the ASIP pulse during the autumn molt of *vulgaris*, but insufficient to cause white coats, could also contribute to lightening fur color by lightening guard hair color. Interestingly, our fine-scale inspection of patterns of association along the *MC1R* region further suggested that the same amino acid variation is perfectly associated with the ragged or straight lines that separate the dorsal and ventral pelage of least weasels (Supplementary Fig. S8). Such spatial segregation in the body of mammals is established during embryonic development^51^, and our results suggest that MC1R plays a role in such differentiation. The mechanism under which an amino acid mutation in MC1R could underlie these differences remains speculative but might involve alternative binding affinities to ASIP isoforms expressed during embryogenesis^52^, causing differential migration or survival of melanocyte precursors^53^.

Finding a simple genetic architecture of a trait with important adaptive value in natural populations allowed dissecting its origin and the evolutionary processes underlying its maintenance as polymorphic in the gene pool of the species. Our phylogenetic analyses showed that the derived winter brown variant is not shared across other weasel species but arose through a species-specific *de novo* mutation in least weasels (Fig. 2). Again, this differs from other systems with winter whitening in which patterns of shared standing variation across species were found, resulting either from recent secondary introgression of alleles causing non-white winter morphs^14,15^ or from the combination of recent introgression and the maintenance of shared ancestral variation along the diversification of the genus^16^. While faster adaptation is expected to be based on standing adaptive variation^2,3^, some conditions could have facilitated this process. First, the dominance of the least weasel winter-brown *vulgaris* allele^22,23^ causes phenotypic expression even in heterozygosity, facilitating a faster initial increase in frequency if advantageous^54^. Second, the *vulgaris* morph has been hypothesized to have arisen in southeastern Europe during a period of warmer climate^19^, and a strong selective advantage would be expected for a winter-brown morph in a snowless environment^13^. Overall, our results show that genetic polymorphisms leading to convergent adaptive responses to variations in snow cover repeatedly evolved across species through distinct genetic bases and following alternative evolutionary routes, but still recruiting the same physiological pathway, the melanin production pathway. These results align with the prediction that convergent phenotype evolution can be constrained at the molecular level, favoring the recruitment of the same pathways, genes, or even mutations to underlie similar phenotypes^55,56^. For instance, empirical reports, both among closely related species^57,58^ and more distantly related groups^59^, often show the recruitment of the same gene underscoring similar adaptive responses. Additionally, our results hint that different genetic bases underlie winter-brown phenotypes across weasel species, as the *vulgaris* winter-brown variant was not found in stoats and long-tailed weasels, the two other mustelids with winter coat color polymorphism (Fig. 2B, Supplementary Fig. S4), and within least weasels, as it is also absent from the few analyzed North American specimens expressing a winter-brown phenotype (Supplementary Table S6). This implies that the repeated evolution of alternative winter phenotypes might extend to the intraspecific level, as also suggested for snowshoe hares (*Lepus americanus*)^60^.

Having established that the derived winter-brown variant arose *de novo* in European least weasels, we investigated the age of the mutation and dated its origin to ∼1 Mya, at the onset of the species diversification, which compares to the age of trans-species adaptive polymorphisms in the radiation of Darwin finches^61,62^. We found that least weasels in Europe form three geographic-explicit divergent lineages (Fig. 3; Supplementary Fig. S6), and that, in all three, the two winter phenotypes are co-segregating (Supplementary Figs. S10 and S11), which agrees with prior models on the probability of distribution of the two winter phenotypes^13^. Two lineages, North and West, were found to be closely related, and the third, Central, was more divergent (Fig. 3D; Supplementary Fig. S6), which is broadly consistent with the distribution of the two major mitochondrial DNA (mtDNA) lineages identified in previous phylogeographic work^20,21^. For example, our results suggest that samples from Poland fall within different genomic clusters (Fig. 3C), which agrees with previous reports of Poland as a suture zone for distinct mtDNA lineages^63^. We estimated that the first lineage split occurred about ∼1 Mya (Fig. 3E), and the 95% confidence interval of our estimate (936 to 1,079 thousand years ago (kya); Supplementary Table S7) slightly overlapped more recent estimates of *M. nivalis* TMRCA based on an mtDNA dataset (560 to 940 kya coalescent interval)^21^. When we focused our analyses on the *MC1R* region, we found a clear structure in two main haplogroups (Fig. 4D), which overlapped the North/West and Central genomic lineages (Supplementary Figs. S10 and S11), with average divergence estimates similar to the remainder of the genome (Supplementary Fig. S12). This result suggests that the structure of *MC1R* conforms to the genome-wide norm and must reflect the underlying demographic history of the populations. The occurrence of the winter-brown mutation in both haplotypic backgrounds (Fig. 4D) with global rapid linkage disequilibrium decay in the flanking region suggests that it arose before population divergence, ∼1 Mya, and was maintained along the least weasel diversification in Europe.

The long-term maintenance of an adaptive variant segregating in natural populations results from the relative strength of different evolutionary forces, particularly genetic drift and selection. Using simulations anchored on the inferred population history, we have shown that genetic drift is insufficient to explain the maintenance of alternative winter phenotypes segregating in two diverging genetic lineages (Fig. 4E), which suggests that selection is likely involved. Given the clinal distribution of alternative winter phenotypes across Europe in response to snow cover variation^13^, spatially varying selective pressures favoring alternative morphs, according to local snow conditions, may have promoted the long-term maintenance of this adaptive polymorphism in least weasels. These results agree with data from other systems in which geographically structured adaptive variation was linked to distinct environmental conditions^64–66^, providing a clear example of how environmental clines can maintain variation with fitness-relevant phenotypic effects as a locally adapted balanced polymorphism^1^. Strikingly, we also find signatures of a selective sweep acting on *vulgaris* variation in the northern parts of the species range (Supplementary Figs. S15 and S16), recovering a previously identified sweep in the Swedish population^22^. This signal might result from post-glacial selection for the winter-brown allele during the northwards species expansion^19,22^, suggesting that temporal variation of environmental conditions, for example, due to glacial and interglacial climatic periods, could further alter phenotypic distribution in response to changing snow cover extent, favoring distinct phenotypes at a given location through time. Collectively, these data indicate that, at a regional scale, selection driven by environmental changes in snow conditions left genomic signals of selective sweeps acting on the locally favored variation. However, at a global scale, spatially and potentially temporally heterogeneous camouflage pressures promoted the balanced maintenance of alternative *MC1R* variants underlying distinct coloration morphs in least weasels.

Overall, our work shows that a single dominant amino acid change in MC1R, which arose at the onset of the diversification of least weasels in Europe, about 1 Mya, functionally affects the determination of alternative winter phenotypes in the species, which evolved in response to local winter snow conditions. Anthropogenic climate change is challenging the adaptive value of seasonal camouflage^17^, endangering the persistence of winter-white populations^67^. The occurrence of standing genetic variation for alternative winter color morphs might thus be fundamental to anchor rapid evolutionary responses to reduced snow cover conditions^16^. Understanding the genetic basis and evolutionary mechanisms underlying key adaptive polymorphisms can provide crucial information to inform eco-evolutionary modeling of climate change responses across species^8^, which in turn can inform species conservation and evolutionary rescue.

## METHODS

### *In vitro* functional assays of MC1R *vulgaris* variant

The impact of the candidate MC1R amino acid substitution^22^ on the receptor function was tested using *in vitro* cellular assays. HEK 293-T cells (ECECC 12022001) were maintained in Dulbecco’s Modified Eagle’s Medium (DMEM, PAN-Biotech, Aidenbach, Bayern, Germany) with phenol red, and supplemented with 10% fetal bovine serum (PAN-Biotech, Aidenbach, Bayern, Germany) and 1% penicillin/streptomycin (PAN-Biotech, Aidenbach, Bayern, Germany) at 37 °C in a humidified chamber containing 5% CO_2_. Before transfection, cells were seeded in 6-well plates at a density of 1 × 10^6^ cells/well. On the following day, cells were transfected using 8 μL Lipofectamine 2000 (Invitrogen, Thermo Fisher Scientific, Waltham, MA, USA), 700 ng of the reporter vector pGL4.29[luc2P/*CRE*/Hygro] (Promega, Madison, WI, USA), 20 ng of the constitutive control reporter pRL-TK[Rluc/*TK*] (Promega, Madison, WI, USA) serving as internal control of transfection efficiency, and 200 ng of pCDNA 3.1 (+) MC1R constructs, in Opti-MEM reduced serum medium (Gibco, Thermo Fisher Scientific, Waltham, MA, USA)^68^. Mouse MC1R wild-type (Genbank accession: NM_008559.2) and mutant sequences (E92K, L99K) were synthesized and cloned into pCDNA 3.1 (+) vector by Genscript (Hong Kong). Transfection procedures were carried out according to the manufacturer’s recommendations. After 5 h of incubation, cells were washed with phosphate-buffered saline and incubated in DMEM medium without phenol red, supplemented with 10% charcoal-stripped fetal bovine serum (PAN-Biotech, Aidenbach, Bayern, Germany) and 1% penicillin/streptomycin (PAN-Biotech, Aidenbach, Bayern, Germany). After 24 h, cells were harvested and plated in 96-well poly-D-lysine coated plates at a density of 1 × 10^5^ cells/well. At 48 h post-transfection, cells were stimulated using different concentrations of mouse α-MSH (Abcam, Netherlands) or mouse ASIP (R&D Systems, Bio-Techne, Ireland) in DMEM medium containing 0.1 mg/ml BSA and 0.1 mM Isobutylmethylxanthine (BPS Bioscience, San Diego, USA) for 6 h at 37 °C in a humidified chamber containing 5% CO_2_. Cells were also stimulated by 10 mM forskolin to normalize across test constructs^29^. Each condition was assayed in duplicate. After 6 h of stimulation, cells were washed and gently lysed for 15 min at 37 °C and 90 rpm, using 50 μL/well of Passive Lysis Buffer (Promega, Madison, WI, USA). Firefly luciferase (pGL4.29) and Renilla luciferase (pRL-TK) luminescent activities were assessed using the Dual luciferase assay system according to the manufacturer’s instructions (Promega, Madison, WI, USA) and measured with a Synergy HT Multi-Mode Microplate Reader (Biotek, Agilent Technologies, Santa Clara, CA, USA). Experiments were repeated at least three times. Transactivation assay data were obtained through the ratio of Firefly luciferase and Renilla luciferase activity and then normalized with respect to forskolin stimulation. Obtained transactivation data are available in a Figshare repository^69^. Response curves were fitted by nonlinear regression using SigmaPlot 11 software (https://grafiti.com/sigmaplot-detail/; see Supplementary Table S1).

Statistically significant differences between transactivation data were tested using one-way analysis of variance (ANOVA), as implemented in SigmaPlot 11. ANOVAs were done per construct (comparing receptor activity of the same MC1R construct at different ligand concentrations) and per condition (comparing receptor activity among constructs for each ligand concentration). Data normality was checked using the Shapiro-Wilk test (P-value > 0.05) and data variance was tested using the Levene test (P-value > 0.05). ANOVAs were conducted when, at minimum, equal variances were found, given that the test is generally robust to deviations from data normality^70^. If significant differences were found, post-hoc pairwise multiple comparisons were conducted using the Student-Newman-Keuls test (Supplementary Tables S2 to S4).

### Phenotyping of museum skins

To characterize potential phenotypic differences among specimens harboring alternative coloration morphs, a total of 134 least weasel skins from the collection of the Swedish Museum of Natural History (NRM) were photographed for phenotypic analyses (Supplementary Data S1). Specimens were classified into four phenotypic groups, according to their coloration morph (*nivalis* or *vulgaris*) and collection season (winter or summer). Classification into each season was based on the following criteria: i) summer specimen, if sampled between May and September, and ii) winter specimen, if sampled between December and February, i.e., the height of winter color^50^. Classification into each coloration morph was based on i) the distinct dorsoventral line patterns (straight or ragged) for summer specimens, and ii) the dorsal pelage color (white or brown) for winter specimens^18,19^. Specimens with suggestions of ongoing molts were not included in the dataset. Digital photographs of the specimens were taken using a Nikon D5200 camera, with an 18-140 mm lens. Photographs were taken with a fixed lens-to-specimen distance and lighting conditions. An X-Rite Mini ColorChecker Passport (X-Rite Inc.) was included in every picture, and images were shot in RAW file format. Original RAW files were imported to Adobe Lightroom Classic v.9.1.0.10 for pre-processing and standardization. First, photos were converted to DNG format with Adobe Lightroom, and a custom camera profile was generated for each specimen using the ColorChecker Camera Calibration software (X-Rite Inc.). After, using Adobe Lightroom, photos were normalized following the approach of ref.^71^: i) the generated camera DNG profiles were applied to each photo; ii) the white balance of each image was manually set using the first grey square of the Mini ColorChecker passport (square no. 20); and iii) the exposure was calibrated until the Red, Green, and Blue (RGB color space) values of the reference grey square matched the expected 200:200:200 (± 5).

Standardized digital images were then imported to the ImageJ software (v.1.53o)^72^ for extraction of RGB color values. For each specimen, one dorsal photograph was analyzed. Four sampling points were defined as circular regions of interest (ROI), with a diameter of 150 pixels and aligned by the body midline: head (H), upper dorsum (UD), middle dorsum (MD), lower dorsum (LD) (Supplementary Fig. S1A). Individual Red, Green, and Blue (RGB color model) values were extracted for each sampling point, using the *Measure* option. Any ROI falling on top of damaged skin (for example, with missing hair or where skin patches were sampled for DNA) was set as missing data. A total of 125 specimens with no missing data (30 winter *nivalis*, 25 summer *nivalis*, 32 winter *vulgaris*, and 38 summer *vulgaris*) were retained for further analysis. Before analyses, RGB values were first converted to Hue, Saturation, and Value (HSV) with the *rgb2hsv* R function (R v.4.1.0; ref.^73^), a color space that is more intuitive and easier to interpret by lay users^74^. Principal component analyses (PCAs) were conducted on the transformed measurements with the *prcomp* R function to characterize major axes of color variation. Biplots of scaled and centralized measures were used to identify the variables driving differentiation among phenotypic groups. The normality of the data was assessed per variable, using a Shapiro-Wilk test. Given small deviations from normality found in three variables (H hue, MD saturation, LD saturation), we conservatively used non-parametric Kruskal-Wallis tests to compare variation in each phenotype measure between pairs of brown groups: i) summer *vulgaris* vs. summer *nivalis*, ii) summer *vulgaris* vs. winter *vulgaris*, and iii) summer *nivalis* vs. winter *vulgaris*. Either “morph” or “season” was used as a predictor variable in the analysis, against each phenotypic variable (used as dependent variables). Statistical tests were conducted with R package *rstatix* (v.0.7.2)^75^, and significance was assessed at Bonferroni-corrected P-values < 0.05. Collected RGB data, transformed HSV values, and code for analyses are available on a Figshare repository^69^.

### Custom targeted-enrichment design

We designed a custom capture assay of 5 Mb of genomic sequence, based on the draft genome assembly of *Mustela nivalis* (GenBank accession: GCA_019141155.1)^22^. Regions to include in the capture were selected following three criteria: i) previously identified regions relevant for the determination of the winter polymorphism (0.5 Mb), ii) randomly selected putative intronic regions (1,000 fragments of 2 kb), and iii) randomly selected putatively intergenic regions (7,143 regions of 300 to 350 bp). For the first criterion, one region of 500 kb surrounding the *MC1R* gene was selected, based on previous results that associate genomic variation at this pigmentation gene with alternative winter morphs in the least weasel^22^. The length of the region to capture was determined based on F_ST_ decay along the association region (as originally described^22^). For the second and third criteria, regions were selected to allow the inference of genetic variation representative of the genomic background, considering genome scaffolds > 50 kb in length. To define intergenic and intronic DNA in the genome of *Mustela nivalis*, for which annotation is currently lacking, we set up a pipeline to allow the inference of genic and non-genic regions along the assembly, based on mRNA and protein homology data from other carnivore species (see Supplementary Text S2 for a detailed description of the methods used for region definition). The final list of targets to include in the capture design was randomly selected from the resulting putative intergenic and intronic datasets, assuring both i) a minimum distance of 250 kb from the trait-associated region including the *MC1R* gene, and ii) a minimum distance of 25 kb between all target regions, to reduce the effect of linkage disequilibrium in downstream analyses. Custom probes were designed by Roche Sequencing Solutions (Madison, WI, USA), using the KAPA HyperExplore MAX 5Mb design approach. Final capture probes covered 97.70% of originally targeted genomic bases, corresponding to ∼4.87 Mb of the genome, but had a total coverage estimate (including adjacent probe regions) of up to 99.36% of targets (∼4.96 Mb of genome sequence). Details on the final capture design, including statistics, original targets, and probe-covered regions, can be found in the Figshare repository^69^.

### Genetic sampling strategy and data collection

A total of 259 *Mustela nivalis* individuals, representative of both European (N = 247) and North American populations (N = 12) of least weasels (*Mustela nivalis*), were included in this work for genetic analyses (Supplementary Data S1). Sampled specimens were representative of three European transition zones (Sweden, N = 103; Poland N = 66; the Alps, Switzerland, N = 20), as well as regions outside polymorphic areas (*nivalis*: Finland, N = 11; and *vulgaris*: Denmark, N = 9; Germany, N = 13; Luxembourg, N = 9, Austria, N = 15, Czech Republic, N = 1). North American samples were originally collected in North Dakota (winter-white area; N = 5) and Ohio (transition zone; N = 7). Dry skin patches were collected from 177 specimens, from the collections of the Swedish Museum of Natural History (NRM; N = 95), the Zoological Research Museum Alexander Koenig (ZFMK, N = 7), the American Museum of Natural History (AMNH; N = 12), and the research collection of the Mammal Research Institute of the Polish Academy of Sciences (MRI; N = 63). Tissue samples (N = 62) were obtained from the Bündner Naturmuseum (BNM; N = 26), the research collection of the Centre for Ecology, Evolution and Environmental Changes (CE3C; N = 26), the Senckenberg Museum of Natural History Goerlitz (SMNG; N = 4), and the Franz Suchentrunk research collection (FSRC; N = 6). DNA extracts (N = 20) were obtained for samples of the CE3C research collection (N = 12) and the NRM collection (N = 8). Scoring of the coloration morph of European individuals relied on distinct criteria (i.e., winter color and/or dorsoventral line type)^18,19,23^ according to the available metadata. For skins sampled at NRM and ZFMK, and BNM tissue samples, individual photographs of whole skins were available, which allowed visually classifying individuals as i) *nivalis* morph, if having a straight dorsoventral line (summer coat) or white fur (winter coat), and ii) *vulgaris* morph, if having a ragged dorsoventral line (summer and winter coats)^18,23^. For MRI skins, direct observations of the color morph were available within collection metadata and were validated against the geographic distribution of each morph^13,76^. For DNA extracts and tissue samples of the CE3C, SMNG, FSRC, and NRM, we could not directly confirm phenotypes during the development of this work. We thus *a priori* classified specimens as “uncertain morph” and later scored individuals as *nivalis* and *vulgaris*, according to their genotype at causal *MC1R* SNPs (confirmed as the only perfect association for winter color and dorsoventral line phenotypes; see *Genotype-to-phenotype mapping* section). For all genotype-based morph assignments, the inferred morph was further validated against sampling locality and the expected phenotype occurrence, given previous morph distribution models^13^. North American least weasels exhibiting both winter phenotypes are described^31^, but subspecies differ from the European ones^18,19,31^. Thus, we scored specimens for winter color and dorsoventral line type based on photographs of whole skins, with collection dates between December and February, but did not attribute a color morph (Supplementary Data S1). We split the least weasel specimens into three sample subsets, based on location and *a priori* phenotype confirmation (Supplementary Data S1): subset 1 (Europe and confirmed phenotype), subset 2 (Europe and uncertain phenotype), and subset 3 (not from Europe), which were used in distinct analyses.

A total of 30 individuals representative of 12 other Mustelidae species (*Mustela altaica*, N = 3; *M. erminea*, N = 4; *M. eversmannii*, N = 3; *M. kathiah*, N = 1; *M. lutreola*, N = 1; *M. nigripes*, N = 3, *M. nudipes*, N = 1; *M. putorious*, N = 3; *M. sibirica*, N = 2; *M. strigidorsa*, N = 2; *Neogale frenata,* N = 4; *N. vison*, N = 3) were added to our dataset for phylogenetic analyses, together with one European otter (*Lutra lutra*)^33^ and one wolverine (*Gulo gulo*)^32^ used as outgroups (Supplementary Data S3). Dry skin samples for 27 specimens were obtained from the museum collections of NRM (N = 19), ZFMK (N = 3), Museum of Comparative Zoology at Harvard University (MCZ; N = 2), Smithsonian National Museum of Natural History (USNM; N = 2), and National Museum of Ireland (NMI; N = 1). Raw sequencing data was obtained for five individuals from publicly available data (Supplementary Data S3)^32,33^. Seasonal color change and winter color polymorphism occur in two of these species, *M. erminea* and *N. frenata*^13,18,31^; therefore, specimens from these species with winter collection dates (see above) were scored for winter color based on whole skin photographs.

### Laboratory work and sequencing

Dry skin samples collected at AMNH, MCZ, MRI, NMI, NRM, USNM, and ZFMK were extracted in rooms appropriate for historical DNA. *M. nivalis* and *N. frenata s*kin patches were extracted following ref.^77^. For all remaining skin samples, the protocol was modified by using the extraction buffer described in ref.^78^. Ethanol-preserved tissue samples (BNM, CE3C, FSRC, and SMNG collections) were extracted using the EasySpin Genomic DNA Tissue Kit (Citomed, Lisbon, Portugal), following the manufacturer’s instructions.

Illumina sequencing double-indexed libraries were prepared for 259 *Mustela nivalis* and 27 additional Mustelidae individuals. Distinct protocols were used for lower-quality (dry skin patches) and higher-quality (ethanol-preserved) samples (see Supplementary Data S1), both adapted from ref.^79^, with the modifications from ref.^80^ and some additional changes (see Supplementary Text S3 for details on modifications to the library protocol). Libraries were checked for quality and insert size estimation using a 2200 TapeStation (Agilent Technologies) and quantified by Qubit Fluorometer (Invitrogen) or qPCR. Lower-quality libraries were subjected to a test sequencing run, to estimate endogenous DNA content. Libraries were then pooled for capture assays in 14 pools (16 to 32 samples each). Samples were pooled based on the DNA insert size, endogenous DNA quantity, and library preparation protocol to minimize capture biases. Pools were organized by species for *M. nivalis*, *M. erminea*, and *N. frenata*. The remaining Mustelidae samples were pooled together in a single assay, though this strategy was expected to result in distinct capture success, according to species divergence from the *M. nivalis* genome used for probe design^81^. Capture assays were performed following the manufacturer’s instructions in the KAPA HyperCap Workflow v.3.0 manual (Roche Sequencing Solutions), using 10 cycles in the post-capture PCR. Capture pools were quantified by qPCR and sequenced in Illumina NovaSeq 6000 platforms at Novogene (Cambridge, UK), using a 2 x 150 bp sequencing strategy.

### Genomic data processing

Raw sequencing data quality was assessed with FastQC (v0.11.7)^82^ and adapter sequences trimmed with Trimmomatic (v.0.39)^83^, using MINLEN:25, TRAILING:15, SLIDINGWINDOW:5:15. Overlapping pairs were merged with PEAR (v.0.9.11)^84^ if minimum read overlap ≥ 10 bp and minimum merged read size ≥ 30 bp. Reads were mapped to the draft *M. nivalis* reference genome (NCBI accession: GCA_019141155.1) using BWA-MEM (v.0.7.17)^85^ with default parameters (using scaffolds > 50 kb, i.e., those used for capture probe design). Read group information was assigned using the *AddorReplaceReadGroups* option of Picard v.2.24.0 (http://broadinstitute.github.io/picard/), and reads were sorted and merged across distinct sequencing runs with samtools (v.1.10)^86^. Duplicates were removed using Picard’s *MarkDuplicates* function, and reads were locally realigned using *RealignerTargetCreator* and *IndelRealigner* from GATK (v.3.8.10)^87^. Bam quality was assessed with Qualimap (v.2.2.1)^88^, and capture statistics were obtained with Picard’s *CollectHsMetrics* tool (Supplementary Data S2 and S4). Samples with an average coverage of capture targets > 60X were randomly subsampled to an average coverage of 60X, to minimize biases in downstream analyses. Given the inclusion of historical samples in our dataset and the use of the USER enzyme to remove DNA damage introduced by cytosine deamination, MapDamage (v.2.2.0)^89^ was run to verify treatment success (Supplementary Fig. S17). For *Mustela erminea* samples, to which the USER treatment was not applied, MapDamage was used to recalibrate base quality scores for positions inferred to have damage patterns (*-- recalibrate* option).

Additionally, raw reads for five Mustelidae specimens were downloaded from the NCBI and ENA databases (Supplementary Data S3) and processed for adapter trimming, read mapping, read-group assignment, read sorting and merging, and realignment in indel regions, as described for capture data, with the following modifications: i) after adapter trimming, retain reads with a minimum length ≥ 36 bp; and ii) merge overlapping read pairs if minimum read overlap ≥ 15 bp. Bam quality was checked with Qualimap, bams were subsampled for reads mapping within capture targets, and coverage statistics within these regions were estimated with Picard’s *CollectHsMetrics* (Supplementary Data S4).

### Genotype and SNP calling

For each analysis, variant or genotype calls were produced if only SNPs or complete genotypes (SNPs and invariant positions) were required, respectively. Multi-sample calls were obtained with bcftools (v.1.10.2)^90^, with the *mpileup*, *call,* and *filter* options, requiring a minimum base (-Q) and mapping (-q) quality of 20, and applying the multiallelic caller (-c). Individual genotypes were filtered for a minimum depth of 6 (FMT/DP ≥ 6) and minimum genotype quality of 30 (FMT/GQ ≥ 30), and only sites with a maximum overall depth lower than two times that of the dataset under analysis (INFO/DP < 2*MEAN(INFO/DP)) were kept. We removed indels, SNPs with QUAL < 30, and any site with over 20% of missing data. For analyses based on anonymous capture regions (i.e., putative intronic and intergenic regions; see capture design above), we further masked SNPs within 5 bp of indels (--SnpGap). More stringent individual genotype filters were applied for population history and demographic analyses, requiring a minimum individual depth of 10 (FMT/DP ≥ 10). Samples with less than 10X average sequencing coverage were excluded from the dataset, which resulted in the exclusion of 15 samples, for a final dataset of five North American and 242 European least weasels, 27 additional *Mustela* and *Neogale* specimens, and two outgroups (see Supplementary Data S2 and S4).

### Species and gene tree inference

To address the evolutionary origin of *MC1R* variants, we used a dataset comprised of 12 *M. nivalis* specimens (selected to represent the diversity of haplotypes identified in the haplotype network analysis; Supplementary Data S2) and all additional Mustelidae samples (*Mustela*, *Neogale*, and outgroup species) to conduct phylogenetic analyses of genome-wide vs. local *MC1R* trees. Analyses were done for two sets of capture regions: i) a set of 2 kb anonymous regions (1,000 genome-wide fragments), and ii) one 40 kb region centered on *MC1R* causal SNPs (scaffold 337:145,924-185,924), defined based on homozygosity decay (Fig. 4C). VCF files were converted to geno files using the *parseVCF.py* script from the *genomics_general* suite (https://github.com/simonhmartin/genomics_general). Heterozygous sites were then transformed to IUPAC notation using the *filterGenotypes.py* script (*-of diplo*), and geno files were converted to fasta alignments using *genoToSeq.py*.

We estimated a genome-wide species tree using a coalescent framework, to account for variations in phylogenetic signals along the genome. First, a maximum likelihood phylogeny was inferred from each 2 kb anonymous fragment, using RAxML (v.8.2.12)^91^, under the GTRGAMMA substitution model and using 1,000 bootstrap replicates for node support values. We then calculated a tree certainty (TC) score for each fragment tree (*-L MRE* option), which allows weighting the support of the bipartition at each tree node against the second most frequently conflicting bipartition^92,93^. We retained only complete trees (with all individuals represented) with a relative TC score ≥ 0.30 (993 trees kept). Fragment trees were unrooted using the *unroot* function of the *ape* R package (v.5.4.1)^94^, and we used ASTRAL-III (v.5.7.8)^34^ to infer the consensus coalescent species tree. The tree was estimated with and without assigning individuals to species (*-a* option), and node support values were calculated using local posterior probabilities and quartet scores^95^.

The *M1CR* gene tree was estimated using the maximum likelihood approach from RAxML, with the same conditions used for anonymous fragment trees (GTRGAMMA substitution model; 1,000 bootstrap replicates). Additionally, genotypes at *MC1R* SNPs were collected for all Mustelidae samples from the VCF files (Supplementary Table S5).

### Population structure analysis

We assessed population structure within European *M. nivalis* (N = 242; sample subsets 1 and 2; Supplementary Data S1), using SNPs from anonymous capture regions. The inferred SNP dataset (in VCF format) was filtered to retain only the first SNP at each 25 kb along the genome (i.e., one per capture target), using Plink (v1.90b6.7)^96^ *--bp-space* option, to minimize non-independence due to linkage, resulting in 8,140 SNPs kept for analysis. A PCA of genome-wide data was conducted using the Plink *--pca* option, and genetic variation was summarized by plotting PC1 and PC2. Population clustering analyses were run using ADMIXTURE (v.1.3.0)^97^, testing K values ranging from 2 to 10, and the best K was selected using a 10-fold cross-validation error (Supplementary Fig. S5A). A neighbor-joining (NJ) tree of European least weasels was reconstructed under the BioNJ algorithm, using FastME (v.2.1.6.3)^98^. Tree inference was based on a matrix of pairwise genetic distances, which was first estimated with the Plink *--distance* option and expressed as genomic proportions (*1-ibs* matrix format). One European polecat (*Mustela putorius*) was included as outgroup (sample ID: NRM-MA605346; Supplementary Data S3).

### Population history and demographic inference

We next inferred the population history of European least weasels (sample subsets 1 and 2; Supplementary Data S1), to provide a neutral evolutionary model against which we could interpret results from local *MC1R* inferences. We first confirmed the relationships among populations inferred from ADMIXTURE, assuming three ancestral lineages (best K, Supplementary Fig. S5A), which we named North, West, and Central, based on their geographic distribution within our European sampling (Supplementary Fig. S5B). Specimens were assigned to a population if their ancestral proportion for a given cluster was above 0.51, resulting in the exclusion of a single individual (BNM12287, assignment North = 0.047; Central = 0.492; West = 0.461). Population relationships were inferred using the approach implemented in TreeMix (v.1.13)^99^. A dataset comprised of the retained European least weasels and three European polecats (outgroup population, Supplementary Data S3) was used to infer SNPs in anonymous capture regions. The resulting VCF was converted to bed format with Plink (*--bfile*), keeping a single SNP at each 25 kb (*--bp-space*). Allele frequencies per population were estimated using the Plink *--freq* option and then converted to the TreeMix input format with the *plink2treemix.py* script. TreeMix was run with sample size correction turned off (-*noss* option) and estimating the standard errors for each run (*-se*).

We next used ∂a∂i (v. 2.3.2)^39^ to infer the joint site frequency spectrum (jSFS) of each population pair. SNPs were inferred along putative intergenic capture regions and filtered to retain a single SNP per capture target (25 kb distance). To account for missing data, reduce computational time, and improve jSFS inference, we down-projected our dataset to 25% of the original population size (North: N = 56 individuals and 112 alleles; West: N = 28 and 56 alleles; Central: N = 36 and 72 alleles), using a folded jSFS approach. Final down-projected jSFS datasets per population pair were as follows: North and West – 4,987 SNPs; North and Central – 5,039 SNPs; West and Central – 4,869 SNPs.

To infer demographic parameters, including effective population sizes (N_e_), divergence times (t), and migration rates (m) among populations, we ran ∂a∂i as implemented in GADMA (v.2.0.0)^40,100^. This method allows the unsupervised inference of the best demographic model through a two-step approach^40^: first, inference of the best model structure is conducted through a global search approach based on the principles of the genetic algorithm; then, optimization of parameters for the best model structure is done using a demographic inference software, here ∂a∂i. We first tested a three-population model, in which population split patterns were set following the population relationships inferred from TreeMix (Fig. 3D; Supplementary Fig. S6A). However, our data was insufficient to achieve convergence into the same set of parameter values across replicate runs. Therefore, we opted to focus our model inference on the most basal divergence identified among our dataset, i.e., the split between North and Central lineages (Fig. 3; Supplementary Fig. S6). In total, 100 replicate runs of GADMA were performed, checking for convergence of parameter estimates, and the best run was selected based on likelihood values and model comparison with the Akaike Information Criterion (AIC), as advised for datasets of unlinked SNPs. Confidence intervals for parameters of the best model were inferred using a non-parametric bootstrap approach, randomly sampling 100 bootstrap datasets from the original SNP set and applying the same down-projection used in the main run. Parameter optimization was run for each bootstrap using the *gadma_run_ls_on_boot_data* script, and 95% confidence intervals were calculated using point estimates ± 1.96 SD (standard deviation). Raw parameter values were converted assuming a general mammalian mutate rate of 2.2 x 10^-9^ (ref.^101^) and a generation time of one year^18^.

To evaluate the fit of the best demographic model, comparisons of the empirical jSFS with the expected under the best model were plotted using ∂a∂i, under a multinomial distribution (Supplementary Fig. S7A). Additionally, we compared the distribution of empirical summary statistics to those simulated under the best model parameter estimates (Supplementary Fig. S7B). To obtain empirical distributions, genotype calls for 1,000 genome-wide anonymous 2 kb capture fragments were converted to geno files using the *parseVCF.py* script from the *genomics_general* suite. Pairwise F_ST_ between the North and Central populations and Tajima’s D within each lineage were estimated with the *popgenWindows.py* script, for each 2 kb fragment, filtering for ≥ 10 sites per fragment (*-m*) and ≥ 50% of individuals with data (*--minData*). Simulated distributions based on the best demographic model were obtained with msprime (v.1.2.0)^102^. We simulated 1,000 fragments of 2 kb length, assuming the same mutation rate and generation time as above, and estimated pairwise F_ST_ and per population Tajima’s D per fragment. Input files used in demographic analysis and the msprime simulations and outputs of the demographic inferences and simulations are deposited in Figshare^69^.

### Genotype-to-phenotype mapping

To explore if the two coloration traits that distinguish the *nivalis* and *vulgaris* morphs (dorsoventral line and winter coat color) are defined by a single locus or two tightly linked loci, both hypotheses consistent with previous crossing experiments^23^ and genomic analyses^22^, we used the full capture sequencing data to perform genotype-to-phenotype association mapping, using Plink (v1.90b6.7)^103^. European specimens were grouped into phenotypic groups, considering only those with *a priori* confirmed phenotypic information (sample subset 1; Supplementary Data S1). For the dorsoventral line, only specimens harboring a brown coat (irrespective of sampling season) that allowed identifying the line type were considered. For the winter color, we included any specimens with winter-white coats or specimens with brown coats and a collection date between December and February^50^ to minimize the misclassification of winter-brown color that could result from variations in autumn or spring molt timings. This resulted in a total of 121 individuals scored for line type (straight, N = 35; ragged, N = 86; 204,777 SNPs capture-wide) and 106 scored for winter color (white, N = 66; brown, N = 40, 156,141 SNPs; Supplementary Data S2). For each dataset, we conducted a case-control association test (*--assoc*) to assess if causal *MC1R* SNPs^22^ were the strongest associated variants in both phenotypic partitions or if other variants would arise. Because no other strong candidates were identified (Supplementary Fig. S8), associations were run for a complete dataset of *a priori* phenotyped *nivalis* or *vulgaris* specimens, based on either the winter color and/or the dorsoventral line phenotype, for a total of 188 individuals (*nivalis,* N = 101; *vulgaris*, N = 87; 218,609 SNPs). A case-control association test, a Fisher’s exact test (*--assoc fisher*), and a Cochran-Armitage trend test, which does not assume Hardy-Weinberg equilibrium (*--model*), were conducted for this dataset. In the figures, we report the results for the case-control association test, but results were consistent across analyses (Supplementary Fig. S9). Significant associations were defined using a Bonferroni-corrected P-value < 0.05.

Given the perfect association found between *MC1R* SNPs and both color traits, as well as the lack of other candidate variants, genotypes at those SNPs were used to infer the coloration morph (*nivalis* or *vulgaris*) of individuals with *a priori* uncertain phenotype, i.e. sample subset 2 (see *Genetic sampling strategy and data collection*; Supplementary Data S1; Supplementary Table S6). The inferred morph was checked against the expected morph distribution^13^ and considered for downstream analyses of local *MC1R* variation that relied on morph classification. Additionally, *MC1R* genotypes for North American least weasels were also recovered from VCF files (Supplementary Table S6).

### Haplotype phasing

To explore *MC1R* haplotypic diversity and local patterns of variation, we statistically phased haplotypes in the *MC1R* capture region (337:0-500,000) across the entirety of the European *M. nivalis* dataset (sample subsets 1 and 2; Supplementary Data S1), using a two-step approach. First, we used WhatsHap (v.1.3)^104^ with default settings to phase genomic variants per individual, using a read-based phasing approach that considers linkage information from sequencing reads to inform haplotype phases. Then, we used ShapeIT (v.4.2.2)^105^ to perform population-level phasing, using WhatsHap results as input (*--use-PS* option) and with the following iteration scheme (*--mcmc-iterations*): 10b, 1p, 5b, 1p, 2b, 1p, 1b, 1p, 1b, 1p, 10m. A phased VCF file was obtained as output and is available in Figshare^69^.

### Haplotype homozygosity analysis

We used the *rehh* R package (v.3.2.2)^42^ to explore haplotype homozygosity decay in the flanking regions of *MC1R* variants. The phased VCF was subsampled to include the complete dataset of *a priori* phenotyped individuals (N = 188; sample subset 1; Supplementary Data S1) used in the genotype-to-phenotype mapping. Alleles were identified as *nivalis* or *vulgaris* based on a consensus call of *a priori* phenotyped *nivalis* specimens (N = 101; Supplementary Data S1). To do so, variant calls were produced for this sampling subset as described above; a consensus sequence of majority alleles (INFO/AF > 0.5) for the *nivalis* morph was produced with *bcftools consensus* and used to inform REF and ALT alleles. The VCF file was used as input to the *data2haplohh* function of the *rehh* package. Extended haplotype homozygosity (EHH) was calculated for SNP 337:165,924 (the first of the two consecutive causal *MC1R* SNPs) using the *calc_ehh* function, with the parameter ‘discard_integration_at_border = FALSE’. The code used for analysis is deposited in Figshare^69^.

### Intraspecific haplotype diversity

To further explore haplotype diversity along the *MC1R* association region, phased haplotypes were used to infer a haplotype network, using a total of 484 haplotypes from all European least weasels (sample subsets 1 and 2; Supplementary Data S1). A target region of 40 kb, centered on *MC1R* SNPs (scaffold 337:145,924-185,924), was defined based on EHH decay along the association region (Fig. 4C). Due to the high number of haplotypes included in the analysis, variant sites were filtered for a minimum minor allele frequency (MAF) of 0.10, to reduce haplotype network complexity for visualization, but the resulting patterns were consistent with less stringent MAF filters. The VCF file was converted to a fasta sequence alignment using the scripts *parseVCF.py* and *genoToSeq.py* from the *genomics_general* suite. The resulting alignment was used as input for network reconstruction using PopART (v.1.7)^106^, under a median-joining algorithm^107^. Haplotype structure was additionally visualized in a haplotype heatmap, using a custom R script (available in Figshare^69^). Samples were clustered by population lineage and color morph to allow visualization of geographic haplotype distribution (Supplementary Fig. S10).

### Haplotype divergence times and allele age

We used the Bayesian inference approach implemented in BEAST (v.1.10.4)^43^ to date *MC1R* haplotype divergence in *M. nivalis* within the context of other mustelid species. To do so, four *M. nivalis* homozygous for each *MC1R* haplogroup-allele combination (haplogroup 1 – HG1, with *nivalis* and *vulgaris* allele; and haplogroup 2 – HG2, with *nivalis* and *vulgaris* allele; Fig. 4D; Supplementary Fig. S10) were selected to be included in the analysis (Supplementary Data S2). The *M. nivalis* phased VCF was used to produce a fasta sequence per haplotype per specimen, using bcftools *consensus* (v.1.11) to mask missing data based on the genotype call. A subset of Mustelidae samples was additionally included to represent the main lineages identified in the genome-wide species tree (Fig. 2A), for a total of eight *Mustela* and *Neogale* species (one specimen each) and two outgroups (Supplementary Data S4). One consensus fasta sequence was produced per individual, from the unphased VCF file, with bcftools *consensus*, masking sites with missing calls, and using the IUPAC notation for heterozygous positions. Variant sites were then randomly phased using the *randbase* option of seqtk (v.1.3; https://github.com/lh3/seqtk) to select a single base per heterozygous site. Sequences were merged in a single file, the complete alignment was visually inspected using MEGA11^108^, and sites with more than 50% missing data were removed with TriSeq (v.1.0.2), from Trifusion (https://github.com/ODiogoSilva/TriFusion). A region of 40 kb centered on *MC1R* SNPs was kept for analysis, based on EHH decay (see above). BEAST was run assuming a strict molecular clock and a GTR substitution model. The Yule process of speciation model was used as a tree prior, and a gamma distribution was used as the prior for the clock rate. Node ages were estimated based on two calibration points, both defined based on fossil records of mustelid species, available in the NOW fossil database^109^: i) a normal distribution of 7.1 ± 1 Mya (mean ± SD) for the ingroup node age (including both *Mustela* and *Neovison* species), and ii) a normal distribution of 3.5 ± 1 Mya for the *M. erminea* node age. Three independent runs of 50 million generations were conducted, sampling trees and parameters every 50,000 generations. Convergence was assessed with Tracer (v.1.7.2)^110^, and the final tree was reconstructed using the combined results of three independent replicates, discarding the first 10% of the run as burn-in. BEAST input and output files are deposited in Figshare^69^.

The derived *vulgaris MC1R* variants were additionally dated following the hidden Markov model (HMM) approach implemented in GEVA (v.1beta)^37^. The complete set of phased European *MC1R* haplotypes was used as input for the analysis. We assumed a mutation rate (μ) of 2.2 x 10^-9^ (ref.^101^) and a recombination rate of 1 x 10^-8^ (based on dog estimates^111^). A global effective population size (N_e_) of 435,466 was assumed based on anonymous nucleotide diversity (π) estimates (see next section) and following the equation N_e_ = π / 4μ. A maximum of 2 k concordant and discordant haplotype pairs were allowed, and we report estimates based on a joint clock model, filtering outlier pairs.

### Genetic diversity and divergence

To explore genetic diversity among specimens with distinct *MC1R* haplotypes, nucleotide diversity (π) and Tajima’s D were calculated for each *MC1R* haplogroup-allele combination (HG1, with *nivalis* and *vulgaris* allele; and HG2, with *nivalis* and *vulgaris* allele; Fig. 4D). For each combination, only homozygous specimens were included (HG1-*vulgaris*, N = 15; HG1-*nivalis*, N = 15; HG2-*vulgaris*, N = 71; HG2-*nivalis*, N = 88), and the complete dataset of European least weasels was used for comparison purposes (N = 242; sample subsets 1 and 2; Supplementary Data S1). Genotype calls for 1,000 genome-wide anonymous 2 kb capture fragments were used as input for the *genomics_general* suite and converted to geno files using the *parseVCF.py* script. The *popgenWindows.py* script was used to estimate nucleotide diversity and Tajima’s D in each 2 kb fragment, for the five datasets described (*--windType* predefined), requiring ≥ 10 sites per fragment (*-m*) and ≥ 50% individuals with data (*--minData*). Estimates were additionally conducted for a 40 kb region surrounding *MC1R* causal SNPs (scaffold 337: 145,924-185,924) for the five datasets, in 2 kb non-overlapping windows, using the same procedure and filtering criteria. We also calculated absolute genetic divergence (D_xy_) between pairs of *MC1R* haplogroup-allele datasets for both window sets (genome-wide and *MC1R*), following the same steps.

### Selection analysis

To test for signatures of positive selection along the *MC1R* capture region, we used the composite likelihood ratio (CLR) approach implemented in SweeD (v.4.0.0)^112^. Inferences were conducted independently for each *MC1R* haplogroup-allele dataset (HG1, *nivalis* or *vulgaris* allele; HG2, *nivalis* or *vulgaris* allele; homozygous specimens only) and the complete datasets of *nivalis* and *vulgaris* coloration morphs (Supplementary Table S6). We first used a VCF file of anonymous genome-wide capture regions to estimate an SFS per scaffold with SweeD, using the *-osfs* output option and keeping only polymorphic sites (*-strictPolymorphic*). One European polecat sample (*M. putorius*; sample ID: NRM-MA605346) was used to polarize the VCF calls. The resulting SFS files per scaffold were then summed to generate a single genome-wide SFS. After, a VCF file of scaffold 337 was used as input to run SweeD along that genomic region, estimating the CLR at ∼10 kb intervals along the scaffold (*-grid* 4139), using the inferred genomic SFS as the baseline against which to estimate sweeps, and keeping only polymorphic sites (*-strictPolymorphic*). CLR outliers were defined based on the 95th percentile of the empirical distribution of CLR values.

We inferred the selection coefficient of the identified selective sweep (Supplementary Fig. S15), following the same approach used in ref.^22^. We first ran SweeD on 1,000 randomly sampled bootstrap replicates of scaffold 337, sampling 50% of the scaffold SNPs. For each replicate, we estimated the mean value of alpha (α) in a 250 kb region including MC1R (scaffold 337:115,000-365,000), defined based on the number of consecutive windows surrounding *MC1R* that fell above the CLR outlier threshold. For each bootstrap replicate, we estimated the selection coefficient (s) using the equation s = r × ln(2 N_e_)/α ^113^ (where r is the recombination rate per base per generation and N_e_ the effective population size), generating a posterior probability distribution of s. We randomly sampled the recombination rate from a uniform distribution based on mouse and human means (0.56 – 1.26 cM/Mb; see ref.^114^) and N_e_ from a uniform distribution based on the 95% confidence interval of N_e_ estimates for the North lineage of European least weasels inferred in our demographic modeling (349,140 – 386,651; see Supplementary Table S7).

### Forward-in-time genetic simulations

To explore if genetic drift alone could maintain the *MC1R* polymorphism segregating in least weasel populations in Europe during ∼1 million years, we conducted Wright-Fisher forward-in-time simulations using SLiM (v.4.2)^44^. We assumed a neutral genetic drift model with two main genetic lineages splitting from a common ancestral, representative of the most basal divergence identified in our population tree (Fig. 3D). Model parameters, including effective population sizes, split times, and growth and migration rates, were informed by our demographic modeling of the Central and North lineages, based on empirical data (Fig. 3E). We seeded a single neutral derived dominant variant in the ancestral population at generation 1, followed by an ancestral population split at generation 2, generating the North and Central populations, followed by independent evolution of each population for 1,007,481 generations, until the present time. Demographic parameters were scaled by a factor of 5 due to the computational time unfeasibility of an unscaled model. We varied the initial frequency of the derived variant in the ancestral population to allele frequencies of 0.051, 0.163, and 0.293, representative of derived phenotype frequencies, i.e., frequency of *vulgaris* morph in the population – f(*vulgaris*), of 10, 30, and 50% respectively, under Hardy-Weinberg equilibrium. This approach allowed us to test distinct possibilities of *vulgaris* phenotype frequency in the ancestral population at the onset of population divergence.

A total of 400 replicates were run for each pre-split f(*vulgaris*). At the end of each replicate, we collected information about allele and genotype frequencies in both lineages. These data were used to infer *vulgaris* phenotype frequencies (considering *vulgaris* allele dominance) at the end of the simulation and compare them with *a priori* expectations for a segregating polymorphism. Our sampling, though not systematic and limited to the specimens available at sampled collections, suggests an intermediate probability of the derived *vulgaris* phenotype within each genetic lineage. The expectation of intermediate frequencies agrees with the probability of occurrence of alternative winter colors estimated by previous modeling of phenotype distribution against environmental variables^13^. When considering a 200 km buffer around our sampling within each genetic lineage, this model suggests a probability of occurrence of the winter-brown *vulgaris* phenotype of about 25% and 77% in the North and Central lineages, respectively (Supplementary Table S9). A final f(*vulgaris*) range between 0.20 and 0.80 was used to represent the maintenance of the alternative color morphs segregating in the populations at the end of the run. The fit of the models to these expectations was thus expressed as P, i.e., the proportion of simulation replicates in which f(*vulgaris*) was kept within the defined range in both genetic lineages at the end of each simulation. Input files used in forward-in-time simulations and resulting outputs are available in Figshare^69^.

## ACKNOWLEDGMENTS

This work was supported by Fundação para a Ciência e a Tecnologia (FCT) project grant under the ERC-Portugal programme to J.M.-F.. I.M. and M.A. were supported by FCT PhD grants (SFRH/BD/143457/2019 and 2021.05642.BD, respectively; funds from the European Social Fund (ESF) and Portuguese MCTES/FCT), and J.M.-F. was supported by an FCT research contract (2021.00150.CEECIND). C.R.F. thanks the support of CE3C through an assistant researcher contract (FCiência.ID contract #366) and FCT for Portuguese National Funds attributed to CE3C within the projects UIDB/00329/2020, UIDP/00329/2020, and LA/P/0121/2020, and FPUL for a contract of invited assistant professor. M.S.-R. was supported by FCT (UIDB/00329/2020). L.D. acknowledges support from the Swedish Research Council (2021-00625) and the European Union (ERC, PrimiGenomes, 101054984). Additional support was obtained from the European Union’s Horizon 2020 Research and Innovation Programme under the Grant Agreement Number 857251. Sampling at the Swedish Museum of Natural History (NRM, Stockholm) was supported by the SYNTHESYS program (access grant SE-TAF-4695 to J.M.-F.; EU FP7 Agreement 226506). We thank museum collections and their curators and staff for providing sample access and/or support at collections: Christian Montermann and Jan Decher (ZFMK, Zoological Research Museum Alexander Koenig, Mammal Section); Hermann Ansorge (SMNG, Senckenberg Museum of Natural History Goerlitz, Mammalogy Collection); Neil Duncan and Marisa Surovy (AMNH, American Museum of Natural History, Department of Mammalogy); Mark Omura and Breda M. Zimkus (MCZ, Museum of Comparative Zoology at Harvard University, Department of Mammalogy and MCZ-CRYO); Darrin P. Lunde and John Ososky (USNM, Smithsonian National Museum of Natural History, Division of Mammals); Paolo Viscardi and Aidan O’Hanlon (NMI, National Museum of Ireland, Natural History); the Swedish Museum of Natural History (NRM, Department of Zoology); and the Bündner Naturmuseum (BNM). We thank P. C. Alves and EVOCHANGE members for helpful discussions. Procedures were revised and approved by the Organism Responsible for the Well-Being of Animals (ORBEA) of CIBIO-InBIO, University of Porto (reference number 2024-07).

## Data availability

Capture-based raw sequencing data will be deposited in the NCBI Sequence Read Archive (SRA) database. Functional assay data, phenotypic measurements, capture design details, analysis code, simulation outputs, and custom scripts will be deposited in Figshare. Custom code will also be made available on GitHub.

## Author contributions

J.M.-F. coordinated the study. I.M., R.R., J. M. G., L.S.M., L.F.C.C., L.D., and J.M.-F. conceptualized and designed the research. Z.B., M.E., D.C.K., J.M., J.P.M., L.S., F.S., J.S., M.R., M.S.-R., C.R.F., K.Z. provided key sampling resources. I.M., R.R., L.F., and M.A. performed laboratory work. I.M. and R.R. analyzed data. I.M. and J.M.-F. wrote the paper. All authors commented, revised, and approved the final version of the manuscript.

