## Supplementary Information for "Ancient balanced polymorphism underlies long-standing adaptation for seasonal camouflage in the least weasel"

**Contents:**

Supplementary Text
Supplementary Figs. S1 to S17
Supplementary Tables S1 to S9
Legends for Supplementary Data S1 to S4
References

### Supplementary Text

#### Text S1 - Phenotyping of museum skins

Having confirmed that the derived *MC1R* alleles identified in *vulgaris* least weasels result in altered receptor response to its ligands (see Main text; Fig. 1C-D), we investigated whether this could be associated with differences in least weasels' brown coat colors. We thus focused on testing potential differences among phenotypically brown individuals (i.e., summer *nivalis*, summer *vulgaris*, and winter *vulgaris*; Fig. 1A), to understand if altered receptor function could result in distinct brown tones in *vulgaris* specimens, either among seasons or when compared to *nivalis* individuals. We used a standardized digital photography approach to characterize phenotypic differences among specimens classified into four phenotypic groups: summer *nivalis*, winter *nivalis*, summer *vulgaris*, and winter *vulgaris*. Principal component analysis (PCA) of all phenotypic data shows that principal component 1 (PC1; 96.62%; Supplementary Fig. S1B) splits specimens into two main groups, which cluster according to the fur color: white (winter *nivalis*) or brown (all remaining specimens). Next, we removed winter *nivalis* specimens from the dataset and redid the analysis on all brown specimens and pairs of phenotypic groups, using a biplot approach to identify the coloration variables that explain the differentiation among PCA axes. Results from all PCA configurations show that the variables that most influence variation on PC1 are those related to hue and value, in the HSV (Hue, Saturation, and Value) color space, whereas variation along PC2 is primarily explained by saturation data (Supplementary Fig. S1C-F). No statistically significant differences were observed between *nivalis* and *vulgaris* summer morphs, for any color measurements (Bonferroni-corrected $P > 0.05$; Kruskal-Wallis test). On the contrary, winter *vulgaris* significantly differed from both summer *vulgaris* ($P < 0.01$) and summer *nivalis* ($P < 0.05$) for hue (H) measurements in all sampling points (Supplementary Fig. S2). Additional significant differences were found in saturation (S) values from the upper (UD) and middle (MD) dorsum, between winter *vulgaris* and other brown specimens ($P < 0.05$; Supplementary Fig. S2). Overall, these results point to a difference in the underlying pigment (here, expressed through hue) and a partially purer color (expressed through higher saturation) of winter *vulgaris* compared to other brown specimens, which are expected to result in lighter coat color.

#### Text S2 - Putative annotation of the *M. nivalis* reference genome

To select the anonymous genome-wide regions to include in the custom capture design, we started by defining putative intergenic and intronic regions along the *Mustela nivalis* reference genome (GCA_019141155.1)[1], to which annotation is currently unavailable. First, a repeat library for *M. nivalis* genome was created using RepeatModeler (v.2.0)[2], and repetitive regions were masked using RepeatMasker (v. open-4.0.9)[3]. Then, a draft annotation of the assembly was conducted using MAKER (v.3.01.03)[4]. Given that no transcriptomic data was available for *M. nivalis*, an initial MAKER run was executed using ferret (*Mustela putorius furo*) mRNA data (retrieved from GenBank, accessed on April 28, 2020), and proteomic data from four Carnivora – ferret, dog (*Canis lupus familiaris)*, cat (*Felis catus)*, and giant panda (*Ailuropoda melanoleuca)* – from the UniProt database (accessed on April 28, 2020), as protein homology evidence from closely related species, by enabling the option *protein2genome*. The resulting annotation was used to build a gene model with SNAP (v.2006-07-28)[5]. The annotation output was converted to SNAP format using the *maker2zff* script from MAKER distribution, using the following parameters: -c 0 -e 0 -o 0.5 -a 0 -t 0 -l 50 -x 0.3. SNAP scripts *fathom* and *forge* were used to prepare the input for model building, and the *hmm-assembler.pl* script was used to build the model; all scripts were run with default parameters. A second MAKER run was then conducted by disabling the *protein2genome* option and using as a single input the built SNAP gene model. Resulting gene predictions were merged in a single gff file with the *gff3_merge* script from MAKER and used for defining putative intergenic and intronic regions along the *M. nivalis* genome.

Intronic regions were defined by subtracting the coordinates of predicted exons from the coordinates of predicted gene sequences between the first and last exon, using bedtools subtract (v.2.29.2)[6]. Similarly, intergenic regions were defined by subtracting the coordinates of predicted genes from the entirety of the genome. To account for potential regulatory regions (e.g., untranslated regions [UTRs] and promoters) occurring in proximity to coding sequences, which are not predicted by the SNAP gene model, coding sequence coordinates were extended by 1 kb in each direction, considering mammal UTR length estimates[7].

The obtained lists of regions were filtered to remove repeat regions, as previously inferred by RepeatMasker. To filter regions for multiple genome-wide copies of the same genomic fragment (e.g., paralogous regions), that might bias the inference of polymorphisms in downstream analyses (e.g. ref.[8]), each dataset (intronic and intergenic) was aligned against the *M. nivalis* reference genome using BLAT (v.36x7)[9], considering hits with a minimum homology of 80%, and we only kept regions with a single hit and no gaps in the reference genome sequence. For each retained region, SNPs were estimated with ANGSD (v.0.923)[10], using a set of 79 previously processed *Mustela nivalis* low-coverage whole-genome sequences (see details in ref.[1]), with the following filtering parameters: -minQ 20 -minMapQ 20 -minInd 30 -setMaxDepth 289 -SNP_pval 1e-6 -minMaf 0.006. Distributions of SNP density were inferred for both intronic and intergenic datasets, and outlier regions at 5% in both extremes of the distribution were also excluded. Finally, given that elevated GC content has been shown to affect the efficiency of enrichment assays[11], the percentage of GC content in the remaining regions was estimated for each dataset, and outliers at 5% in the right tail of the distribution of GC content values were removed.

**Text S3 - Modifications to the protocol for genomic library preparation**

Double-indexed genomic libraries were prepared using the protocol of ref.[12], with the modifications of ref.[13] and additional changes, described below.

For historical/lower-quality samples (dry skin patches), DNA fragment size was initially checked by electrophoresis and, for samples with higher DNA integrity, DNA was sheared to ~350 bp with Bioruptor Pico. For more degraded samples, no shearing step was performed. Up to 1 µg of DNA was used for blunt-end repair, adapter ligation, and adapter fill-in. The USER enzyme (New England Biolabs) was used in the blunt-end repair for all skin samples (except *Mustela erminea*; see below) to account for cytosine deamination, as in ref.[13]. The number of cycles for the indexing polymerase chain reaction (PCR) was determined through quantitative PCR (qPCR). The indexing PCR was modified by using AccuPrime Pfx DNA polymerase (Invitrogen), an enzyme with proofreading activity, and including four to six PCR replicates per sample, to increase library complexity. Reactions included 1 X AccuPrime Pfx buffer, 0.4 μM P5 and P7 indexing primers, 0.025 U/μL AccuPrime Pfx DNA polymerase, and 6 μL DNA template, for a total reaction volume of 25 μL. For *Mustela erminea* samples, the USER enzyme was not used in blunt-end repair, and the indexing PCR was performed with the KAPA Hifi DNA Polymerase (Roche). Reactions included 1 X KAPA Hifi HotStart U+ ReadyMix, 0.2 μM P5 and P7 indexing primers, and 6 μL DNA template, for a total volume of 25 μL.

For modern/higher-quality samples (ethanol-preserved tissues), DNA was sheared to ~350 bp. Library preparation was done with up to 1 μg of DNA per sample, and the USER enzyme was not included. Indexing PCRs were done with Herculase II Fusion DNA polymerase. Reactions included 1 X Herculase II reaction buffer, 0.25 mM each dNTP, 0.2 μM P5 and P7 indexing primers, 0.5 μL Herculase polymerase, and 3.5 μL DNA template, for a total reaction volume of 50 μL. Four PCR replicates were used per sample.

### Supplementary Figures

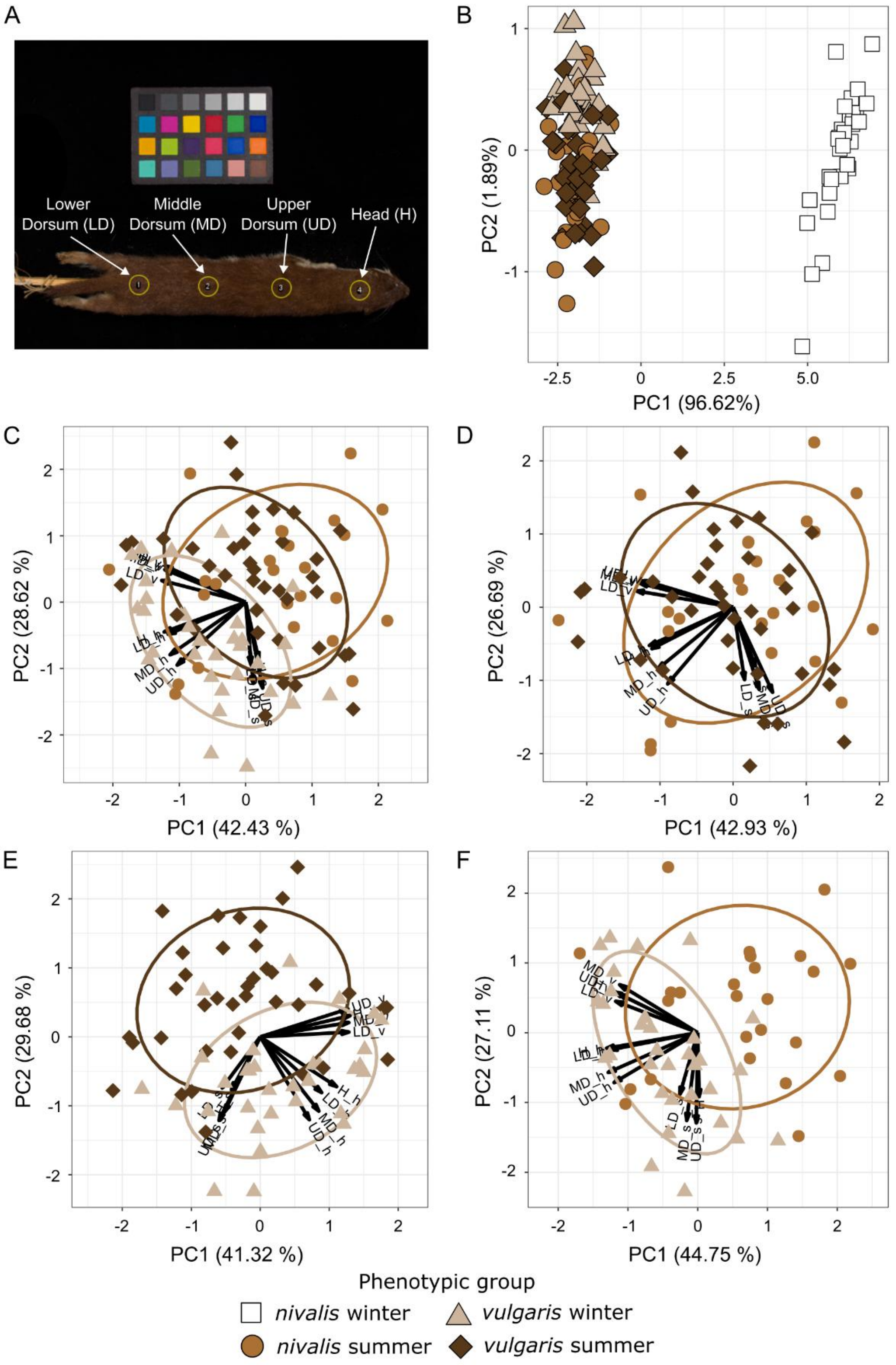


**Fig. S1 - Phenotyping of European least weasel skins. (A)** Sampling points used to retrieve color measurements of 134 least weasel skins from the NRM collection. Four sampling points were defined along the dorsal midline of each specimen. **(B)** Principal component analysis (PCA) of phenotypic data collected from 125 specimens without missing data, representative of *vulgaris* and *nivalis* morphs, collected during both winter and summer (winter *nivalis*, N =

30; summer *nivalis*, N = 25; winter *vulgaris*, N = 32; summer *vulgaris* N = 38). **(C-F)** Biplots of color measurements recovered for **(C)** phenotypically brown individuals (i.e., summer *vulgaris*, winter *vulgaris*, summer *nivalis*; Fig. 1A); **(D)** summer specimens (both *nivalis* and *vulgaris*); **(E)** *vulgaris* specimens (both from summer and winter); and **(F)** summer *nivalis* and winter *vulgaris* specimens. In each biplot PCA, black arrows show variable loadings in the principal components. Variables are labeled by sampling point (as in panel A) and color measurements (hue – H, saturation – S, value – V) [e.g., UD_s is the saturation value in the upper dorsum sampling point].

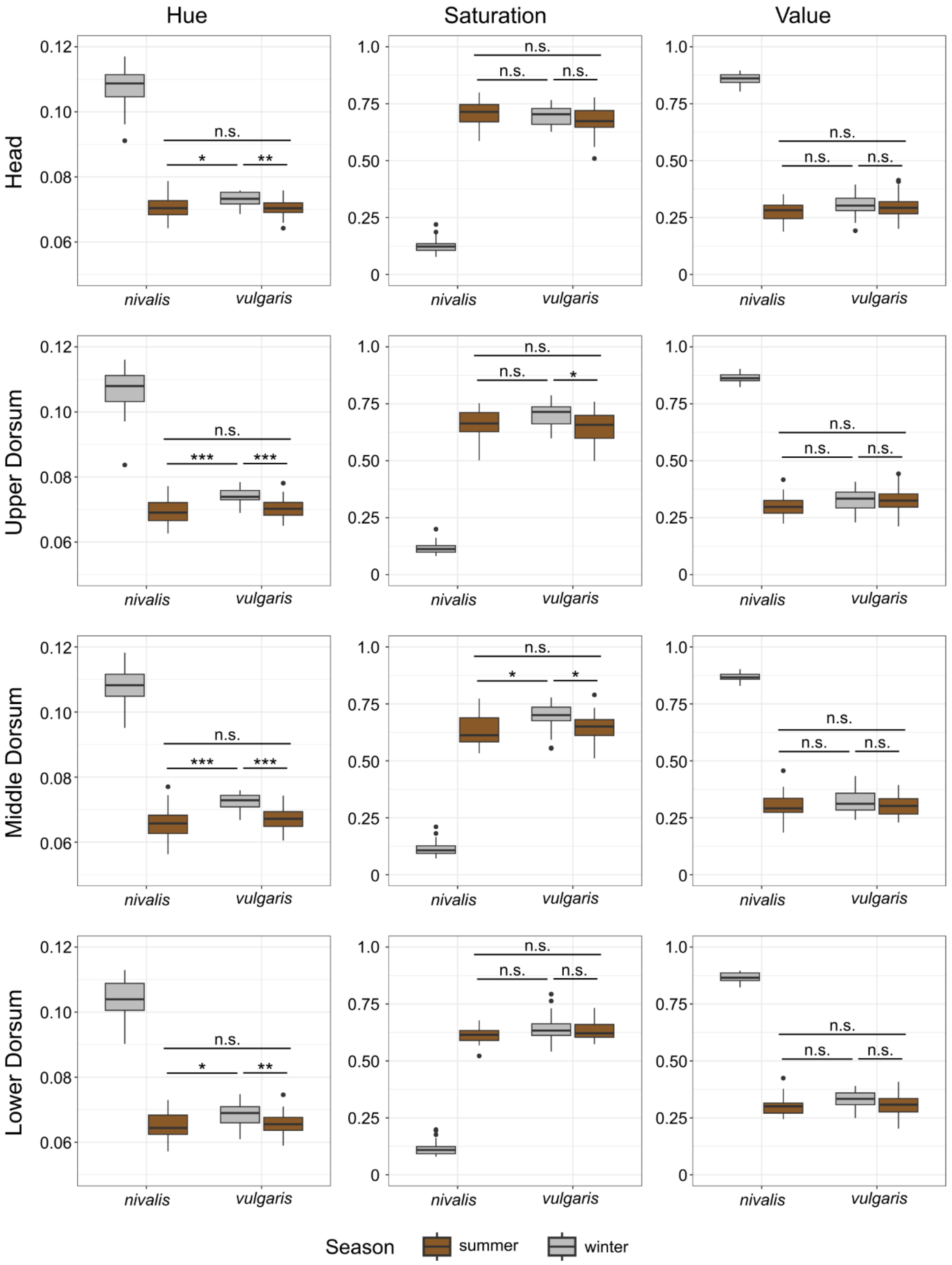


**Fig. S2 - Hue, Saturation, and Value measurements of European least weasel skins.** Boxplots show the distribution of color measurements obtained from 125 least weasel skins. Data on Hue, Saturation, and Value (HSV color space) were obtained for four sampling points (Supplementary Fig. S1A). Specimens were grouped into four phenotypic groups according to color morph (*nivalis* or *vulgaris*) and collection season (summer or winter). The significance of measurement differences among phenotypically brown specimens was assessed with a Kruskal-

Wallis test and is denoted in each panel as: n.s. – $P > 0.05$; * – $P < 0.05$; ** – $P < 0.01$ (Bonferroni-corrected P-values).

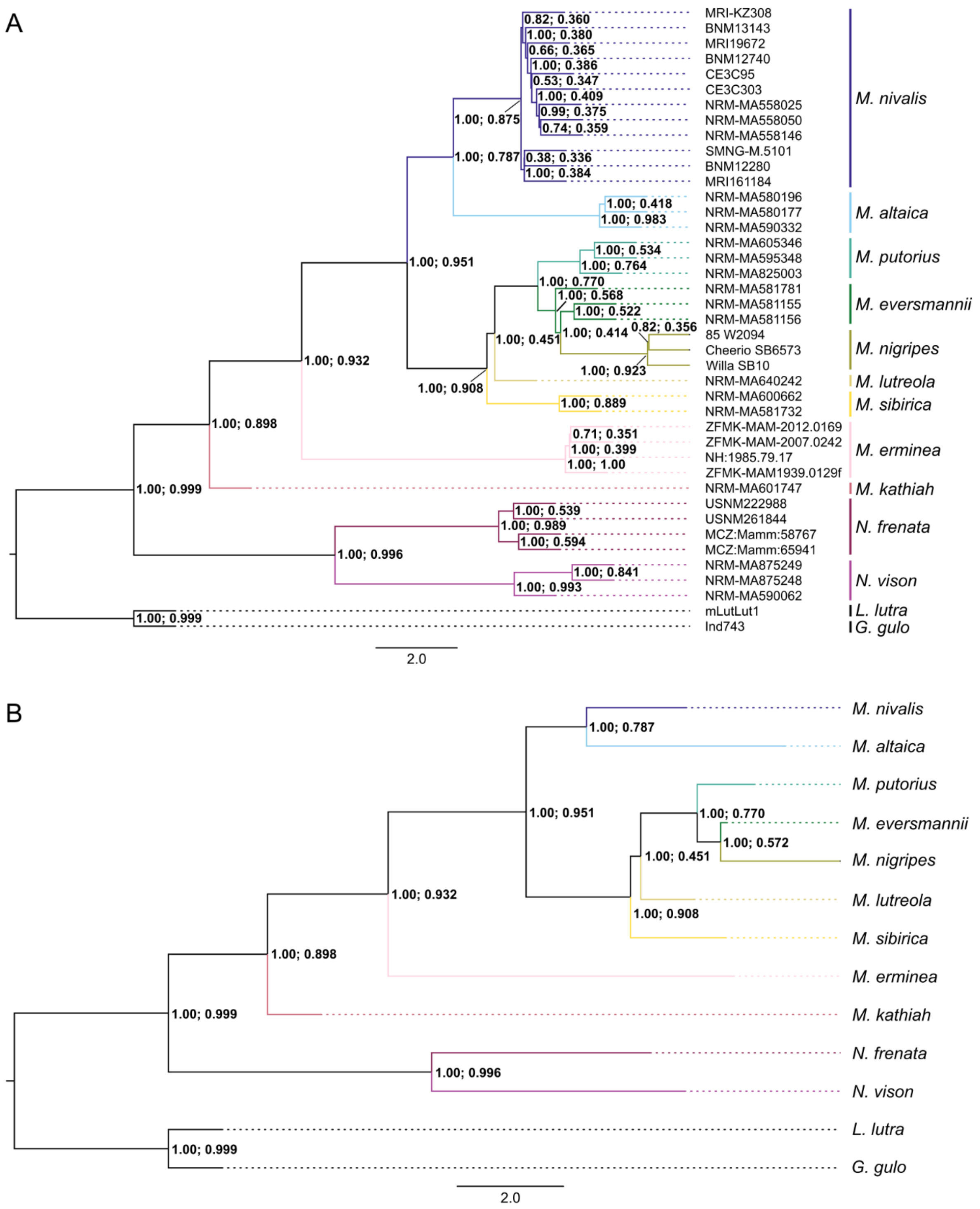


**Fig. S3 - Mustelidae species tree.** Coalescent species tree, inferred with ASTRAL-III, based on 993 capture fragment trees (~1.91 Mb of genomic sequence), **(A)** without or **(B)** with individuals' assignment to species. Node annotations in both trees show posterior probabilities and quartet scores support per node, respectively. *Lutra lutra* and *Gulo gulo* were used as outgroups.

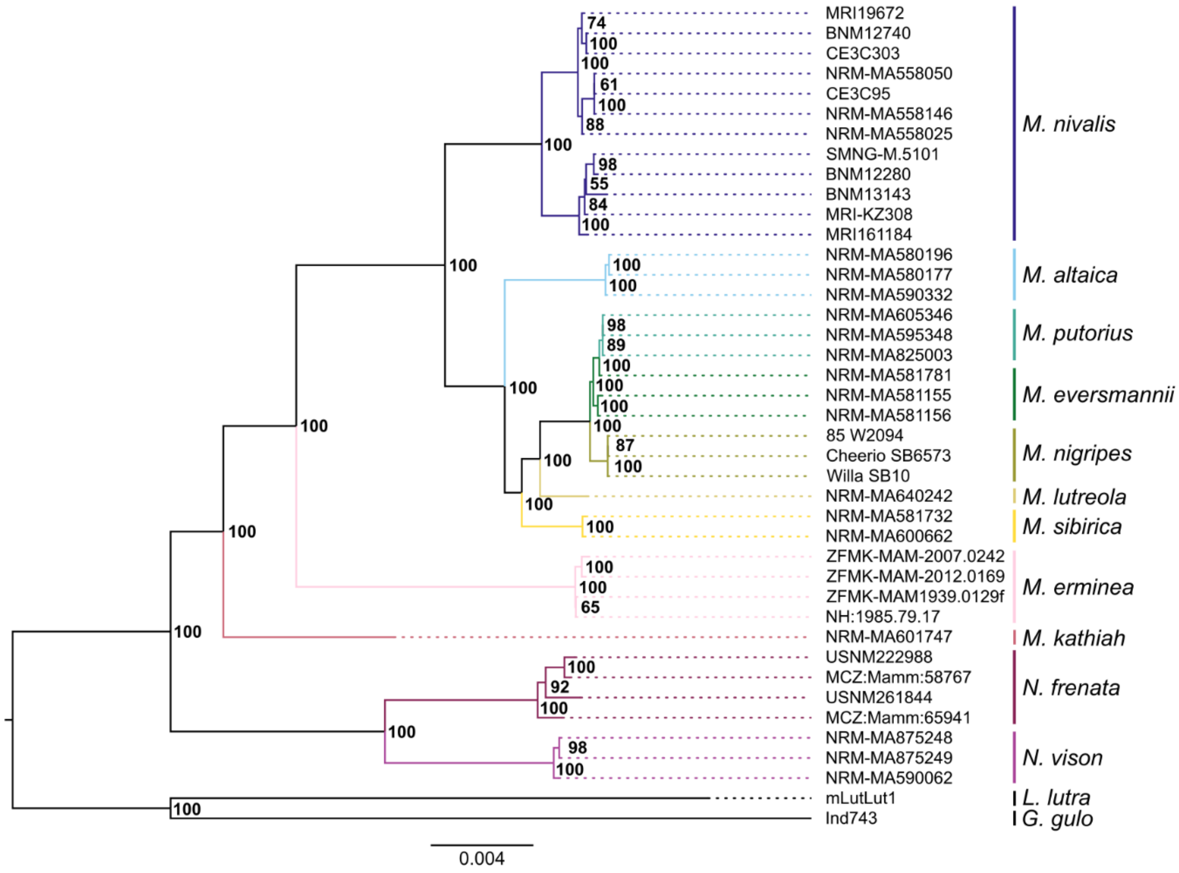


**Fig. S4 - Mustelidae *MC1R* gene tree.** Maximum likelihood gene tree, inferred with RAxML, based on a 40 kb region, centered on causal *MC1R* SNPs (Figs. 1C and 4C). Node annotations show bootstrap support values, based on 1,000 bootstrap replicates. *Lutra lutra* and *Gulo gulo* were used as outgroups.

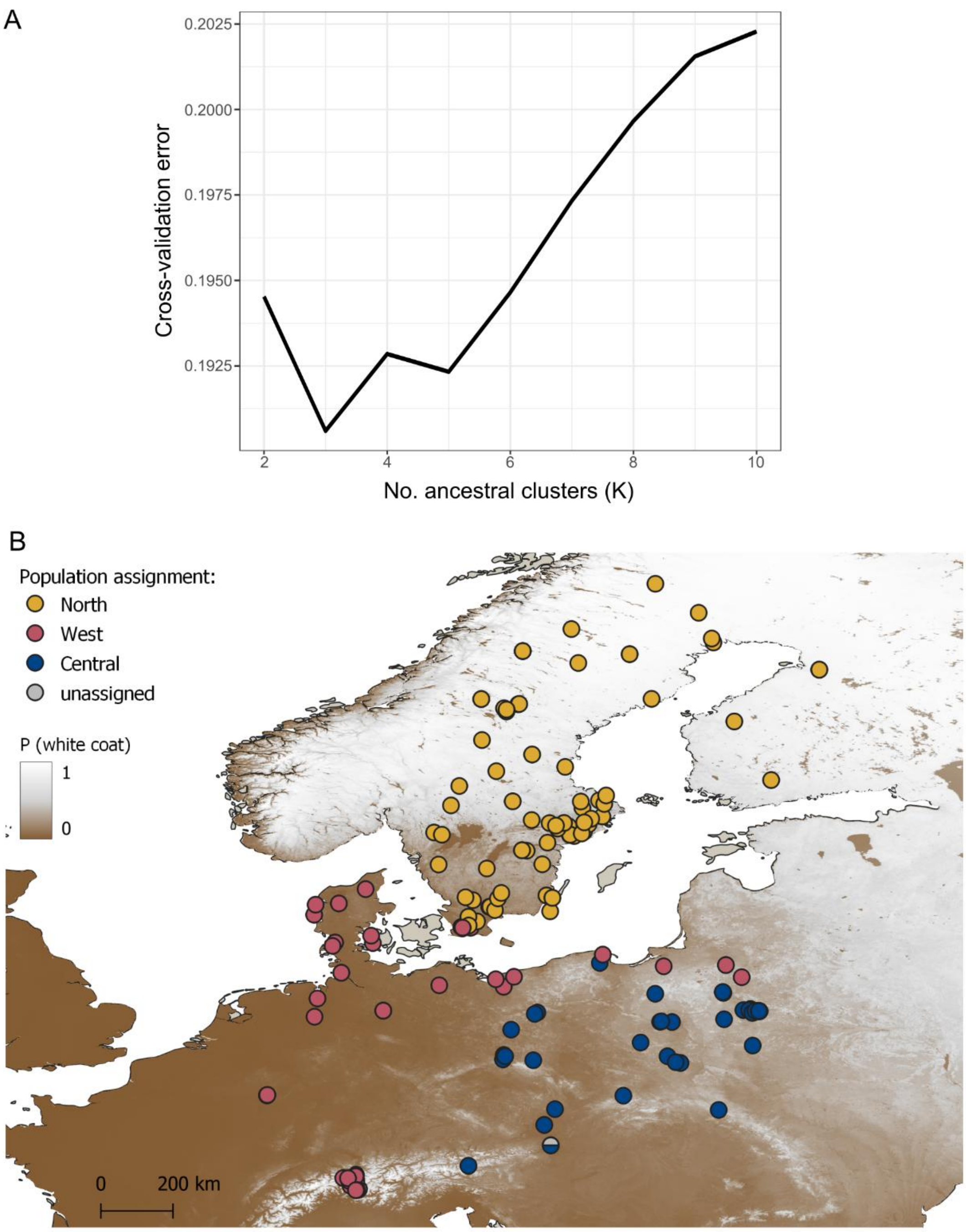


**Fig. S5 - Genome-wide population structure of European least weasel populations. (A)** Cross-validation error of the ADMIXTURE analysis indicates K = 3 as the best number of ancestral clusters. **(B)** Sampling localities of European least weasels used in this study, colored by population genetic assignment, estimated from the ADMIXTURE analysis (Fig. 3C), considering the best K value. Overlapping localities are indicated by a single symbol. Sampling is overlaid in the clinal distribution of winter colors, shown as the probability of winter-white coats across the European distribution range of the species (adapted from ref.[14]; data available at ref.[15]).

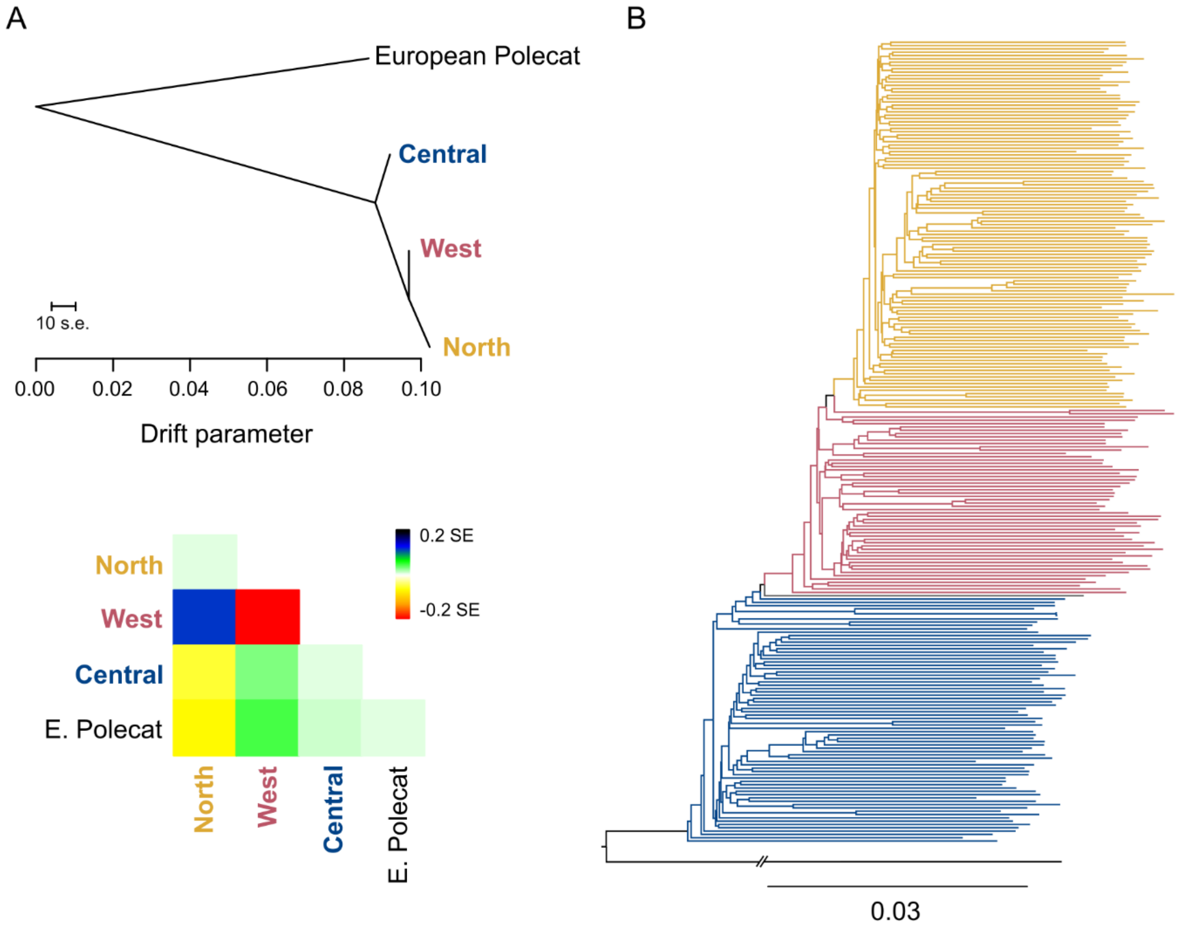


**Fig. S6 - Evolutionary relationships of European least weasel populations. (A)** TreeMix population tree (top), derived from allele frequencies estimated for 8,141 genome-wide SNPs, for the three genetic populations inferred from ADMIXTURE (Fig. 3C; Supplementary Fig. S5). Three European polecats (*M. putorius*) were used as the outgroup population. No migration edges were allowed. Residuals for the resulting tree estimate (bottom) are also shown. **(B)** Neighbor-joining tree of European least weasels, reconstructed from individual pairwise genetic distances (N = 242 individuals) estimated based on 8,141 SNPs. One European polecat (*Mustela putorius*) was used as the outgroup. Individuals are colored according to population assignment in ADMIXTURE (Fig. 3C).

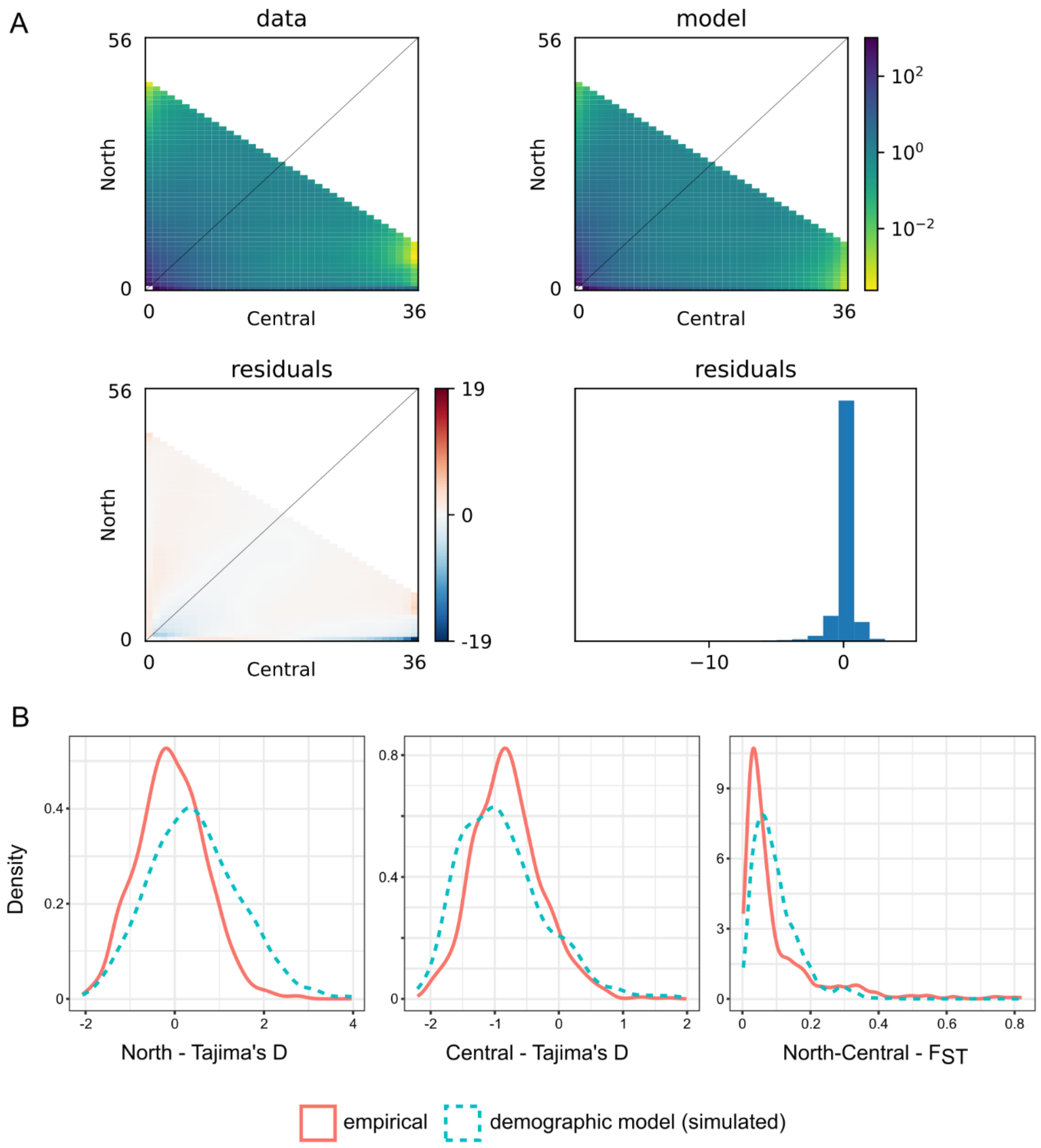


**Fig. S7 - Validations of the demographic model of North and Central European least weasel populations. (A)** The empirical joint site frequency spectrum (jSFS) between North and Central populations, down-projected to 25% of the original sampling size (top-left), the inferred jSFS under the best demographic scenario selected with GADMA (top-right), their comparison (bottom-left), and the distribution of residual values (bottom-right). **(B)** Comparison of the empirical and predicted distributions of Tajima's D for North (left) and Central (middle) populations, and pairwise $F_{ST}$ estimates between populations (right). Empirical distributions were estimated from 1,000 anonymous 2 kb regions, and predicted distributions were simulated based on the best demographic scenario for the population pair.

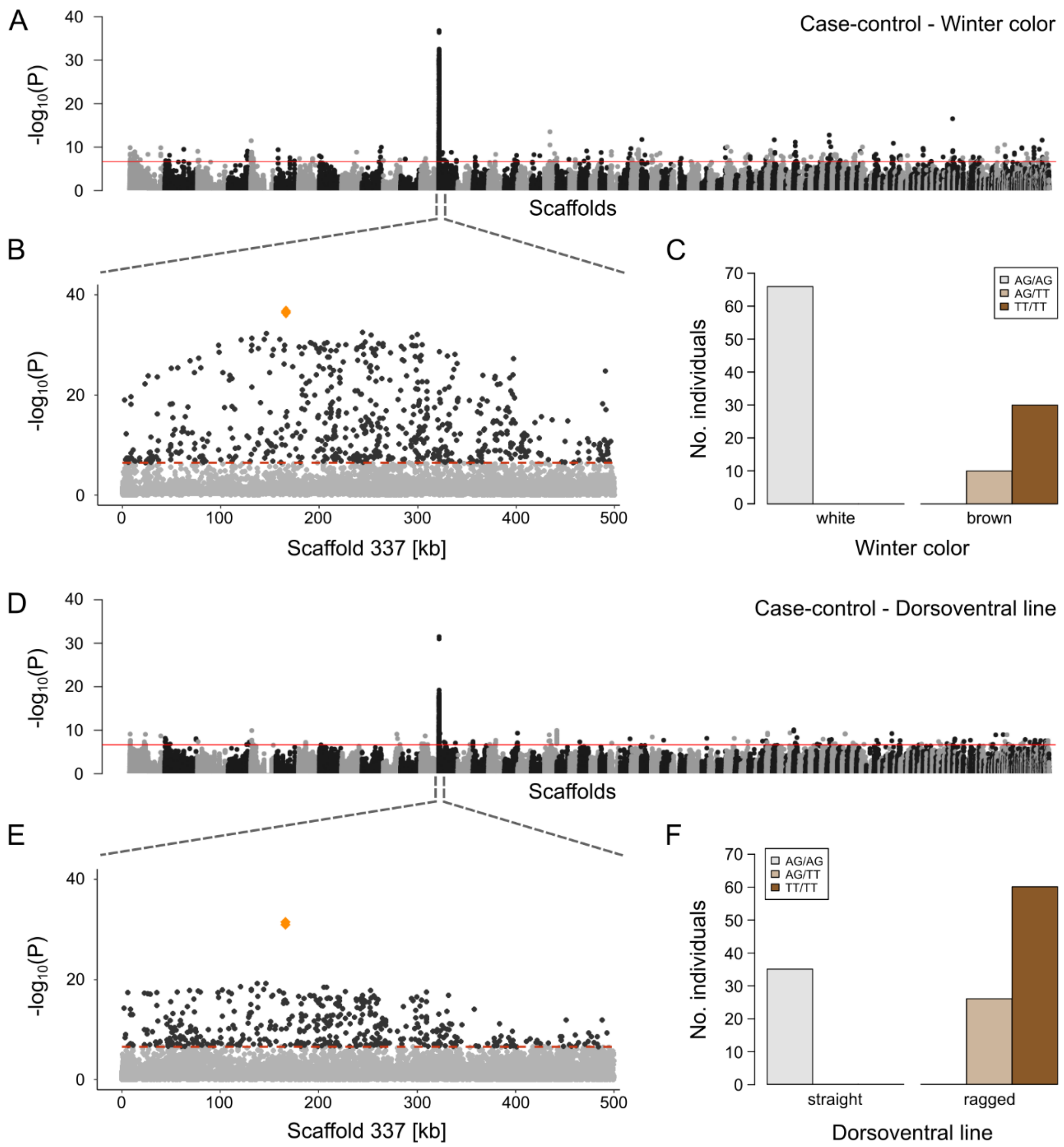


**Fig. S8 - Genotype-to-phenotype mapping of coloration traits. (A)** Case-control test of allele frequency differences between specimens with alternative winter colors (white, N = 66; brown, N = 40; Supplementary Data S2), based on 156,141 SNPs across all captured regions. **(B)** Zoom-in of the strongest association peak, located in scaffold 337 of the *M. nivalis* draft reference genome. Orange diamonds identify the two consecutive SNPs in the *MC1R* coding sequence. **(C)** Genotypes per phenotype at the two *MC1R* SNPs, for all specimens used in (A). **(D)** Case-control test of allele frequency differences between specimens with alternative dorsoventral lines (straight, N = 35; ragged, N = 86; Supplementary Data S2), based on 204,777 SNPs across all captured regions. **(E)** Zoom-in of the strongest association peak, located in scaffold 337. Orange diamonds identify the two candidate SNPs in the *MC1R* coding sequence. **(F)** Genotypes per phenotype at the two candidate SNPs, for all specimens used in (D). In both scans, scaffolds are ordered from left to right by decreasing length. Red lines are the Bonferroni-corrected threshold of P = 0.05.

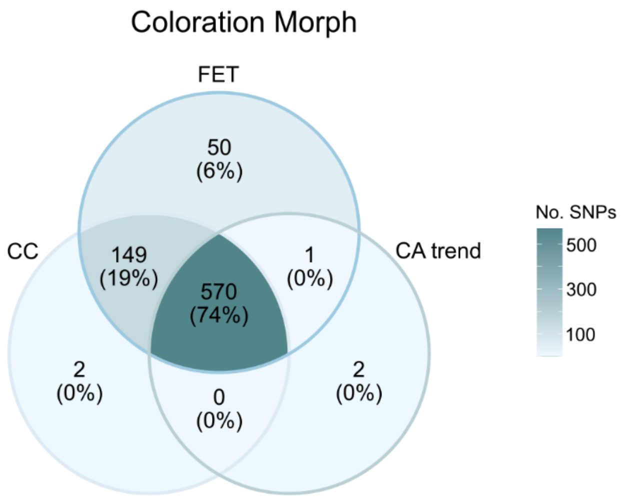


**Fig. S9 - Genotype-to-phenotype mapping outliers.** Venn diagram of common outlier SNPs in three genotype-to-phenotype association tests – case-control (CC), Fisher's exact test (FET), and Cochran-Armitage trend test (CA trend) – for alternative coloration morphs (*nivalis*, N = 101; *vulgaris*, N = 88).

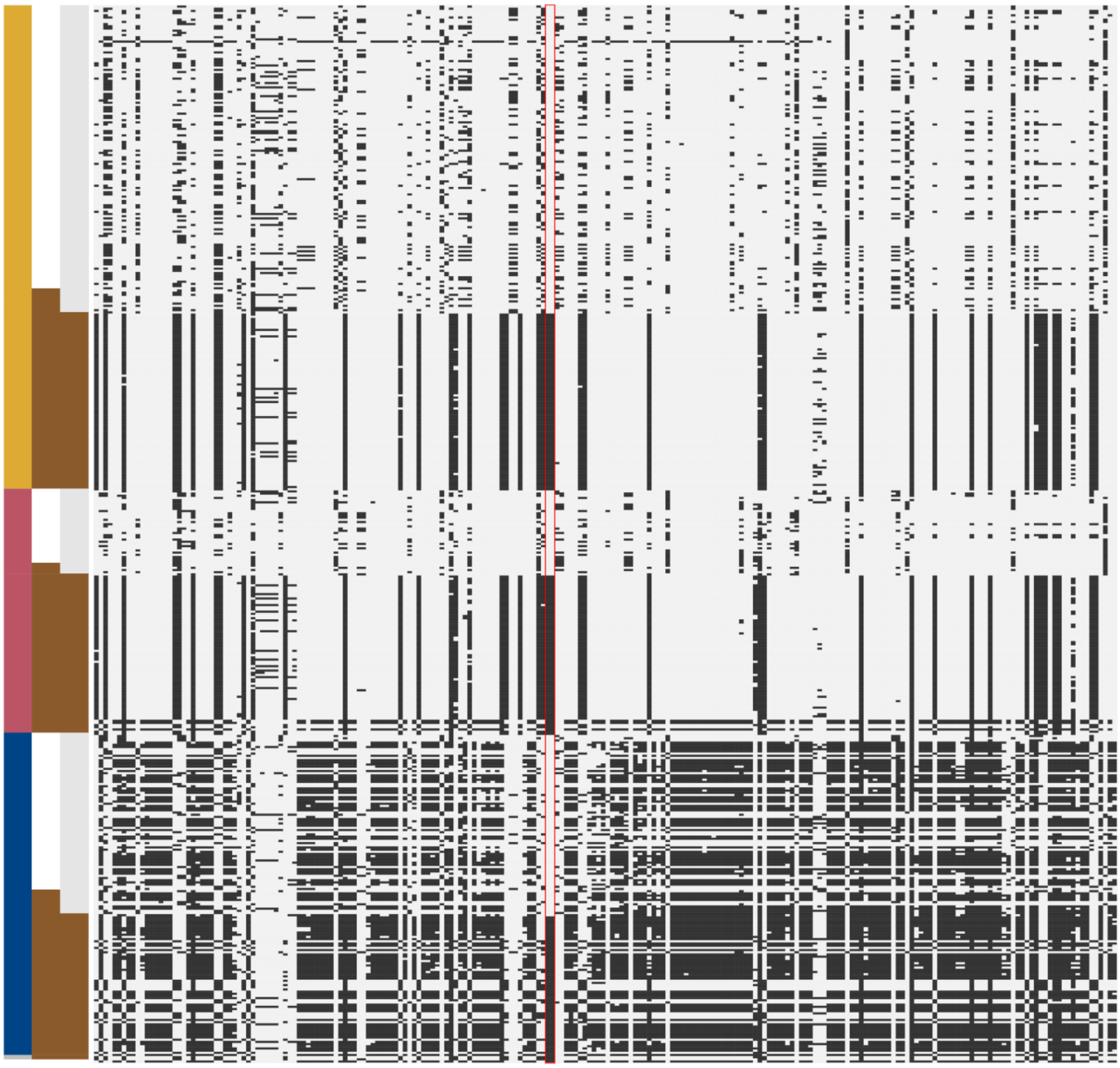

**Fig. S10 - Haplotypes of European least weasels in the region surrounding the *MC1R* gene.** Phased *Mustela nivalis* haplotypes, spanning a 40 kb region centered on *MC1R* variants (scaffold 337:145,924–185,924). Each row depicts a haplotype, and each column represents a variant locus. Alleles are colored as follows: grey – predominant *nivalis* allele; black – alternative allele. The red box highlights the two SNPs in the *MC1R* coding sequence. Colored columns on the left side of the plot represent, from left to right, the population of origin of the haplotypes (yellow – North, pink – West, blue – Central; grey - unassigned), the coloration morph of the specimen (white – *nivalis*, brown – *vulgaris*), and the allele at *MC1R* variants (grey – *nivalis* allele, brown – *vulgaris* allele).

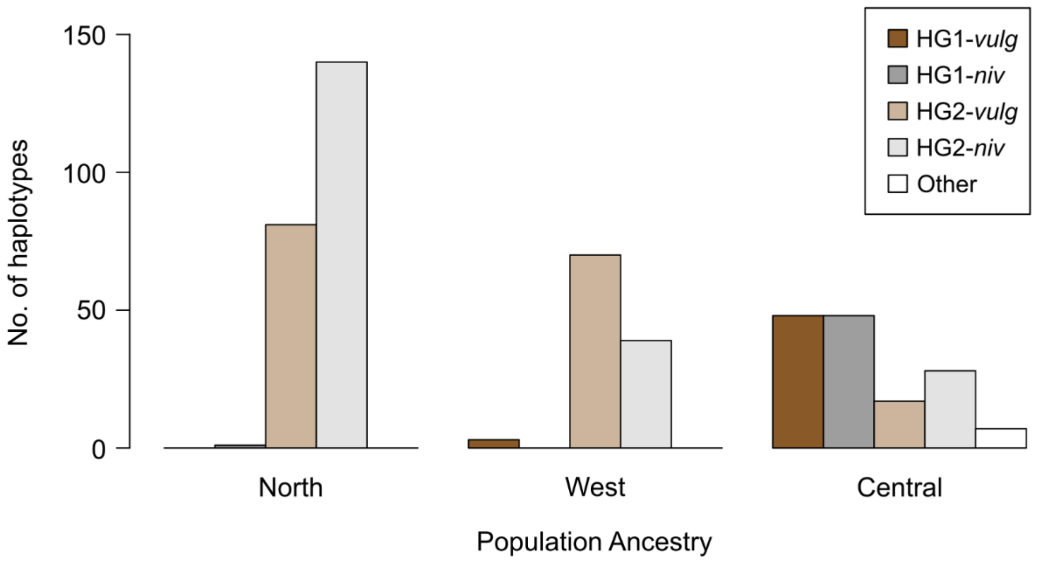


**Fig. S11 - *MC1R* haplogroup prevalence in European least weasel populations.** Counts of haplotypes present in the three ancestry groups identified in our dataset, segregated by both haplogroup (following the haplotypic structure identified in the median-joining network, Fig. 4D) and *MC1R* variants: HG1-*vulg* (haplogroup 1, *vulgaris* allele), HG1-*niv* (haplogroup 1, *nivalis* allele), HG2-*vulg* (haplogroup 2, *vulgaris* allele), HG2-*niv* (haplogroup 2, *nivalis* allele), and Other (intermediate haplotypes in the haplotype network).

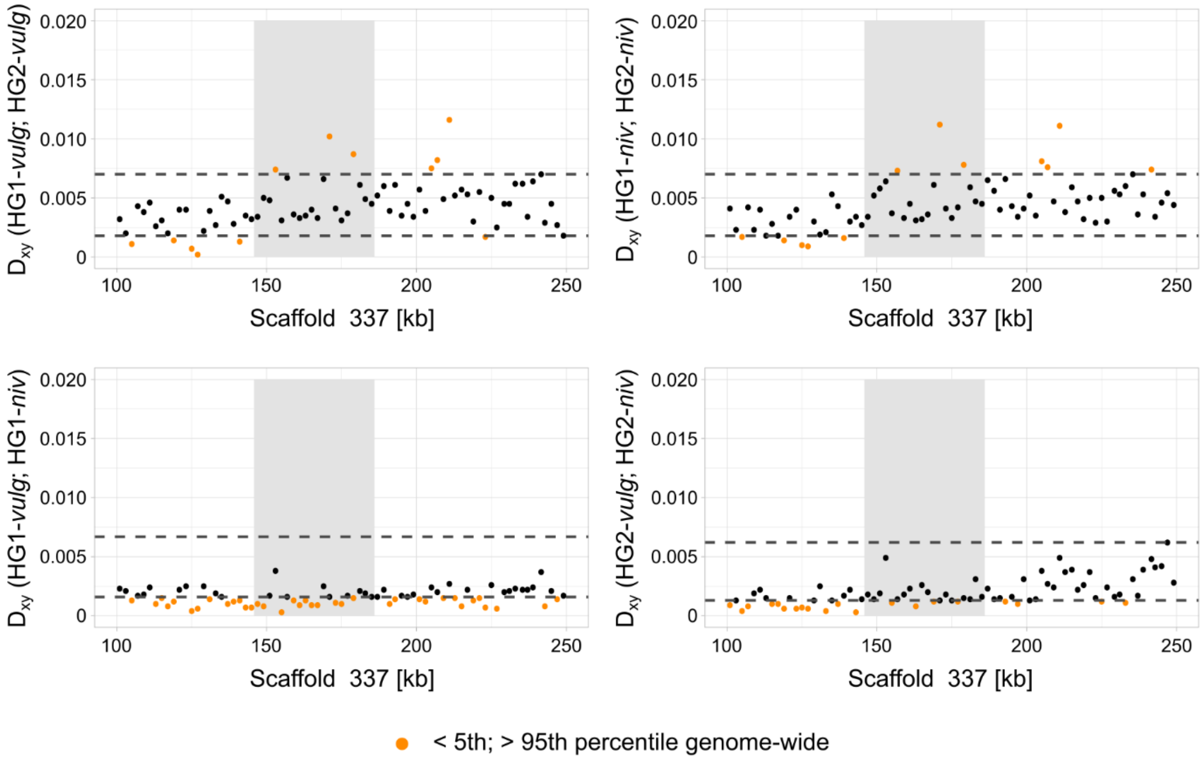


**Fig. S12 - Genetic divergence estimates between European least weasels**. Estimates of absolute genetic divergence ($D_{xy}$) between pairs of distinct *MC1R* haplogroup-allele datasets (haplogroup 1 – HG1; haplogroup 2 – HG2; *nivalis* allele – *niv*; *vulgaris* allele – *vulg*). Each dataset is comprised only of specimens homozygous for each haplogroup-allele combination (HG1-*vulg*, N = 15; HG1-*niv*, N = 15; HG2-*vulg*, N = 71; HG2-*niv*, N = 88). Plots show the average estimates of $D_{xy}$ along the *MC1R* capture target (scaffold 337:100,000–250,000), where each dot represents a 2 kb non-overlapping window. Dashed black lines represent the 5th and 95th percentile of the distribution of genome-wide estimates (based on 1,000 anonymous 2 kb capture fragments), and orange dots depict *MC1R* estimates that fall outside the 5th to 95th genome-wide percentile range. The grey-shaded area shows a 40 kb region centered on *MC1R* variants, where extended haplotype homozygosity decays to approximately zero (see Fig. 4C).

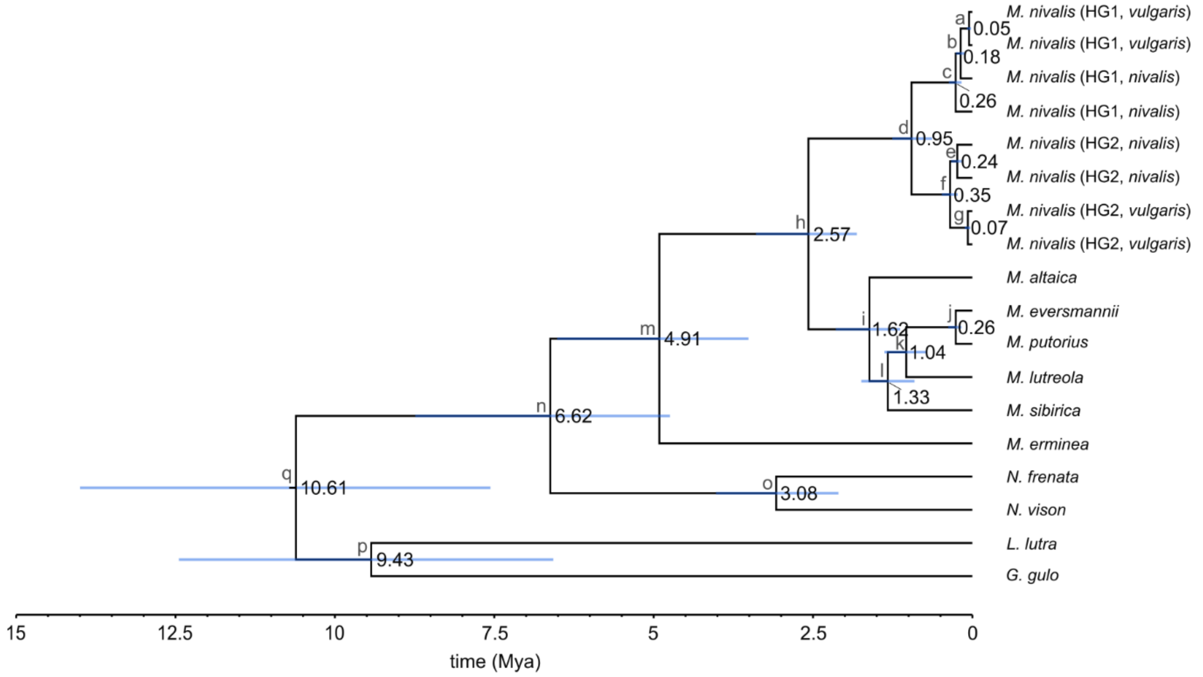


**Fig. S13 - Divergence time of *MC1R* haplotypes in Mustelidae species.** Bayesian inference tree of divergence time estimates between haplotypes of *Mustela nivalis* and other mustelid species, for a 40 kb region centered on *MC1R* variants (scaffold 337:145,924–185,924). Divergence time point estimates (millions of years; Mya) are shown next to nodes, and 95% high probability density intervals (HPD; here shown as blue bars) are provided in Supplementary Table S8. Phased haplotypes for four *M. nivalis* specimens, homozygous for each *MC1R* haplogroup-allele combination, and one randomly phased sequence for each of the remaining mustelid species are included.

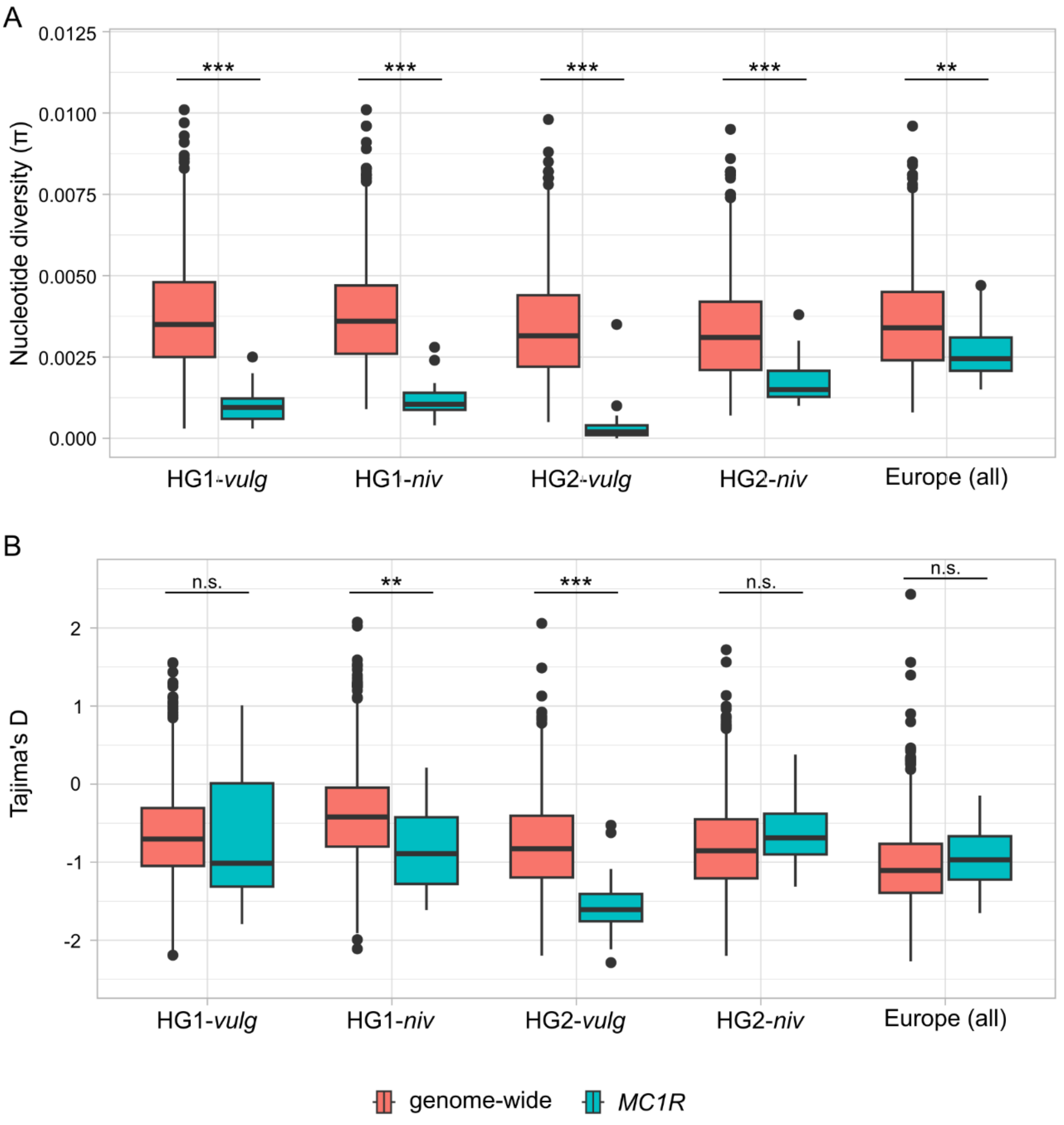


**Fig. S14 - Genetic diversity estimates within European least weasels.** Estimates of **(A)** nucleotide diversity (π) and **(B)** Tajima's D, for datasets comprised of homozygous specimens for each *MC1R* haplogroup-allele combination (haplogroup 1 – HG1; haplogroup 2 – HG2; *nivalis* allele – *niv*; *vulgaris* allele – *vulg*; HG1-*vulg*, N = 15; HG1-*niv*, N = 15; HG2-*vulg*, N = 71; HG2-*niv*, N = 88) and the complete European dataset (N = 242). Estimates were calculated for two distinct capture fragment sets: i) 1,000 anonymous genome-wide 2 kb capture fragments, and ii) a 40 kb region centered on *MC1R* variants (scaffold 337:145,924-185,924), split in 2 kb non-overlapping windows. Significant differences between estimates for genome-wide and *MC1R* windows in each sample set were assessed with a Kruskal-Wallis test and are denoted in each panel as: n.s. – $P > 0.05$; * – $P < 0.05$; ** – $P < 0.01$; *** – $P < 0.001$.

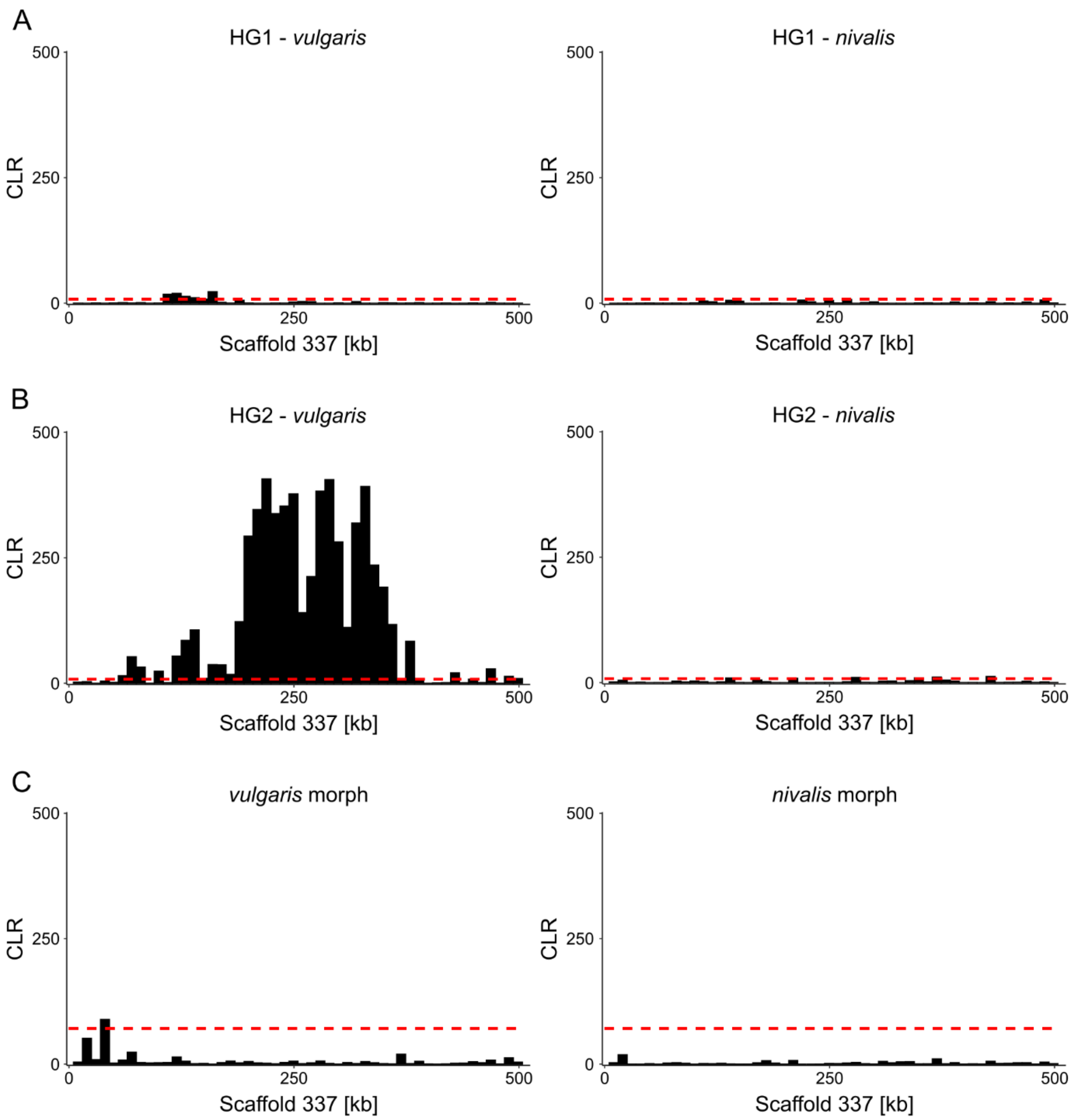


**Fig. S15 - Inference of selective sweep signatures in European least weasels.** SweeD composite likelihood ratio (CLR) estimates along the *MC1R* region capture target (scaffold 337:1–500,000). Estimates were conducted for **(A)** homozygous specimens for haplogroup 1 (HG1), carrying the *vulgaris* (left; N = 15) and *nivalis* (right: N = 15) allele; **(B)** homozygous specimens for haplogroup 2 (HG2), carrying the *vulgaris* (left; N = 71) and *nivalis* (right; N = 88) allele; and **(C)** all specimens assigned to the *vulgaris* (left; N = 124) and *nivalis* (right; N = 118) coloration morphs. The red dashed lines show the 95th percentile of the empirical distribution of CLR values estimated along scaffold 337.

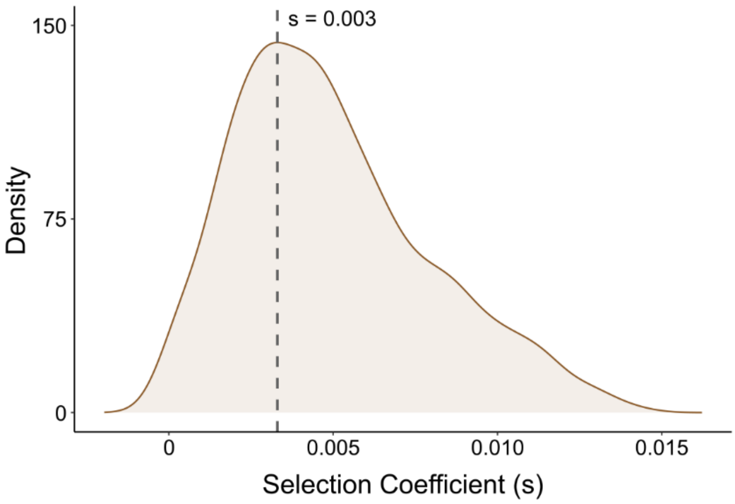


**Fig. S16 - Selection coefficient estimate for the HG2-*vulgaris* haplotype selective sweep.** Inferred probability distribution for the selection coefficient (s) acting on least weasel specimens harboring the HG2-*vulgaris* allele combination at *MC1R* haplotypes. The dashed line identifies the highest probability value of s.

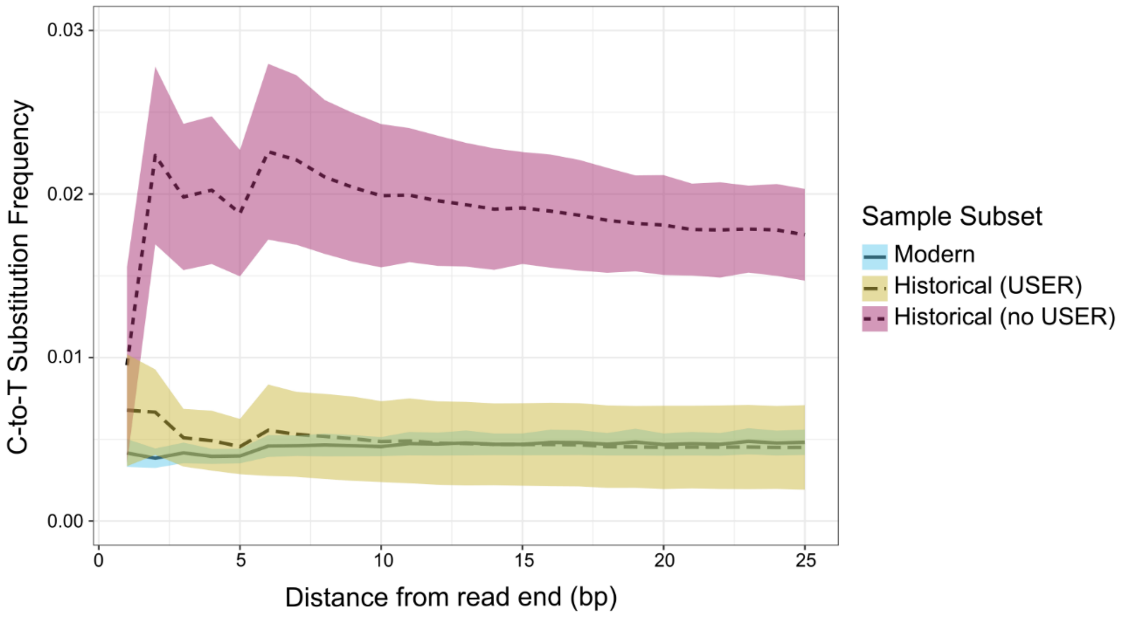


**Fig. S17 - DNA damage estimates in the complete sampling dataset used for capture assays.** Frequency of C-to-T substitutions by distance from the read end, for three distinct sample subsets (see details in Methods): modern samples (no USER treatment, N = 80), USER-treated historical samples (N = 187), and not-USER-treated historical samples (N = 4). Estimates per sample were obtained with MapDamage. The plot shows estimates averaged per sample subset: lines depict mean values, and shaded areas represent the standard deviation. USER-treated historical samples show a successful removal of most post-mortem damage patterns.

**Table S1 - Non-linear regression (f= a/(1+exp(-(x-x0)/b))) fit report for *in vitro* cellular assays of MC1R constructs.** The significance of P-values is indicated as follows: n.s. – non-significant; * – P < 0.05; ** – P < 0.01; *** – P < 0.001.

| **Ligand** | **MC1R Construct** | $R^2$ | **F** | **P-value** | | **Normality Test (Shapiro-Wilk)** | **Equal Variance Test (Levene)** |
|---|---|---|---|---|---|---|---|
| | | | | **P** | **Signif.** | | |
| **α-MSH** | **blank** | 0.9316 | 34.0514 | 0.0012 | ** | Passed (P = 0.5295) | Passed (P = 0.5779) |
| | **wild-type** | 0.9389 | 38.4155 | 0.0009 | *** | Passed (P = 0.7198) | Passed (P = 0.1388) |
| | **E92K** | 0.7029 | 5.9146 | 0.0481 | * | Passed (P = 0.9305) | Passed (P = 0.0474) |
| | **L99K** | 0.8359 | 12.7317 | 0.0109 | * | Passed (P = 0.9790) | Passed (P = 0.1823) |
| **ASIP** | **blank** | 0.3329 | 0.9979 | 0.4451 | n.s. | Passed (P = 0.4764) | Passed (P = 0.1209) |
| | **wild-type** | 0.8361 | 10.2048 | 0.0269 | * | Passed (P = 0.0418) | Passed (P = 0.4907) |
| | **E92K** | 0.4060 | 1.3670 | 0.3528 | n.s. | Passed (P = 0.9450) | Passed (P = 0.1209) |
| | **L99K** | 0.6379 | 3.523 | 0.1311 | n.s. | Passed (P = 0.3211) | Passed (P = 0.2965) |

**Table S2 - One-way analysis of variance (ANOVA) per construct for *in vitro* MC1R assays.** **(A)** ANOVA results across conditions (ligand concentrations) per test construct. **(B)** Post-hoc pairwise multiple comparisons for significant ANOVA results, i.e., wild-type construct assays, using the Student-Newman-Keuls test. Significance of P-values as in Supplementary Table S1.

**(A)**

| MC1R Construct | Ligand | Normality Test (Shapiro-Wilk) | Equal Variance Test (Levene) | P-value | |
|---|---|---|---|---|---|
| | | | | P | Signif. |
| **wild-type** | **α-MSH** | Passed (P = 0.548) | Passed (P = 0.220) | <0.001 | *** |
| | **ASIP** | Passed (P = 0.098) | Passed (P = 0.703) | 0.007 | ** |
| **E92K** | **α-MSH** | Passed (P = 0.252) | Passed (P = 0.756) | 0.976 | n.s. |
| | **ASIP** | Passed (P = 0.845) | Passed (P = 0.346) | 0.696 | n.s. |
| **L99K** | **α-MSH** | Passed (P = 0.193) | Passed (P = 0.921) | 0.415 | n.s. |
| | **ASIP** | Passed (P = 0.070) | Passed (P = 0.857) | 0.673 | n.s. |

**(B)**

| Ligand | Conc. (M) Comparison | P-value | | Ligand | Conc. (M) Comparison | P-value | |
|---|---|---|---|---|---|---|---|
| | | P | Signif. | | | P | Signif. |
| **α-MSH** | **0 vs. 1e-12** | 0.7 | n.s. | **ASIP** | **0 vs. 1e-11** | 0.345 | n.s. |
| | **0 vs. 1e-11** | 0.716 | n.s. | | **0 vs. 1e-10** | 0.443 | n.s. |
| | **0 vs. 1e-10** | 0.077 | n.s. | | **0 vs. 1e-9** | 0.496 | n.s. |
| | **0 vs. 1e-9** | 0.003 | ** | | **0 vs. 1e-8** | 0.231 | n.s. |
| | **0 vs. 1e-8** | 0.003 | ** | | **0 vs. 1e-7** | 0.293 | n.s. |
| | **0 vs. 1e-7** | <0.001 | *** | | **0 vs. 2.5e-7** | 0.223 | n.s. |
| | **0 vs. 1e-6** | 0.014 | * | | **1e-11 vs. 1e-10** | 0.949 | n.s. |
| | **1e-12 vs. 1e-11** | 0.698 | n.s. | | **1e-11 vs. 1e-9** | 0.946 | n.s. |
| | **1e-12 vs. 1e-10** | 0.094 | n.s. | | **1e-11 vs. 1e-8** | 0.84 | n.s. |
| | **1e-12 vs. 1e-9** | 0.004 | ** | | **1e-11 vs. 1e-7** | 0.064 | n.s. |
| | **1e-12 vs. 1e-8** | 0.004 | ** | | **1e-11 vs. 2.5e-7** | 0.032 | * |
| | **1e-12 vs. 1e-7** | 0.001 | *** | | **1e-10 vs. 1e-9** | 0.824 | n.s. |
| | **1e-12 vs. 1e-6** | 0.02 | * | | **1e-10 vs. 1e-8** | 0.956 | n.s. |
| | **1e-11 vs. 1e-10** | 0.084 | n.s. | | **1e-10 vs. 1e -7** | 0.066 | n.s. |
| | **1e-11 vs. 1e-9** | 0.006 | ** | | **1e-10 vs. 2.5e-7** | 0.03 | * |
| | **1e-11 vs. 1e-8** | 0.006 | ** | | **1e-9 vs. 1e-8** | 0.963 | n.s. |
| | **1e-11 vs. 1e-7** | 0.002 | ** | | **1e-9 vs. 1e-7** | 0.105 | n.s. |
| | **1e-11 vs. 1e-6** | 0.025 | * | | **1e-9 vs. 2.5e-7** | 0.05 | * |
| | **1e-10 vs. 1e-9** | 0.148 | n.s. | | **1e-8 vs. 1e-7** | 0.057 | n.s. |
| | **1e-10 vs. 1e-8** | 0.13 | n.s. | | **1e-8 vs. 2.5e-7** | 0.032 | * |
| | **1e-10 vs. 1e-7** | 0.051 | n.s. | | **1e-7 vs. 2.5e-7** | 0.453 | n.s. |
| | **1e-10 vs. 1e-6** | 0.294 | n.s. | | | | |
| | **1e-9 vs. 1e-8** | 0.841 | n.s. | | | | |
| | **1e-9 vs. 1e-7** | 0.441 | n.s. | | | | |
| | **1e-9 vs. 1e-6** | 0.482 | n.s. | | | | |
| | **1e-8 vs. 1e-7** | 0.591 | n.s. | | | | |
| | **1e-8 vs. 1e-6** | 0.344 | n.s. | | | | |
| | **1e-7 vs. 1e-6** | 0.24 | n.s. | | | | |

**Table S3 - One-way analysis of variance (ANOVA) per α-MSH condition for *in vitro* MC1R assays. (A)** ANOVA results across test constructs per α-MSH condition (same concentration). **(B)** Post-hoc pairwise multiple comparisons for significant ANOVA results, using the Student-Newman-Keuls test. Significance of P-values as in Supplementary Table S1.

**(A)**

| α-MSH Conc. (M) | Normality Test (Shapiro-Wilk) | Equal Variance Test (Levene) | P-value | |
|---|---|---|---|---|
| | | | P | Signif. |
| **0** | Passed (P = 0.306) | Passed (P = 0.218) | 0.004 | ** |
| **1e-12** | Passed (P = 0.985) | Passed (P = 0.398) | <0.001 | *** |
| **1e-11** | Passed (P = 0.511) | Failed (P < 0.050) | n/a | n/a |
| **1e-10** | Passed (P = 0.622) | Passed (P = 0.603) | <0.001 | *** |
| **1e-9** | Passed (P = 0.570) | Passed (P = 0.246) | 0.001 | *** |
| **1e-8** | Passed (P = 0.476) | Passed (P = 0.340) | <0.001 | *** |
| **1e-7** | Passed (P = 0.397) | Passed (P = 0.434) | <0.001 | *** |
| **1e-6** | Passed (P = 0.609) | Passed (P = 0.249) | 0.002 | ** |

**(B)**

| α-MSH Conc. (M) | Construct Comparison | P-value | |
|---|---|---|---|
| | | P | Signif. |
| **0** | **Blank vs. WT** | 0.02 | * |
| | **Blank vs. E92K** | 0.003 | ** |
| | **Blank vs. L99K** | 0.007 | ** |
| | **WT vs. E92K** | 0.092 | n.s |
| | **WT vs. L99K** | 0.216 | n.s. |
| | **E92K vs. L99K** | 0.305 | n.s. |
| **1e-12** | **Blank vs. WT** | 0.003 | ** |
| | **Blank vs. E92K** | <0.001 | *** |
| | **Blank vs. L99K** | 0.001 | *** |
| | **WT vs. E92K** | 0.039 | * |
| | **WT vs. L99K** | 0.141 | n.s. |
| | **E92K vs. L99K** | 0.202 | n.s. |
| **1e-10** | **Blank vs. WT** | <0.001 | *** |
| | **Blank vs. E92K** | 0.001 | *** |
| | **Blank vs. L99K** | 0.001 | *** |
| | **WT vs. E92K** | 0.174 | n.s. |
| | **WT vs. L99K** | 0.148 | n.s. |
| | **E92K vs. L99K** | 0.55 | n.s. |
| **1e-9** | **Blank vs. WT** | <0.001 | *** |
| | **Blank vs. E92K** | 0.011 | * |
| | **Blank vs. L99K** | 0.006 | ** |
| | **WT vs. E92K** | 0.022 | * |
| | **WT vs. L99K** | 0.04 | * |
| | **E92K vs. L99K** | 0.858 | n.s. |

| αMSH Conc. (M) | Construct Comparison | P-value | |
|---|---|---|---|
| | | P | Signif. |
| **1e-8** | **Blank vs. WT** | <0.001 | *** |
| | **Blank vs. E92K** | 0.002 | ** |
| | **Blank vs. L99K** | 0.002 | ** |
| | **WT vs. E92K** | 0.023 | * |
| | **WT vs. L99K** | 0.03 | * |
| | **E92K vs. L99K** | 0.481 | n.s. |
| **1e-7** | **Blank vs. WT** | <0.001 | *** |
| | **Blank vs. E92K** | 0.005 | ** |
| | **Blank vs. L99K** | 0.004 | ** |
| | **WT vs. E92K** | 0.039 | * |
| | **WT vs. L99K** | 0.061 | n.s. |
| | **E92K vs. L99K** | 0.419 | n.s. |
| **1e-6** | **Blank vs. WT** | 0.002 | ** |
| | **Blank vs. E92K** | 0.005 | ** |
| | **Blank vs. L99K** | 0.003 | ** |
| | **WT vs. E92K** | 0.303 | n.s. |
| | **WT vs. L99K** | 0.404 | n.s. |
| | **E92K vs. L99K** | 0.801 | n.s. |

**Table S4 - One-way analysis of variance (ANOVA) per ASIP condition for *in vitro* MC1R assays. (A)** ANOVA results across test constructs per ASIP condition (same concentration). **(B)** Post-hoc pairwise multiple comparisons for significant ANOVA results, using the Student-Newman-Keuls test. Significance of P-values as in Supplementary Table S1.

**(A)**

| ASIP Conc. (M) | Normality Test (Shapiro-Wilk) | Equal Variance Test (Levene) | P-value | |
|---|---|---|---|---|
| | | | P | Signif. |
| **0** | Failed (P < 0.050) | Passed (P = 0.082) | <0.001 | *** |
| **1e-11** | Passed (P = 0.119) | Passed (P = 0.057) | <0.001 | *** |
| **1e-10** | Failed (P < 0.050) | Passed (P = 0.254) | <0.001 | *** |
| **1e-9** | Passed (P = 0.818) | Failed (P < 0.050) | n/a | n/a |
| **1e-8** | Passed (P = 0.288) | Passed (P = 0.054) | <0.001 | *** |
| **1e-7** | Passed (P = 0.489) | Failed (P < 0.050) | n/a | n/a |
| **2.5e-7** | Passed (P = 0.091) | Passed (P = 0.065) | <0.001 | *** |

**(B)**

| ASIP Conc. (M) | Construct Comparison | P-value | |
|---|---|---|---|
| | | P | Signif. |
| **0** | **Blank vs. WT** | <0.001 | *** |
| | **Blank vs. E92K** | <0.001 | *** |
| | **Blank vs. L99K** | <0.001 | *** |
| | **WT vs. E92K** | 0.084 | n.s. |
| | **WT vs. L99K** | 0.061 | n.s. |
| | **E92K vs. L99K** | 0.78 | n.s. |
| **1e-11** | **Blank vs. WT** | <0.001 | *** |
| | **Blank vs. E92K** | <0.001 | *** |
| | **Blank vs. L99K** | <0.001 | *** |
| | **WT vs. E92K** | 0.415 | n.s. |
| | **WT vs. L99K** | 0.193 | n.s. |
| | **E92K vs. L99K** | 0.342 | n.s. |
| **1e-10** | **Blank vs. WT** | <0.001 | *** |
| | **Blank vs. E92K** | <0.001 | *** |
| | **Blank vs. L99K** | <0.001 | *** |
| | **WT vs. E92K** | 0.198 | n.s. |
| | **WT vs. L99K** | 0.096 | n.s. |
| | **E92K vs. L99K** | 0.396 | n.s. |
| **1e-8** | **Blank vs. WT** | <0.001 | *** |
| | **Blank vs. E92K** | <0.001 | *** |
| | **Blank vs. L99K** | <0.001 | *** |
| | **WT vs. E92K** | 0.198 | n.s. |
| | **WT vs. L99K** | 0.096 | n.s. |
| | **E92K vs. L99K** | 0.396 | n.s. |
| **2.5e-7** | **Blank vs. WT** | 0.053 | n.s. |
| | **Blank vs. E92K** | 0.001 | *** |
| | **Blank vs. L99K** | <0.001 | *** |
| | **WT vs. E92K** | 0.018 | * |
| | **WT vs. L99K** | 0.009 | ** |
| | **E92K vs. L99K** | 0.738 | n.s. |

**Table S5 - Genotypes at *MC1R* SNPs for Mustelidae samples.** For each sample, information is given about *a priori* phenotyped winter color (if applicable) and genotypes at causal *MC1R* SNPs. Sample origin (museum collection or public dataset) is indicated in Supplementary Data S3 and at the end of this table.

| Species | Sample Code[1] | Winter color | Genotype at *MC1R* SNPs | |
|---|---|---|---|---|
| | | | 337:165,924 | 337:165,925 |
| *Mustela altaica* | NRM-MA580177 | n/a | A/A | G/G |
| *Mustela altaica* | NRM-MA580196 | n/a | A/A | G/G |
| *Mustela altaica* | NRM-MA590332 | n/a | A/A | G/G |
| *Mustela erminea* | ZFMK-MAM-1939.0129f | white | A/A | G/G |
| *Mustela erminea* | ZFMK-MAM-2007.0242 | brown | A/A | G/G |
| *Mustela erminea* | ZFMK-MAM-2012.0169 | white | A/A | G/G |
| *Mustela erminea* | NH:1985.79.17 | brown | A/A | G/G |
| *Mustela eversmannii* | NRM-MA581155 | n/a | A/A | G/G |
| *Mustela eversmannii* | NRM-MA581156 | n/a | A/A | G/G |
| *Mustela eversmannii* | NRM-MA581781 | n/a | A/A | G/G |
| *Mustela kathiah* | NRM-MA601747 | n/a | A/A | G/G |
| *Mustela lutreola* | NRM-MA640242 | n/a | A/A | G/G |
| *Mustela nigripes* | Cheerio SB6573 [(a)] | n/a | A/A | G/G |
| *Mustela nigripes* | 85 W2094 [(a)] | n/a | A/A | G/G |
| *Mustela nigripes* | Willa SB10 [(a)] | n/a | A/A | G/G |
| *Mustela putorius* | NRM-MA595348 | n/a | A/A | G/G |
| *Mustela putorius* | NRM-MA605346 | n/a | A/A | G/G |
| *Mustela putorius* | NRM-MA825003 | n/a | A/A | G/G |
| *Mustela sibirica* | NRM-MA581732 | n/a | A/A | G/G |
| *Mustela sibirica* | NRM-MA600662 | n/a | A/A | G/G |
| *Neogale frenata* | MCZ:Mamm:58767 | white | A/A | G/G |
| *Neogale frenata* | MCZ:Mamm:65941 | brown | A/A | G/G |
| *Neogale frenata* | USNM222988 | white | A/A | G/G |
| *Neogale frenata* | USNM261844 | brown | N/N | N/N |
| *Neogale vison* | NRM-MA590062 | n/a | A/A | G/G |
| *Neogale vison* | NRM-MA875248 | n/a | A/A | G/G |
| *Neogale vison* | NRM-MA875249 | n/a | A/A | G/G |
| *Gulo gulo* | Ind743 [(a)] | n/a | A/A | G/G |
| *Lutra lutra* | mLutLut1 [(a)] | n/a | A/A | G/G |

[1] Museum/Research Collections:

MCZ - Museum of Comparative Zoology at Harvard University
NMI - National Museum of Ireland
NRM - Swedish Museum of Natural History
USNM - Smithsonian National Museum of Natural History
ZFMK - Zoological Research Museum Alexander Koenig
(a) Samples obtained from public datasets; see details in Supplementary Data S3.

**Table S6 - Genotypes at *MC1*R SNPs for least weasel samples.** For each sample, information is given about *a priori* phenotyped winter color, dorsoventral line type, and color morph, and genotypes at causal *MC1R* SNPs. For samples with *a priori* uncertain color morph, SNP-informed morph is also indicated. Sample origin (museum or research collection) is indicated in Supplementary Data S1 and at the end of this table.

| Sample Code[1] | Winter color | Line Type | Color morph | Genotypes at *MC1R* SNPs | | SNP-informed coloration morph |
|---|---|---|---|---|---|---|
| | | | | 337:165,924 | 337:165,925 | |
| AMNH232206 | brown | ragged | n/a | A/A | G/G | n/a |
| AMNH31669 | white | uncertain | n/a | A/A | G/G | n/a |
| AMNH99436 | brown | ragged | n/a | A/A | G/G | n/a |
| AMNH99438 | brown | ragged | n/a | A/A | G/G | n/a |
| AMNH99439 | white | uncertain | n/a | A/A | G/G | n/a |
| BNM12280 | uncertain | ragged | *c* | T/T | T/T | n/a |
| BNM12281 | uncertain | ragged | *vulgaris* | T/T | T/T | n/a |
| BNM12282 | uncertain | ragged | *vulgaris* | T/T | T/T | n/a |
| BNM12284 | uncertain | ragged | *vulgaris* | T/T | T/T | n/a |
| BNM12286 | uncertain | ragged | *vulgaris* | T/T | T/T | n/a |
| BNM12287 | uncertain | ragged | *vulgaris* | T/T | T/T | n/a |
| BNM12692 | uncertain | straight | *nivalis* | A/A | G/G | n/a |
| BNM12740 | white | uncertain | *nivalis* | A/A | G/G | n/a |
| BNM13143 | uncertain | ragged | *vulgaris* | T/T | T/T | n/a |
| BNM13145 | uncertain | straight | *nivalis* | A/A | G/G | n/a |
| BNM13250 | uncertain | straight | *nivalis* | A/A | G/G | n/a |
| BNM13363 | uncertain | straight | *nivalis* | A/A | G/G | n/a |
| BNM13735 | uncertain | straight | *nivalis* | A/A | G/G | n/a |
| BNM16656 | uncertain | straight | *nivalis* | A/A | G/G | n/a |
| BNM16888 | white | uncertain | *nivalis* | A/A | G/G | n/a |
| BNM17141 | molting | uncertain | *nivalis* | A/A | G/G | n/a |
| BNM17183 | uncertain | straight | *nivalis* | A/A | G/G | n/a |
| BNM17184 | uncertain | straight | *nivalis* | A/A | G/G | n/a |
| BNM17185 | uncertain | ragged | *vulgaris* | A/T | G/T | n/a |
| BNM17186 | uncertain | ragged | *vulgaris* | A/T | G/T | n/a |
| BNM17187 | uncertain | straight | *nivalis* | A/A | G/G | n/a |
| BNM17190 | uncertain | ragged | *vulgaris* | A/T | G/T | n/a |
| BNM17192 | uncertain | straight | *nivalis* | A/A | G/G | n/a |
| BNMV | uncertain | ragged | *vulgaris* | A/T | G/T | n/a |
| SB1177 | uncertain | straight | *nivalis* | A/A | G/G | n/a |
| SB1180 | uncertain | ragged | *vulgaris* | A/T | G/T | n/a |
| CE3C112 | uncertain | uncertain | uncertain | A/A | G/G | *nivalis* |
| CE3C113 | uncertain | uncertain | uncertain | A/A | G/G | *nivalis* |
| CE3C114 | uncertain | uncertain | uncertain | A/A | G/G | *nivalis* |
| CE3C116 | uncertain | uncertain | uncertain | A/A | G/G | *nivalis* |
| CE3C118 | uncertain | uncertain | uncertain | A/A | G/G | *nivalis* |

| | | | | | | |
|---|---|---|---|---|---|---|
| CE3C120 | uncertain | uncertain | uncertain | A/A | G/G | *nivalis* |
| CE3C121 | uncertain | uncertain | uncertain | A/A | G/G | *nivalis* |
| CE3C122 | uncertain | uncertain | uncertain | A/A | G/G | *nivalis* |
| CE3C123 | uncertain | uncertain | uncertain | A/A | G/G | *nivalis* |
| CE3C16 | uncertain | uncertain | uncertain | T/T | T/T | *vulgaris* |
| CE3C160 | uncertain | uncertain | uncertain | T/T | T/T | *vulgaris* |
| CE3C161 | uncertain | uncertain | uncertain | A/A | G/G | *nivalis* |
| CE3C17 | uncertain | uncertain | uncertain | T/T | T/T | *vulgaris* |
| CE3C171 | uncertain | uncertain | uncertain | A/A | G/G | *nivalis* |
| CE3C24 | uncertain | uncertain | uncertain | T/T | T/T | *vulgaris* |
| CE3C303 | uncertain | uncertain | uncertain | A/A | G/G | *nivalis* |
| CE3C305 | uncertain | uncertain | uncertain | A/A | G/G | *nivalis* |
| CE3C33 | uncertain | uncertain | uncertain | T/T | T/T | *vulgaris* |
| CE3C35 | uncertain | uncertain | uncertain | T/T | T/T | *vulgaris* |
| CE3C43 | uncertain | uncertain | uncertain | T/T | T/T | *vulgaris* |
| CE3C44 | uncertain | uncertain | uncertain | T/T | T/T | *vulgaris* |
| CE3C45 | uncertain | uncertain | uncertain | T/T | T/T | *vulgaris* |
| CE3C47 | uncertain | uncertain | uncertain | T/T | T/T | *vulgaris* |
| CE3C50 | uncertain | uncertain | uncertain | T/T | T/T | *vulgaris* |
| CE3C81 | uncertain | uncertain | uncertain | T/T | T/T | *vulgaris* |
| CE3C82 | uncertain | uncertain | uncertain | T/T | T/T | *vulgaris* |
| CE3C83 | uncertain | uncertain | uncertain | T/T | T/T | *vulgaris* |
| CE3C84 | uncertain | uncertain | uncertain | T/T | T/T | *vulgaris* |
| CE3C88 | uncertain | uncertain | uncertain | T/T | T/T | *vulgaris* |
| CE3C89 | uncertain | uncertain | uncertain | T/T | T/T | *vulgaris* |
| CE3C90 | uncertain | uncertain | uncertain | T/T | T/T | *vulgaris* |
| CE3C91 | uncertain | uncertain | uncertain | T/T | T/T | *vulgaris* |
| CE3C94 | uncertain | uncertain | uncertain | T/T | T/T | *vulgaris* |
| CE3C95 | uncertain | uncertain | uncertain | T/T | T/T | *vulgaris* |
| CE3C96 | uncertain | uncertain | uncertain | T/T | T/T | *vulgaris* |
| CE3C97 | uncertain | uncertain | uncertain | T/T | T/T | *vulgaris* |
| CE3C98 | uncertain | uncertain | uncertain | T/T | T/T | *vulgaris* |
| FSRC_01 | uncertain | uncertain | uncertain | T/T | T/T | *vulgaris* |
| FSRC_02 | uncertain | uncertain | uncertain | T/T | T/T | *vulgaris* |
| FSRC_04 | uncertain | uncertain | uncertain | T/T | T/T | *vulgaris* |
| FSRC_05 | uncertain | uncertain | uncertain | T/T | T/T | *vulgaris* |
| FSRC_06 | uncertain | uncertain | uncertain | T/T | T/T | *vulgaris* |
| MRI-KZ080818 | uncertain | ragged | *vulgaris* | A/T | G/T | n/a |
| MRI-KZ100415 | uncertain | ragged | *vulgaris* | T/T | T/T | n/a |
| MRI-KZ251 | uncertain | straight | *nivalis* | A/A | G/G | n/a |
| MRI-KZ257 | uncertain | ragged | *vulgaris* | T/T | T/T | n/a |
| MRI-KZ267 | uncertain | straight | *nivalis* | A/A | G/G | n/a |
| MRI-KZ279 | uncertain | straight | *nivalis* | A/A | G/G | n/a |
| MRI-KZ284 | uncertain | straight | *nivalis* | A/A | G/G | n/a |

| | | | | | | |
|---|---|---|---|---|---|---|
| MRI-KZ285 | uncertain | straight | *nivalis* | A/A | G/G | n/a |
| MRI-KZ286 | uncertain | ragged | *vulgaris* | A/T | G/T | n/a |
| MRI-KZ287 | uncertain | ragged | *vulgaris* | T/T | T/T | n/a |
| MRI-KZ288 | uncertain | ragged | *vulgaris* | T/T | T/T | n/a |
| MRI-KZ292 | uncertain | straight | *nivalis* | A/A | G/G | n/a |
| MRI-KZ295 | uncertain | straight | *nivalis* | A/A | G/G | n/a |
| MRI-KZ298 | uncertain | straight | *nivalis* | A/A | G/G | n/a |
| MRI-KZ306 | uncertain | ragged | *vulgaris* | A/T | G/T | n/a |
| MRI-KZ308 | uncertain | straight | *nivalis* | A/A | G/G | n/a |
| MRI-KZ314 | uncertain | ragged | *vulgaris* | A/T | G/T | n/a |
| MRI-KZ316 | uncertain | straight | *nivalis* | A/A | G/G | n/a |
| MRI-KZ318 | uncertain | ragged | *vulgaris* | T/T | T/T | n/a |
| MRI-KZ320 | uncertain | ragged | *vulgaris* | A/T | G/T | n/a |
| MRI-KZ321 | uncertain | straight | *nivalis* | A/A | G/G | n/a |
| MRI-KZ322 | uncertain | straight | *nivalis* | A/A | G/G | n/a |
| MRI-KZ324 | uncertain | straight | *nivalis* | A/A | G/G | n/a |
| MRI-KZ326 | uncertain | straight | *nivalis* | A/A | G/G | n/a |
| MRI-KZ366 | uncertain | ragged | *vulgaris* | A/T | G/T | n/a |
| MRI-KZ367 | uncertain | ragged | *vulgaris* | A/T | G/T | n/a |
| MRI-KZ376 | uncertain | ragged | *vulgaris* | A/T | G/T | n/a |
| MRI-KZ385 | uncertain | ragged | *vulgaris* | A/T | G/T | n/a |
| MRI-KZ388 | uncertain | ragged | *vulgaris* | A/T | G/T | n/a |
| MRI-KZ458 | uncertain | straight | *nivalis* | A/A | G/G | n/a |
| MRI103650 | uncertain | straight | *nivalis* | A/A | G/G | n/a |
| MRI104349 | white | uncertain | *nivalis* | A/A | G/G | n/a |
| MRI104350 | white | uncertain | *nivalis* | A/A | G/G | n/a |
| MRI108522 | white | uncertain | *nivalis* | A/A | G/G | n/a |
| MRI152404 | molting | uncertain | *nivalis* | A/A | G/G | n/a |
| MRI155673 / MRI-KZ201 | uncertain | straight | *nivalis* | A/A | G/G | n/a |
| MRI161184 | uncertain | straight | *nivalis* | A/A | G/G | n/a |
| MRI19672 / MRI-KZ126 | uncertain | straight | *nivalis* | A/A | G/G | n/a |
| MRI27063 | uncertain | ragged | *vulgaris* | T/T | T/T | n/a |
| MRI30257/1657 | uncertain | straight | *nivalis* | A/A | G/G | n/a |
| MRI30435/1835 | uncertain | straight | *nivalis* | A/A | G/G | n/a |
| MRI34457 | white | uncertain | *nivalis* | A/A | G/G | n/a |
| MRI34532 | molting | uncertain | *nivalis* | A/A | G/G | n/a |
| MRI36183 | uncertain | straight | *nivalis* | A/A | G/G | n/a |
| MRI36395 | uncertain | straight | *nivalis* | A/A | G/G | n/a |
| MRI37429/669 | uncertain | ragged | *vulgaris* | T/T | T/T | n/a |
| MRI41492 | uncertain | ragged | *vulgaris* | A/T | G/T | n/a |
| MRI42030 | white | uncertain | *nivalis* | A/A | G/G | n/a |
| MRI45143 | uncertain | ragged | *vulgaris* | T/T | T/T | n/a |
| MRI481/4726 | uncertain | ragged | *vulgaris* | T/T | T/T | n/a |
| MRI48469 | uncertain | ragged | *vulgaris* | T/T | T/T | n/a |

| | | | | | | |
|---|---|---|---|---|---|---|
| MRI51145 | white | uncertain | *nivalis* | A/A | G/G | n/a |
| MRI51148 | white | uncertain | *nivalis* | A/A | G/G | n/a |
| MRI51152 | white | uncertain | *nivalis* | A/A | G/G | n/a |
| MRI59832/10322 | uncertain | ragged | *vulgaris* | T/T | T/T | n/a |
| MRI61813/16427 | uncertain | ragged | *vulgaris* | T/T | T/T | n/a |
| MRI66797 | molting | uncertain | *nivalis* | A/A | G/G | n/a |
| MRI69800/5856 | uncertain | ragged | *vulgaris* | T/T | T/T | n/a |
| MRI72621 | uncertain | straight | *nivalis* | A/A | G/G | n/a |
| MRI7337 | white | uncertain | *nivalis* | A/A | G/G | n/a |
| MRI78813/22636 | uncertain | ragged | *vulgaris* | T/T | T/T | n/a |
| MRI87940 | white | uncertain | *nivalis* | A/A | G/G | n/a |
| NRM-MA20005001 | uncertain | uncertain | uncertain | T/T | T/T | *vulgaris* |
| NRM-MA20045270 | uncertain | uncertain | uncertain | T/T | T/T | *vulgaris* |
| NRM-MA20065154 | uncertain | uncertain | uncertain | A/T | G/T | *vulgaris* |
| NRM-MA20075291 | uncertain | uncertain | uncertain | A/A | G/G | *nivalis* |
| NRM-MA20075325 | uncertain | uncertain | uncertain | T/T | T/T | *vulgaris* |
| NRM-MA20075403 | uncertain | uncertain | uncertain | A/A | G/G | *nivalis* |
| NRM-MA20095145 | uncertain | uncertain | uncertain | A/A | G/G | *nivalis* |
| NRM-MA20115255 | uncertain | uncertain | uncertain | A/A | G/G | *nivalis* |
| NRM-MA557916 | white | uncertain | *nivalis* | A/A | G/G | n/a |
| NRM-MA557922 | white | uncertain | *nivalis* | A/A | G/G | n/a |
| NRM-MA557940 | white | uncertain | *nivalis* | A/A | G/G | n/a |
| NRM-MA557950 | white | uncertain | *nivalis* | A/A | G/G | n/a |
| NRM-MA557953 | white | uncertain | *nivalis* | A/A | G/G | n/a |
| NRM-MA557970 | white | uncertain | *nivalis* | A/A | G/G | n/a |
| NRM-MA557971 | white | uncertain | *nivalis* | A/A | G/G | n/a |
| NRM-MA557975 | white | uncertain | *nivalis* | A/A | G/G | n/a |
| NRM-MA557977 | white | uncertain | *nivalis* | A/A | G/G | n/a |
| NRM-MA557982 | white | uncertain | *nivalis* | A/A | G/G | n/a |
| NRM-MA557983 | white | uncertain | *nivalis* | A/A | G/G | n/a |
| NRM-MA557985 | white | uncertain | *nivalis* | A/A | G/G | n/a |
| NRM-MA557988 | white | uncertain | *nivalis* | A/A | G/G | n/a |
| NRM-MA557989 | white | uncertain | *nivalis* | A/A | G/G | n/a |
| NRM-MA557990 | white | uncertain | *nivalis* | A/A | G/G | n/a |
| NRM-MA557996 | white | uncertain | *nivalis* | A/A | G/G | n/a |
| NRM-MA557997 | white | uncertain | *nivalis* | A/A | G/G | n/a |
| NRM-MA557998 | white | uncertain | *nivalis* | A/A | G/G | n/a |
| NRM-MA557999 | white | uncertain | *nivalis* | A/A | G/G | n/a |
| NRM-MA558000 | white | uncertain | *nivalis* | A/A | G/G | n/a |
| NRM-MA558001 | white | uncertain | *nivalis* | A/A | G/G | n/a |
| NRM-MA558002 | white | uncertain | *nivalis* | A/A | G/G | n/a |
| NRM-MA558003 | white | uncertain | *nivalis* | A/A | G/G | n/a |
| NRM-MA558004 | white | uncertain | *nivalis* | A/A | G/G | n/a |
| NRM-MA558006 | white | uncertain | *nivalis* | A/A | G/G | n/a |

| | | | | | | |
|---|---|---|---|---|---|---|
| NRM-MA558007 | white | uncertain | *nivalis* | A/A | G/G | n/a |
| NRM-MA558008 | white | uncertain | *nivalis* | A/A | G/G | n/a |
| NRM-MA558015 | white | uncertain | *nivalis* | A/A | G/G | n/a |
| NRM-MA558016 | white | uncertain | *nivalis* | A/A | G/G | n/a |
| NRM-MA558018 | white | uncertain | *nivalis* | A/A | G/G | n/a |
| NRM-MA558019 | white | uncertain | *nivalis* | A/A | G/G | n/a |
| NRM-MA558020 | white | uncertain | *nivalis* | A/A | G/G | n/a |
| NRM-MA558025 | white | uncertain | *nivalis* | A/A | G/G | n/a |
| NRM-MA558026 | white | uncertain | *nivalis* | A/A | G/G | n/a |
| NRM-MA558034 | white | uncertain | *nivalis* | A/A | G/G | n/a |
| NRM-MA558035 | molting | uncertain | *nivalis* | A/A | G/G | n/a |
| NRM-MA558036 | white | uncertain | *nivalis* | A/A | G/G | n/a |
| NRM-MA558037 | white | uncertain | *nivalis* | A/A | G/G | n/a |
| NRM-MA558038 | molting | uncertain | *nivalis* | A/A | G/G | n/a |
| NRM-MA558039 | white | uncertain | *nivalis* | A/A | G/G | n/a |
| NRM-MA558040 | molting | uncertain | *nivalis* | A/A | G/G | n/a |
| NRM-MA558050 | brown | ragged | *vulgaris* | T/T | T/T | n/a |
| NRM-MA558051 | brown | ragged | *vulgaris* | T/T | T/T | n/a |
| NRM-MA558052 | brown | ragged | *vulgaris* | A/T | G/T | n/a |
| NRM-MA558053 | uncertain | ragged | *vulgaris* | T/T | T/T | n/a |
| NRM-MA558054 | uncertain | ragged | *vulgaris* | T/T | T/T | n/a |
| NRM-MA558055 | uncertain | ragged | *vulgaris* | T/T | T/T | n/a |
| NRM-MA558056 | uncertain | ragged | *vulgaris* | T/T | T/T | n/a |
| NRM-MA558057 | uncertain | ragged | *vulgaris* | T/T | T/T | n/a |
| NRM-MA558075 | brown | ragged | *vulgaris* | T/T | T/T | n/a |
| NRM-MA558085 | brown | ragged | *vulgaris* | T/T | T/T | n/a |
| NRM-MA558087 | brown | ragged | *vulgaris* | T/T | T/T | n/a |
| NRM-MA558090 | brown | ragged | *vulgaris* | T/T | T/T | n/a |
| NRM-MA558091 | brown | ragged | *vulgaris* | T/T | T/T | n/a |
| NRM-MA558094 | brown | ragged | *vulgaris* | T/T | T/T | n/a |
| NRM-MA558095 | brown | ragged | *vulgaris* | T/T | T/T | n/a |
| NRM-MA558098 | brown | ragged | *vulgaris* | T/T | T/T | n/a |
| NRM-MA558099 | brown | ragged | *vulgaris* | T/T | T/T | n/a |
| NRM-MA558115 | brown | ragged | *vulgaris* | T/T | T/T | n/a |
| NRM-MA558117 | brown | ragged | *vulgaris* | T/T | T/T | n/a |
| NRM-MA558120 | brown | ragged | *vulgaris* | T/T | T/T | n/a |
| NRM-MA558121 | brown | ragged | *vulgaris* | T/T | T/T | n/a |
| NRM-MA558124 | brown | ragged | *vulgaris* | T/T | T/T | n/a |
| NRM-MA558127 | brown | ragged | *vulgaris* | A/T | G/T | n/a |
| NRM-MA558132 | brown | ragged | *vulgaris* | T/T | T/T | n/a |
| NRM-MA558134 | brown | ragged | *vulgaris* | T/T | T/T | n/a |
| NRM-MA558142 | brown | ragged | *vulgaris* | T/T | T/T | n/a |
| NRM-MA558143 | brown | ragged | *vulgaris* | T/T | T/T | n/a |
| NRM-MA558146 | brown | ragged | *vulgaris* | T/T | T/T | n/a |

| | | | | | | |
|---|---|---|---|---|---|---|
| NRM-MA558148 | brown | ragged | *vulgaris* | A/T | G/T | n/a |
| NRM-MA558150 | white | uncertain | *nivalis* | A/A | G/G | n/a |
| NRM-MA558152 | white | uncertain | *nivalis* | A/A | G/G | n/a |
| NRM-MA558153 | white | uncertain | *nivalis* | A/A | G/G | n/a |
| NRM-MA558160 | brown | ragged | *vulgaris* | A/T | G/T | n/a |
| NRM-MA558162 | brown | ragged | *vulgaris* | A/T | G/T | n/a |
| NRM-MA558163 | brown | ragged | *vulgaris* | A/T | G/T | n/a |
| NRM-MA558164 | brown | ragged | *vulgaris* | T/T | T/T | n/a |
| NRM-MA558165 | brown | ragged | *vulgaris* | A/T | G/T | n/a |
| NRM-MA558167 | brown | ragged | *vulgaris* | A/T | G/T | n/a |
| NRM-MA558168 | brown | ragged | *vulgaris* | T/T | T/T | n/a |
| NRM-MA558172 | brown | ragged | *vulgaris* | T/T | T/T | n/a |
| NRM-MA558175 | white | uncertain | *nivalis* | A/A | G/G | n/a |
| NRM-MA558178 | white | uncertain | *nivalis* | A/A | G/G | n/a |
| NRM-MA558179 | white | uncertain | *nivalis* | A/A | G/G | n/a |
| NRM-MA558180 | white | uncertain | *nivalis* | A/A | G/G | n/a |
| NRM-MA558182 | brown | ragged | *vulgaris* | A/T | G/T | n/a |
| NRM-MA558183 | brown | ragged | *vulgaris* | A/T | G/T | n/a |
| NRM-MA558185 | brown | ragged | *vulgaris* | T/T | T/T | n/a |
| NRM-MA558188 | uncertain | ragged | *vulgaris* | T/T | T/T | n/a |
| NRM-MA600136 | brown | ragged | *vulgaris* | T/T | T/T | n/a |
| NRM-MA600137 | molting | uncertain | *nivalis* | A/A | G/G | n/a |
| NRM-MA600142 | brown | uncertain | *vulgaris* | T/T | T/T | n/a |
| NRM-MA600144 | molting | uncertain | *nivalis* | A/A | G/G | n/a |
| NRM-MA611919 | brown | ragged | *vulgaris* | T/T | T/T | n/a |
| SMNG-M.5101 | uncertain | uncertain | uncertain | T/T | T/T | *vulgaris* |
| SMNG-M.5629 | uncertain | uncertain | uncertain | T/T | T/T | *vulgaris* |
| SMNG-M.5628 | uncertain | uncertain | uncertain | T/T | T/T | *vulgaris* |
| SMNG-M.5627 | uncertain | uncertain | uncertain | T/T | T/T | *vulgaris* |
| ZFMK-MAM-1939.0149 | uncertain | ragged | *vulgaris* | T/T | T/T | n/a |
| ZFMK-MAM-1983.0276 | uncertain | ragged | *vulgaris* | T/T | T/T | n/a |
| ZFMK-MAM-2007.0274 | brown | ragged | *vulgaris* | T/T | T/T | n/a |
| ZFMK-MAM-2007.0275 | brown | ragged | *vulgaris* | T/T | T/T | n/a |
| ZFMK-MAM-2008.0055 | uncertain | ragged | *vulgaris* | T/T | T/T | n/a |
| ZFMK-MAM-2008.0059 | uncertain | ragged | *vulgaris* | T/T | T/T | n/a |

[1] Museum/Research Collections:
AMNH - American Museum of Natural History
BNM - Bündner Naturmuseum
CE3C - Centre for Ecology, Evolution and Environmental Changes
FSRC - Franz Suchentrunk research collection (private collection)
MRI - Mammal Research Institute of the Polish Academy of Sciences
NRM - Swedish Museum of Natural History
SMNG - Senckenberg Museum of Natural History Goerlitz
ZFMK - Zoological Research Museum Alexander Koenig

**Table S7 - Demographic inference parameter point estimates and confidence intervals, estimated with GADMA.** Estimates for a two-population demographic model including North and Central ancestry lineages (see Fig. 3E) are shown, including effective population sizes ($N_e$), time of divergence (T), and migration rates (m). Parameters were scaled assuming a mutation rate of 2.2 x 10e-9 (ref.[16]) and a generation time of one year (ref.[17]).

| Parameter | Point estimate (best model) | 95% Confidence Intervals |
|---|---|---|
| $N_e$ (Ancestral) | 693 107 | 669 828 – 716 387 |
| $N_e$ (North, after split) | 1 838 146 | 1 642 910 – 2 033 382 |
| $N_e$ (North, current) | 367 895 | 349 140 – 386 651 |
| $N_e$ (Central, after split) | 6 931 | 6 588 – 7 274 |
| $N_e$ (Central, current) | 2 026 672 | 1 916 816 – 2 136 527 |
| T (North-Central) | 1 007 481 | 935 646 – 1 079 315 |
| m (North → Central) | 1.77 x 10e-6 | 1.68 x 10e-6 – 1.85 x 10e-6 |
| m (Central → North) | 3.19 x 10e-7 | 2.72 x 10e-7 – 3.66 x 10e-7 |

**Table S8 - Estimated *MC1R* haplotype divergence times across *Mustela* and *Neogale* species.** Haplotype divergence was estimated using the Bayesian coalescent framework implemented in BEAST. Node codes are those of Supplementary Fig. S13. For each node, the divergence time point estimate, and the 95% highest posterior density (95% HPD) are shown in million years (Mya).

| Node code | Divergence estimate (Mya) | 95% HPD |
|---|---|---|
| a | 0.05 | 0.02 – 0.09 |
| b | 0.18 | 0.12 – 0.26 |
| c | 0.26 | 0.17 – 0.37 |
| d | 0.95 | 0.63 – 1.26 |
| e | 0.24 | 0.15 – 0.33 |
| f | 0.35 | 0.23 – 0.48 |
| g | 0.07 | 0.03 – 0.12 |
| h | 2.57 | 1.81 – 3.39 |
| i | 1.62 | 1.13 – 2.14 |
| j | 0.26 | 0.17 – 0.38 |
| k | 1.04 | 0.71 – 1.38 |
| l | 1.33 | 0.91 – 1.74 |
| m | 4.91 | 3.51 – 6.51 |
| n | 6.62 | 4.74 – 8.74 |
| o | 3.08 | 2.10 – 4.02 |
| p | 9.43 | 6.57 – 12.45 |
| q | 10.61 | 7.56 – 14.00 |

**Table S9 - Probability of occurrence of alternative winter phenotypes in the inferred North and Central genetic lineages.** Probabilities (P) were averaged based on the phenotype probability models of ref.[14], considering a 200 km buffer around the sampling locations of individuals attributed to each genetic lineage.

| | Lineage | |
|---|---|---|
| **P** | **North** | **Central** |
| winter-brown | 0.253 | 0.768 |
| winter-white | 0.747 | 0.232 |

**Data S1 (separate file) - Least weasel (*Mustela nivalis*) specimens used in this study and associated metadata.** Sample codes refer to the collection numbers of the museum or research collection of origin. Information on coloration morph (for European samples), dorsoventral line and winter color (for European and North American samples), collection date, country, and locality are given, when available. If the specimen was not visually observed during the development of this work, winter color, dorsoventral line, and coloration morph were deemed "uncertain". Samples were included in analyses set 1 to 3 based on this classification. DNA sources and information about generated data (capture experiments and phenotyping data) are also provided.

**Data S2 (separate file) - Sequencing and capture statistics for least weasel (*Mustela nivalis*) capture data.** For each sample, information is given on the number of independent capture pools where the sample was included, the number of raw paired-end reads, the mapping percentage of sequencing data, and capture success. For capture success, statistics include the percentage of bases off-bait, mean target coverage, and percentage of target bases at 10X coverage (for all capture targets, *MC1R* region, 2 kb fragments, and SNP fragments). Inclusion in different analysis datasets is indicated per sample.

**Data S3 (separate file) - Mustelidae specimens used in this study and associated metadata. (A)** Samples from museum collections, with newly generated capture data. Sample codes refer to the museum's collection numbers. Information on collection date, country, and locality are given, when available. Winter color is indicated, when applicable. **(B)** Samples from publicly available datasets, with whole-genome resequencing data. Sample codes refer to the original dataset in which sequencing was generated. Information on collection data, country, and locality are given, when available. Original data sources are indicated, including accession number and database (NCBI or ENA).

**Data S4 (separate file) - Sequencing and capture statistics for other Mustelidae species capture data. (A)** Samples from museum collections, with newly generated capture data. For each sample, information is given on the number of independent capture pools where the sample was included, the number of raw paired-end reads, the mapping percentage of sequencing data, and capture success. For capture success, statistics include the percentage of bases off-bait, mean target coverage, and percentage of target bases at 10X coverage (for all capture targets, *MC1R* region, 2 kb fragments, and SNP fragments). **(B)** Samples from publicly available datasets, with whole-genome resequencing data. For each sample, details on the number of raw reads, mapping percentage against *M. nivalis* reference genome, and genome coverage are given. Coverage details for the capture regions are also provided. Inclusion in different analysis datasets is indicated per sample.
